# EndoMAP.v1, a Structural Protein Complex Landscape of Human Endosomes

**DOI:** 10.1101/2025.02.07.636106

**Authors:** Miguel A. Gonzalez-Lozano, Ernst W. Schmid, Enya Miguel Whelan, Yizhi Jiang, Joao A. Paulo, Johannes C. Walter, J. Wade Harper

## Abstract

**Early/sorting endosomes are dynamic organelles that play key roles in proteome control by triaging plasma membrane proteins for either recycling or degradation in the lysosome^1,2,3^. These events are coordinated by numerous transiently-associated regulatory complexes and integral membrane components that contribute to organelle identity during endosome maturation^4^. While a subset of the several hundred protein components and cargoes known to associate with endosomes have been studied at the biochemical and/or structural level, interaction partners and higher order molecular assemblies for many endosomal components remain unknown. Here, we combine cross-linking and native gel mass spectrometry^5-8^ of purified early endosomes with AlphaFold^9,10^ and computational analysis to create a systematic human endosomal structural interactome. We present dozens of structural models for endosomal protein pairs and higher order assemblies supported by experimental cross-links from their native subcellular context, suggesting structural mechanisms for previously reported regulatory processes. Using induced neurons, we validate two candidate complexes whose interactions are supported by crosslinks and structural predictions: TMEM230 as a subunit of ATP8/11 lipid flippases^11^ and TMEM9/9B as subunits of CLCN3/4/5 chloride-proton antiporters^12^. This resource and its accompanying structural network viewer provide an experimental framework for understanding organellar structural interactomes and large-scale validation of structural predictions.**

## INTRODUCTION

Plasma membrane (PM) protein flux within cells is controlled, in part, through a series of membrane-bound organelles referred to as the endolysosomal system^2^. Endosomes are born on the cytosolic face of the PM through clathrin-dependent and independent vesicle budding processes, and rapidly undergo conversion to RAB5-positive vesicles referred to as early/sorting endosomes or, for simplicity, as early endosomes^3^. Early endosomes receive regulatory proteins by fusion with Golgi-derived transport vesicles and additional cargo by fusion with other early endosomes^13^. Early endosomes serve as platforms for protein sorting, which includes the formation of recycling vesicles for cargo transport back to the PM or Golgi. Moreover, in a dynamic process, RAB5-positive endosomes mature to RAB7-positive endosomes, referred to as late endosomes, where ESCRT-mediated delivery of primarily PM protein cargo to intralumenal vesicles facilitates cargo degradation upon late endosome maturation into lysosomes^2,4,14^. Fully degradative lysosomes also play key roles in the elimination of intracellular proteins and organelles through autophagy, wherein cargo-laden autophagosomes fuse with the lysosome to facilitate cargo degradation by lumenal hydrolases^15^.

Our understanding of the endolysosomal system has been advanced through the identification of several functional modules that participate in core endosomal functions, including vesicle fusion, cargo trafficking, and organelle maturation^1-4,14,16,17,18,19^. Moreover, recent developments in purification methods have facilitated the identification of proteins that spend all or part of their lifecycle in the endosomal system, including several proteins linked with neurodegenerative and lysosomal storage diseases^20-23^. However, the dynamic nature of these organelles has made some protein assignments controversial^24,25^. Much of the biology of endosomes occurs on the membrane surface and often involves transient interactions between integral membrane and peripheral proteins whose association may be lost during co-immunoprecipitation and other protein isolation methods. Consequently, gaps exist in our understanding of the proteins, structures, and interactions across the endosomal system.

Here, we present “EndoMAP.v1”, a structural protein interactome of human early endosomes. We focused on an endosomal sub-population characterized by association with Early Endosome Antigen 1 (EEA1), used here as an organelle isolation handle via Endo-IP^25^. EndoMAP.v1 combines cross-linking-mass spectrometry (XL-MS)^5-7,26^ and blue-native gel co-fractionation-MS (BN-MS) to generate a comprehensive network of protein interactions and candidate complexes in EEA1-associated endosomes (**Fig. 1a**). Large-scale AlphaFold Multimer (AF-M)^9,10^ and AlphaLink2^27^ analysis across the network generated hundreds of structural predictions supported by cross-link distance constraints, which are available via the EndoMAP.v1 structural interactome viewer (https://endomap.hms.harvard.edu/). We demonstrated the value of this resource: 1) through validation of transmembrane subunits of endosomal lipid flippases and chloride-proton (Cl^-^-H^+^) antiporters, and 2) through cross-link-informed structural predictions of dozens of protein interactions and multi-protein assemblies across diverse core endosomal functional categories. Together, EndoMAP.v1 provides a resource for mechanistic analysis of early endosome complexes and an experimental framework for understanding structural interactomes for specific organelles.

**Fig. 1.**
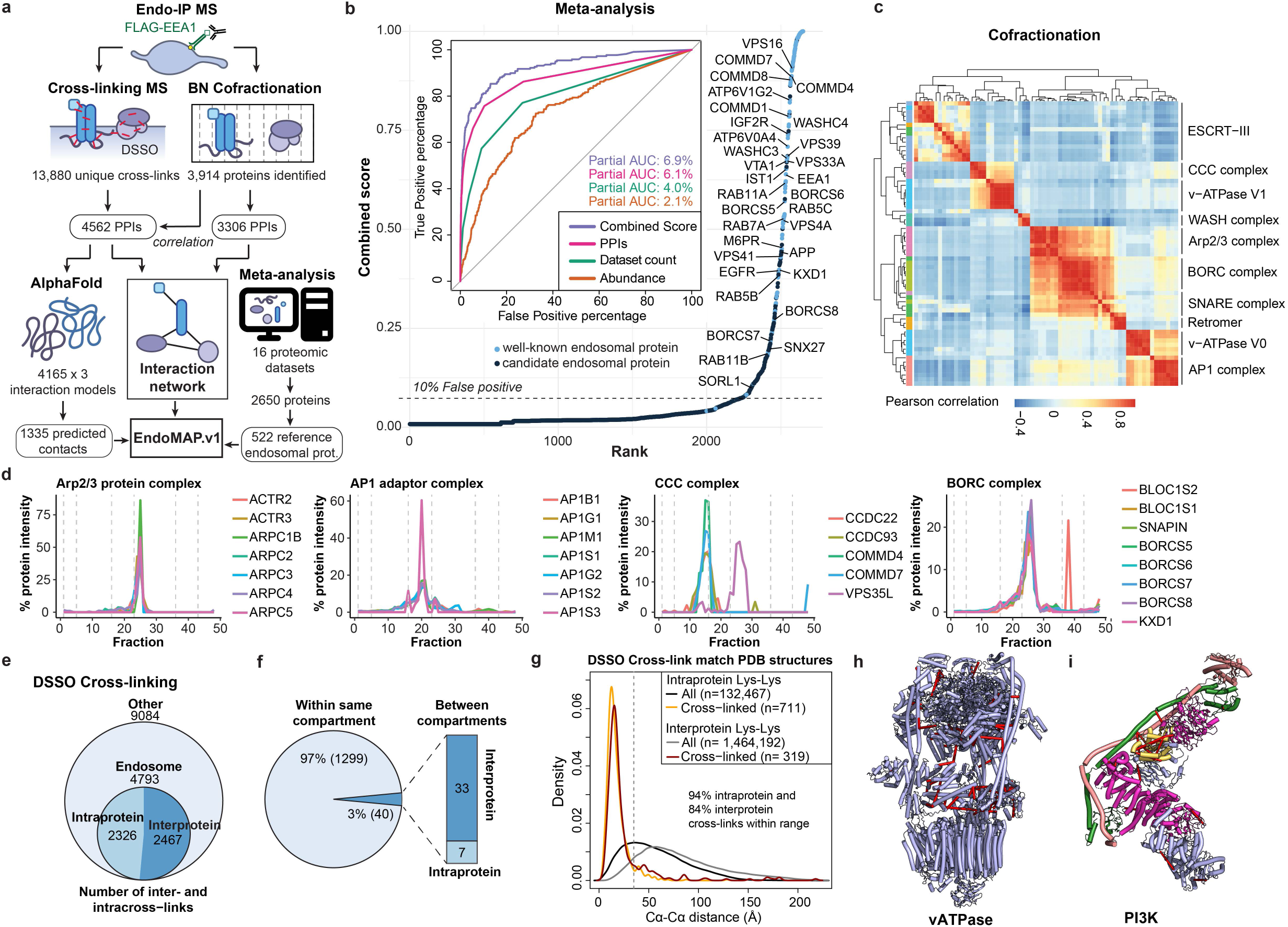
EEA1-positive Endosomal Proteome Analysis via Dual Complexomics Strategies. **a**, EndoMAP.v1 workflow schematic depicting integration of XL-MS, BN-MS, meta-analysis and structural predictions to create an endosomal protein complex structural interaction landscape. **b**, Known (blue) and candidate (black) endosomal proteins ranked based on meta-analysis combined score, with higher values indicating higher probability of a protein being endosomal. Inset shows receiver operating characteristic (ROC) curves for each individual metric and its combination for annotating endosomal proteins. Partial Area Under the Curve (AUC) is indicated at 10% false positive percentage. **c**, Correlation heatmap of BN-MS co-fractionation data showing unsupervised clustering of well-known endosomal complexes. **d**, Co-fractionation profiles of selected protein complexes from BN-MS. **e**, Summary of DSSO cross-links identified in Endo-IP samples, including intraprotein and interprotein cross-links involving high-confidence endosomal proteins. **f**, Pie chart showing the number of DSSO cross-links within and between topological compartments based on Uniprot. **g**, Density plots showing the distribution of Cα-Cα distances (Å) for intra and interprotein DSSO cross-links for all structures available in the Protein Data Bank (PDB) for the entire XL-MS dataset. The vertical dotted line indicates the maximum distance allowed by the cross-linker. **h,i**, Identified DSSO cross-links (red lines) mapped into the endolysosomal V-ATPase (panel **h**, PDB:6WM2)^66^ and the Class II PI3P lipid kinase complex (panel **i**, PDB:7BL1)^35^.

## RESULTS

### Early Endosome Interactome via Dual Complexomics Approaches

To understand protein interactions associated with EEA1-positive endosomes, we developed an experimental and informatic complexomics pipeline (**Fig. 1a**). Given the dynamic nature of the endosomal system, we first wanted to define and characterize the endosomal proteome, including resident and transient endosomal proteins such as cargo, based on published experimental data. We analyzed the proteins identified in 16 studies that involved a variety of purification approaches and cell types (**Extended Data Fig. 1a, Supplementary Table 1**). Multiple Correspondence Analysis (MCA) revealed extensive diversity across the datasets that segregated, in part, based on purification method (**Extended Data Fig. 1b**). Endo-IP^25^ and correlation-based gradient fractionation^24^ approaches recovered the largest number of well-known endosomal proteins (111 and 107, respectively; **Extended Data Fig. 1a**, see **METHODS**). Nevertheless, the presence of individual proteins in multiple datasets was predictive of endosomal localization, including proteins transiently localized to endosomes throughout the dynamics of endosomal function and maturation to lysosomes (**Extended Data Fig. 1c**). In addition, the presence of specific endosomal proteins in multiple datasets correlated with higher protein abundance in samples purified by Endo-IP^25^ (**Extended Data Fig. 1d**). These metrics were compared as predictors of endosomal localization, including: **1)** the number of independent datasets in which each protein was identified, **2)** the protein abundance in Endo-IP, and **3)** the number of interactions with well-known endosomal proteins (**Fig. 1b**). Predictors were evaluated using a manually curated set of well-known endosomal proteins as reference (**Supplementary Table 1,** see **METHODS**). The combination of all three predictors best captured many well-characterized endosomal proteins with high confidence (**Fig. 1b**). Moreover, this combined score identified numerous predicted endosomal proteins that appear to be understudied based on large-scale protein interaction studies (BioPlex and OpenCell)^28,29^ and published literature (**Extended Data Fig. 1e-g**). In total, this analysis identified 522 endosomal proteins (**Supplementary Table 1**), including known and predicted endosomal proteins that were the target for further characterization with our complexomics pipeline as “reference” endosomal proteome (**Fig. 1a**).

We then employed BN-MS and cross-linking by XL-MS to identify candidate protein-protein interactions (PPIs) (**Fig. 1a**). We further optimized and extensively evaluated the Endo-IP approach in HEK293 cells^25^, with early endosomes eluted from the affinity matrix under detergent-free conditions for XL-MS or using detergent for BN-MS (**Extended Data Fig. 1h-l**, see **METHODS**). Triplicate Endo-IP samples were fractionated by BN gel electrophoresis, and 48 individual fractions across all mass ranges were subjected to MS analysis, identifying 3914 unique proteins (**Supplementary Table 2**). Numerous well characterized endosomal protein complexes were found to co-fractionate based on Pearson coefficients of normalized elution profiles (**Fig. 1c**). These include the <u>B</u>LOC-<u>o</u>ne <u>R</u>elated <u>C</u>omplex (BORC) involved in endolysosomal positioning^30^, components of the <u>Ho</u>motypic Fusion and <u>P</u>rotein <u>S</u>orting (HOPS) complex^31^, and the AP1 adaptor complex that traffics cargo to endolysosomes^32^, among others (**Fig. 1c,d; Extended Data Fig. 2a**). Unbiased correlation profiling using PCProphet^33^ revealed the presence of 3,306 candidate interacting proteins pairs. To recover high confidence candidate interactions, we only considered interactions with a score of at least 0.7 in two replicates, which maximized the recovery of interactions reported in Bioplex (**Extended Data Fig. 2b,c, Supplementary Table 2,** see **METHODS**).

In parallel, duplicate matrix- and detergent-free Endo-IP samples were cross-linked using the MS-cleavable DSSO Lys-Lys cross-linker and analyzed by XL-MS to identify proximal protein pairs in intact organelles^5,7^ (**Fig. 1a**). We identified 13,877 unique DSSO cross-links, of which 4,793 involved intra or inter-protein cross-links among our reference endosomal proteins (inclusive of the EEA1 endosomal purification handle) (**Fig. 1e, Supplementary Table 2**). Ninety-seven % of the cross-links matched the expected topological connectivity (within cytosolic, luminal or extracellular regions), consistent with the purification of intact organelles with Endo-IP (**Fig. 1f**). This is within the range of the 5% false-discovery rate (FDR) employed for cross-link identification. To evaluate the quality and specificity of cross-linking across the full dataset (including all non-endosomal proteins), we compared 1,030 cross-linked Lys(Cα)- Lys(Cα) distances for all available <u>P</u>rotein <u>D</u>ata <u>Bank</u> (PDB) structures (219 in total). The vast majority of intraprotein (94%) and interprotein (84%) cross-links were within the 35Å maximum distance for DSSO cross-linker (considering in-solution flexibility)^34^ (**Fig. 1g**). Representative endosomal multiprotein complexes (V-ATPase and the Class II PI3-kinase, PIK3C3-BECN1-UVRAG-PIK3R4)^35^ are shown in **Fig 1h,i**, with multiple cross-links among proteins within each complex. Although there is mild bias toward more abundant proteins (**Extended Data Fig. 2d-e**), cross-links are detected across the complete span of protein copy number (**Extended Data Fig. 2f**). In terms of protein-protein interactions, proteins with a higher number of cross-link supported interactions were correlated with copy number and number of interactions in BioPlex, but not with molecular weight (**Extended Data Fig. 2g-i**), as previously observed^7^. Limited overlap between cross-linked pairs and interaction pairs reported in BioPlex^28^ or yeast two hybrid datasets^36^ is consistent with the maintenance of weaker interactions in the context of organelle cross-linking (**Extended Data Fig. 2j**). Importantly, interactions with higher numbers of cross-links have better co-elution SECAT (<u>S</u>ize-<u>E</u>xclusion <u>C</u>hromatography <u>A</u>lgorithmic <u>T</u>oolkit) p-values in BN (**Extended Data Fig. 2k, Supplementary Table 2**). Additionally, previously reported cross-linked interactions have a better co-elution SECAT p-value than novel candidate interactions, which most likely include interactions that are transient and difficult to identify by other methods (**Extended Data Fig. 2l,m**). Taken together, we identified a total of 4,562 and 3,306 protein interactions by cross-linking and BN-MS, respectively, which provide a useful dataset for exploration of early endosome protein interactions (**Extended Data Fig. 2n,o, Supplementary Table 2**).

### Integrated Early Endosome Interaction Landscape

To create an early endosome interaction map, we integrated XL-MS and BN-MS data with our reference endosomal proteome, applying stringent filters (see **METHODS**). The resulting network displayed an average shortest path distance of 6.2 and followed a Power Law distribution with R^2^>0.95 (**Extended Data Fig. 3a-c**). Exploring the connectivity between localization descriptors, we found that endosomal proteins were highly connected with other endosomal proteins or proteins annotated as lysosomal or Golgi (88% of the endosomal interactions), with significantly fewer connections with other organelles (i.e. mitochondria or nucleus) (**Extended Data Fig. 3d**, see **METHODS**). Additional filtering, including centering the network around our reference endosomal proteome and filtering of doubtful connectivity (i.e. nuclear proteins, which correspond to up to 8.5% of the interactions with endosomal proteins) (**Extended Data Fig. 3e**, see **METHODS**), yielded a network containing 1,933 nodes and 4,282 edges. The core component of the network (without disconnected modules) included 1,722 protein and 3,489 interactions organized in 41 communities, which were significantly enriched for several well-known endosomal complexes, including V-ATPase, SNAREs, and the CCC (CCDC22, CCDC93, COMMD) complex (**Fig. 2a, Supplementary Table 2**). Indeed, proteins belonging to the same known complex were significantly closer and in direct contact within the network (**Fig. 2b,c, Extended Data Fig. 3f**). Through an unbiased enrichment analysis of all disease pathways in DisGenNET, we found that Parkinson’s disease-related genes were the most highly enriched in our reference endosomal proteome (**Extended Data Fig. 3g-i, Supplementary Table 1**). Other neurodegenerative disorders were also enriched, including lysosomal storage disorder (LSD) proteins, many of which are actively trafficked to early endosomes^20,21^. Proteins linked with these disorders display the shortest path distance (∼5.0) reflective of their density within the network (**Extended Data Fig. 3j,k**). As elaborated below, this network provides a discovery platform for understanding the interaction landscape of early endosomes.

**Fig. 2.**
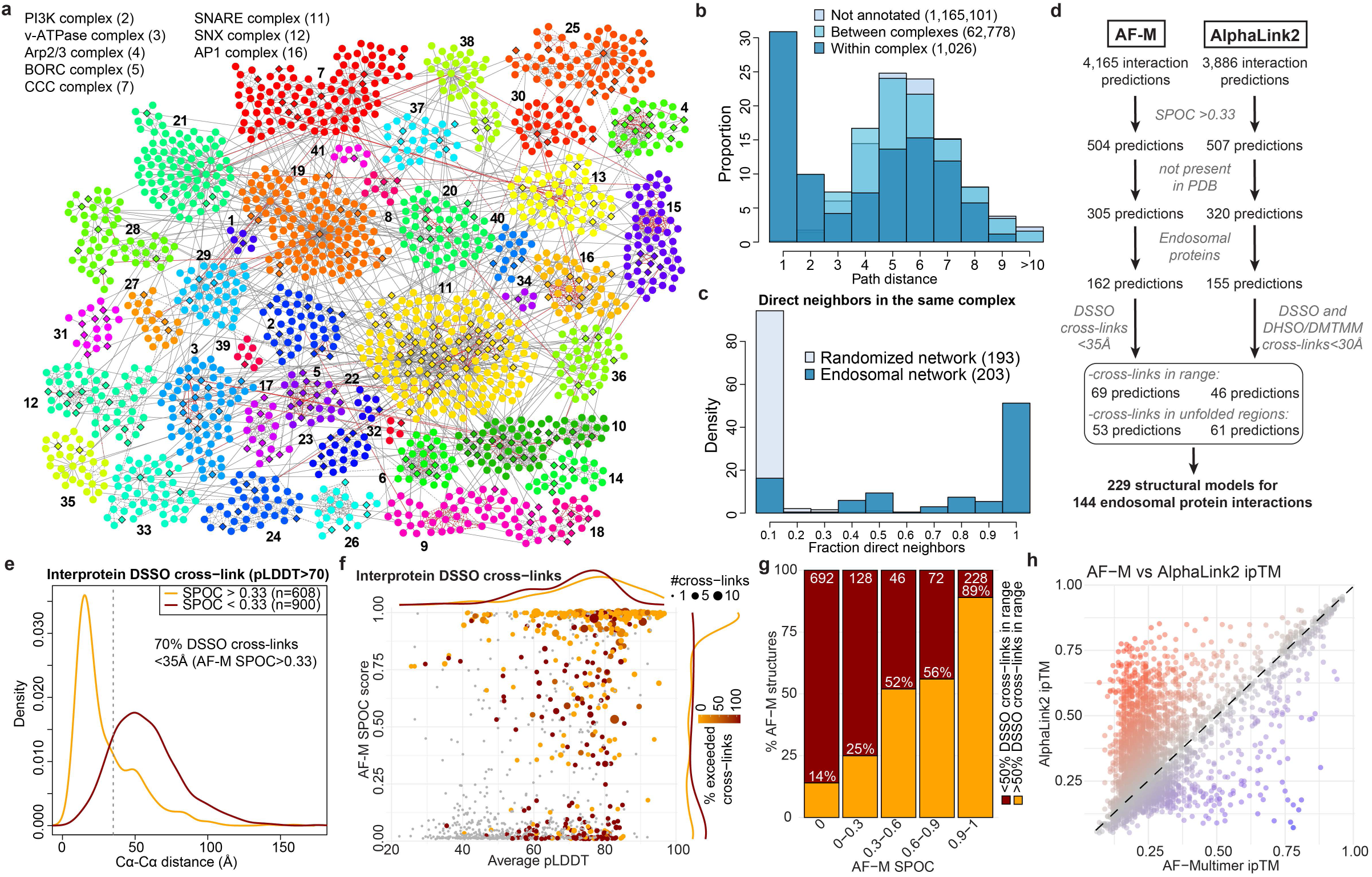
Assembly of an Integrated Endosome Protein Complex Structural Landscape. **a**, Core component of the network containing 1,722 nodes organized into 41 communities (indicated by numbers) and 3,489 edges. Significantly enriched protein complexes of selected communities are provided in the top left (see **Supplementary Table 2** for full list of communities). Diamonds and circular nodes represent high confidence endosomal and other proteins, respectively. Solid and dotted edges represent interactions identified by at least one cross-link or only co-fractionating, respectively. Red edges indicate interaction previously reported. **b**, Distribution of path distances between proteins within and between the same complex compared to proteins without complex annotation. **c**, Distribution of fraction of direct neighbors in the same complex for each protein compared to a randomized network control. **d**, Systematic AF-M and AlphaLink2 predictions of protein interactions identified by XL-MS and match with the cross-link distance constraints. **e**, Distribution of Cα-Cα distances (Å) for interprotein DSSO cross-links reflecting AF-M predictions with SPOC > 0.33 (orange) and SPOC < 0.33 (red). **f**, Distribution of AF-M SPOC scores and average pLDDT for predictions with SPOC > 0. Number of interprotein cross-links evaluated and exceeding the DSSO cross-linker distance constraints are indicated by point size and the color, respectively. **g**, Percentage of pairwise AF-M predictions with more or less than 50% of cross-links within the distance constrain (orange and red, respectively) relative to the SPOC score. **h**, ipTM scores for AF-M compared to AlphaLink2 predictions. Color gradient represents the score difference; higher in AlphaLink2 (red) or AF-M (blue).

### Large-scale AlphaFold Predictions of Cross-linked Proteins

To transform the endolysosomal network into a structurally informed interactome, we performed large-scale AF-M predictions^9,10^. We analyzed 4,165 protein pairs identified by XL-MS (total residue length <3600 amino acids due to computational constraints), including both endosomal and non-endosomal protein pairs. We ranked each pair using a <u>S</u>tructure <u>P</u>rediction and <u>O</u>mics-informed <u>C</u>lassifier (SPOC)^37^ to evaluate complex plausibility (**Supplementary Table 3**). SPOC considers ipTM and PAE scores of the predicted interface (among other metrics) together with biological correlations among the interacting proteins (such as colocalization and genetic co-dependency) and scores above 0.33 (scale 0-1) can indicate direct interactions^37^. We then independently assessed the reliability of the predictions by evaluating the extent to which structural predictions were consistent with DSSO cross-link distance constraints (**Fig. 2d, Supplementary Table 3**). As expected, there was a strong correlation between distances in AF-M predictions and the corresponding structures in the PDB, both for intra- and inter-protein cross-links (**Extended Data Fig. 3l**). Moreover, within all pairwise predictions, 93% and 38% of intraprotein and interprotein DSSO cross-link distances, respectively, were within range (<35Å)^34^ (**Extended Data Fig. 3m,n**). In the latter case, the bi-modal distribution was largely explained by protein pairs where AF-M was unable to predict an interaction (SPOC<0.33), since 70% of interprotein cross-link distances were within range for pairs with SPOC>0.33 (**Fig. 2e**). Importantly, the fraction of predictions with interprotein cross-links satisfying the length requirements correlated with the SPOC score (**Fig. 2f,g**). We also observed a correlation between the number of cross-links identified for an interaction and its prediction SPOC score **(Extended Data Fig. 3o,p**). Predictions involving at least one endosomal protein had a similar distribution of cross-link matches as predictions from the full dataset (**Extended Data Fig. 3q-s**). Therefore, SPOC scores and cross-linking data are complementary approaches that provide structural and experiment support to the interactions identified in EndoMAP.v1. With AF-M, we obtained 162 unique, endosomal pairwise structural predictions not present in the PDB with SPOC>0.33, including 69 structures matching interprotein cross-link constraints, 53 with cross-links in unstructured regions and 40 structures not matching cross-link constraints (**Fig. 2d**).

Three approaches were used to further strengthen and extend structural modeling within EndoMAP.v1. First, DSSO cross-linking data was evaluated using the recently reported Scout search engine^38^ (**Supplementary Table 2**). Scout with 1% FDR recovered 43% of those cross-links identified by XLinkX at 5% FDR, 66% between endosomal proteins (**Extended Data Fig. 4a**), including most examples described below. Regarding protein interactions, our pipeline filtering criteria substantially increased the overlap with Scout, with up to 79% overlap for the interactions between endosomal proteins with good AF-M predictions (SPOC>0.33) matching the DSSO cross-link distance restraint (**Extended Data Fig. 4b**). Nevertheless, Scout recovered only 61% of previously reported interactions identified using XlinkX, suggesting that there is still true connectivity that was missed by the more stringent Scout search (**Extended Data Fig. 4b**). All interactions found at 1%FDR are indicated in the web portal and **Supplementary Table 2**.

Second, we used AlphaLink2^27^ to generate structural predictions assisted by DSSO cross-link data and compare them to AF-M. We generated models for 3886 protein pairs identified by XL-MS (total residue length <3000 amino acids due to computational constraints) (**Supplementary Table 3**). Typically, predictions with strong scores showed comparable ipTM and SPOC for AF-M and AlphaLink2, while predictions with AF-M ipTM<0.3 showed frequently higher AlphaLink2 ipTM (**Fig. 2h**, **Extended Data Fig. 4c**). DSSO cross-link distances were comparable between AF-M and AlphaLink2 predictions, both for intra- and inter-protein cross-links (**Extended Data Fig. 4d**). For instance, ipTM scores for AF-M and AlphaLink2 were comparable for the interaction of ARL8B with the N-terminal RUN domain of RUFY2 (0.86 and 0.81, respectively), and the cross-link remained exceeding the acceptable distance (**Extended Data Fig. 4e**). Several examples illustrate cases of endosomal interactions with score or cross-link distance differences between AF-M and AlphaLink2 (**Extended Data Fig. 4f-i**). For VPS35-RAB7A, AlphaLink2 predicted a shorter DSSO cross-link distance (35.9Å versus 23.2Å), but with reduced ipTM (0.83 to 0.64) compared to AF-M, by most likely placing RAB7A on the incorrect surface of the VPS35 solenoid (**Extended Data Fig. 4f**). In contrast, AlphaLink2 showed higher scores for VAMP3 interaction with either SCAMP1 or SCAMP3 (**Extended Data Fig. 4h,i**). AlphaLink2 places the cytosolic helical domain of SCAMP3 in an orientation compatible with the membrane topology, the cross-link within the distance constraint (12.0Å), and a higher ipTM score (0.38) (**Extended Data Fig. 4i**).

Third, we performed an additional Endo-IP XL-MS experiment using alternative cross-linkers (DHSO and DMTMM) to evaluate AlphaLink2 and provide further evidence for AF-M structural predictions. DHSO and DMTMM can cross-link pairs of acidic residues or acid residues with Lys, respectively^6,39^. We identified 237 and 3084 cross-links with DHSO and DMTMM (1% FDR), respectively, which was in the expected range compared to DSSO^39^ (**Extended Data Fig. 4j, Supplementary Table 2**). Around 90% of the DHSO/DMTMM cross-links could be mapped to the same proteins and interactions identified with DSSO (69 and 623 interprotein interactions with DHSO and DMTMM, respectively), such as V-ATPase (**Extended Data Fig. 4k,l**). Within AlphaLink2, 88% and 87% of intraprotein DSSO and DMTMM cross-links distances, respectively, were within range (<30Å, **Extended Data Fig. 4m**, see **METHODS**). Only 51 DHSO cross-links could be mapped to structured regions (pLDDT>70) of AlphaLink2 predictions, all within the distance constraint. For interprotein cross-links, 65% and 39% of DSSO and DMTMM distances, respectively, were within range for pairs with SPOC>0.33 (**Extended Data Fig. 4n,o**). In summary, we obtained 155 endosomal predictions with AlphaLink2 (SPOC>0.33) that, together with AF-M, make 229 structural predictions for 144 endosomal interactions not present in the PDB that match interprotein cross-link constraints (or with cross-links in unstructured regions) (**Fig. 2d**). Together, we generated an experimentally supported structural interactome of the early endosomal system.

### TMEM230 as New Lipid Flippase Subunit

To validate interactions and structural models within EndoMAP.v1, we initially selected the TMEM230-ATP11B-TMEM30A complex given: 1) strong structural prediction scores for TMEM230-ATP11B (AF-M SPOC=0.64, ipTM=0.75) (**Fig. 3a**), 2) clear co-migration in BN-MS (**Fig. 3b**), and 3) a TMEM230-ATP11B cross-link satisfying distance constraints (**Fig. 3a, Supplementary Table 2**). ATP11 proteins (A, B and C) are P4-type ATP-dependent enzymes that flip lipids from exo to cytosolic leaflets of a bilayer^40^, mainly the endolysosomal membrane for ATP11B^11^. ATP11, as well as ATP8A1/2, interact with TMEM30A/B (also called CDC50A/B)^41^, required for flippase trafficking from ER to Golgi^42,43^. The AF-M TMEM230-ATP11B-TMEM30A heterotrimer prediction closely matched previously reported ATP11-TMEM30 structures^40,41^ and predicted packing of TMEM230’s transmembrane (TM) and N-terminal cytosolic segments with ATP11B TM1 and cytosolic catalytic domain, respectively (**Fig. 3a**). The AlphaLink2 prediction for TMEM230-ATP11B displayed a similar TMEM230-ATP11B interface with slightly longer cross-link distance as AF-M (**Fig. 3c**, **Extended Data Fig. 5a**). The ATPase domain of ATP11B in the predicted heterotrimer approximates the EP2 conformation of the corresponding orthologous yeast DNF2 protein (**Extended Data Fig. 5b**). ATP11B interaction with TMEM230/TMEM30A was confirmed reciprocally via co-immunoprecipitation (co-IP) in HEK293 cells (**Fig. 3d, Extended Data Fig. 5c**)^28^. This data identified TMEM230 as a subunit of the ATP11 family of lipid flippases and provided structural predictions of the complexes.

**Fig. 3.**
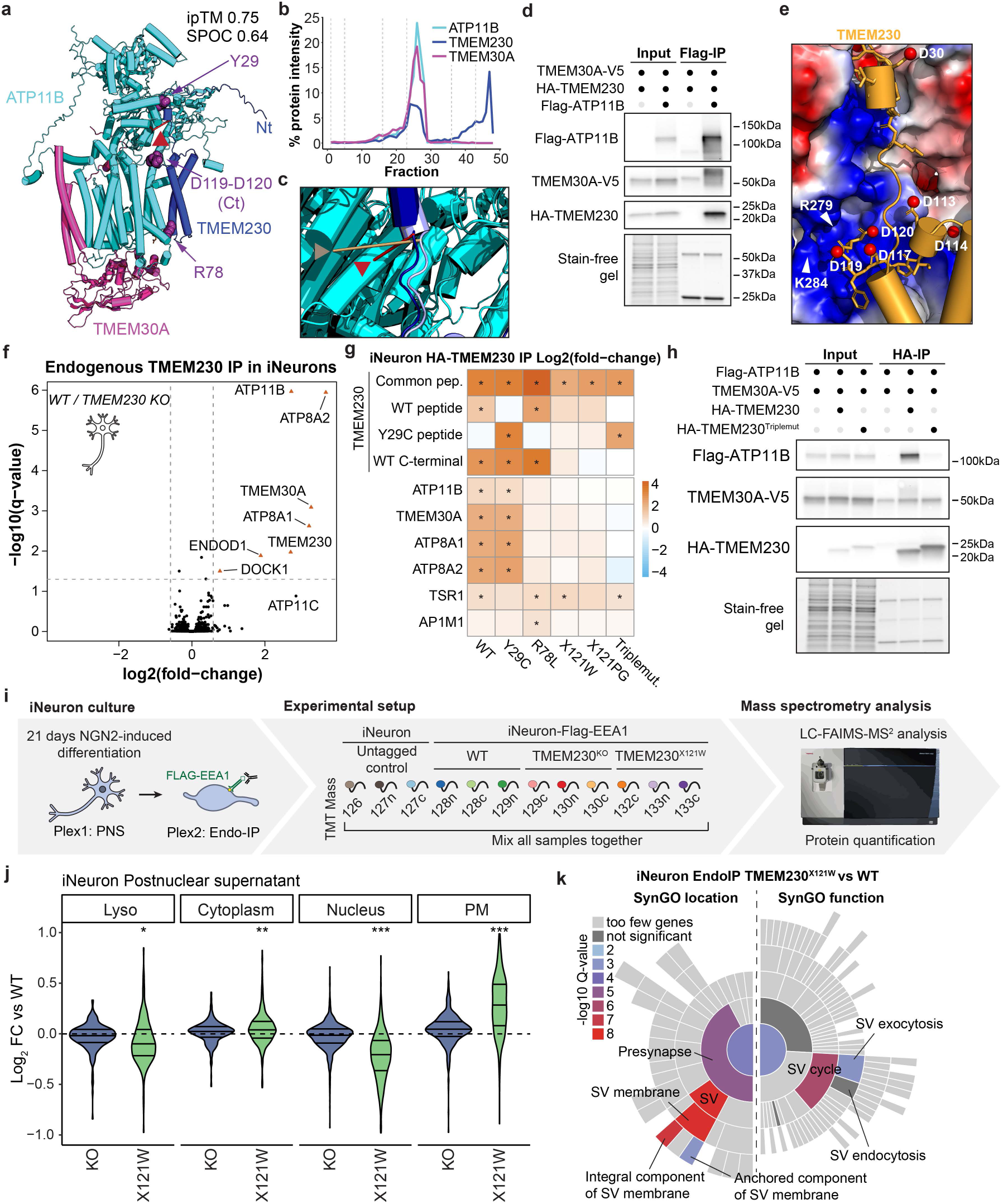
TMEM230 associates with endosomal P4 lipid flippases ATP11B and ATP8A1/2 and interface variants disrupt interaction. **a**, AF-M prediction for TMEM230-ATP11B-TMEM30A, in blue, cyan, and magenta, respectively. TMEM230 Y29, R78, and C-terminal (Ct) D120-D121, as purple space fill, and N-terminal (Nt) are indicated. ipTM/SPOC, ATP11B-TMEM230. **b**, TMEM230-ATP11B-TMEM30A BN-MS profiling. **c**, Overlay of AF-M and AlphaLink2 predictions for TMEM230-ATP11B. AF-M: TMEM230 (light blue), ATP11B (cyan), cross-link (red line and arrowhead). AlphaLink2: TMEM230 (dark blue), ATP11B (teal), cross-link (wheat line and arrowhead). **d**, HA-TMEM230 and Flag-ATP11B co-precipitation after transfection into HEK293 cells. α-Flag immunoprecipitates or input samples were immunoblotted for the indicated proteins. **e**, Basic pocket in ATP11B predicted to interact with the acidic C-terminus of TMEM230 (yellow). Red spheres indicate aspartic residues of TMEM230. **f**, Identification of TMEM230 interacting proteins in iNeurons. Volcano plot showing the proteomic analysis of α-TMEM230 immune complexes from WT H9 compared to H9-TMEM230^-/-^ iNeurons. **g**, Heatmap showing the abundance fold-changes (log_2_) of all significantly enriched proteins in TMEM230 immunoprecipitations in H9-TMEM230^-/-^ iNeurons with or without lentiviral expression of WT and variant HA-TMEM230. Asterisks indicate significantly enriched proteins (q-value<0.05, fold-change>1.5). **h**, Co-precipitation of HA-TMEM230 and HA-TMEM230 carrying mutations in Y29C-R78L-X121W (triple mutant) with Flag-ATP11B and TMEM30A-V5 in transfected HEK293 cells, as examined using immunoblotting of α-HA immune complexes. **i**, Schematic of experimental design for proteomic analysis of early endosomes and postnuclear supernatant (PNS) in 21-day iNeurons derived from WT, TMEM230^-/-^ and TMEM230^X121W^ cells in biological triplicate. **j**, Violin plots (log_2_FC) for the indicated cohorts of proteins of PNS from TMEM230^X121W^ and TMEM230^-/-^ iNeuron, relative to WT cells. Paired t-test pval<0.01 *; pval<0.001 **; pval<0.0001 ***. **k**, SynGO location and function enrichment analysis of protein significantly regulated in early endosomes from TMEM230^X121W^. The indicated categories were significantly enriched (-log_10_Q-value).

TMEM230 was initially reported as a protein linked with rare familial forms of Parkinson’s disease with a potential role in neuronal synaptic activity, and three variants were described (R78L, and two variants X121W and X121P that cause 6-residue C-terminal extensions)^44^. Although the evidence that these variants represent actual causal drivers of Parkinson’s is controversial^45-49^, we were intrigued by the finding that these variants map to the predicted TMEM230-ATP11B interface (**Fig. 3a,e**). TMEM230-R78 is located in proximity with ATP11B-D82 in TM1 and TMEM230 C-terminal (D119-D120) is predicted to bind into a basic pocket of ATP11 (**Fig. 3e**), wherein TMEM230 variants causing C-terminal extension would be expected to sterically disrupt these interactions. To test the impact of these variants on ATP11B interactions and given ATP11B/TMEM230’s apparent role in neuronal function^44,50^, we deleted TMEM230 in human embryonic stem (ES) cells (H9^AAVS1-NGN2;Flag-EEA1^, H9-Flag-EEA1) (**Extended Data Fig. 5d,e**), and converted the cells to cortical-like induced neurons (iNeurons) using the NGN2 driver during lentiviral-based expression of WT or variant TMEM230 (see **METHODS**). TMEM230^WT^ co-immunoprecipitated with ATP11B, ATP8A1/A2 and TMEM30A, compared to TMEM230^-/-^ iNeurons as control (**Fig. 3f, Supplementary Table 4**). In contrast, TMEM230 interactions with TMEM30A, ATP11B and ATP8A1/2 were lost in R78L and both stop codon variants as determined by Tandem Mass Tagging (TMT)-MS (**Fig. 3f,g, Extended Data Fig. 5f-h, Supplementary Table 4**). An apparent polymorphic variant Y29C^44,47,48^ was without effect. Loss of interaction of the triple mutant was also validated in HEK293 cells (**Fig. 3h**). AF-M predicts TMEM230 interaction with ATP8A1 and A2 (ipTM > 0.73) in a manner very similar to that seen with ATP11 isoforms (**Extended Data Fig. 5i**), consistent with loss of interaction in the context of interface variants (**Fig. 3g**).

To examine the effect of TMEM230 variants on early endosomes, we analyzed Endo-IP and post-nuclear supernatant (PNS) proteomes in TMEM230^-/-^ and TMEM230^X121W^ iNeurons^51^ (**Fig. 3i, Extended Data Fig. 5d-e, 5j-m, Supplementary Table 4**). For PNS proteomes, the abundance of PM and synaptic proteins based on the SynGO database were selectively elevated in TMEM230^X121W^ iNeurons relative to WT cells, while minimal abundance changes were found in TMEM230^-/-^ iNeuron (**Fig. 3j, Extended Data Fig. 6a-d**). For endosomal proteomes, we found a lower number of proteins whose abundance was altered compared to PNS (**Extended Data Fig. 6e-g**), and involved synaptic vesicle cycle and its membrane components (**Fig. 3k**) in TMEM230^X121W^ iNeuron endosomes. Proteins whose abundance was increased on early endosomes in the context of TMEM230^X121W^ included several RAB proteins (e.g. RAB3A/B), as well as endocytic cargo (e.g. SORL1), while levels of DNM1/2 involved in endosomal vesicle budding were decreased (**Extended Data Fig. 6g**). ATP8/11 and TMEM30A abundance was unaffected in total or endosomal proteomes (**Extended Data Fig. 6h**). Thus, controversial variants in TMEM230^47-49^ disrupt interactions with multiple lipid flippases and alter the abundance of endosomal and PM proteins in iNeurons.

Next, we extended our disease variant-interface analysis to all pairwise interactions in our dataset. Disease variants in proteins often involve either residues critical for folding, the disruption of which can promote misfolding and reduced stability, or residues that participate in interactions with other proteins. The latter class includes residues that directly contact an interacting protein, or whose mutation alters the conformation of the interaction surface in a way that reduces binding affinity of a partner protein. The large-scale prediction of interaction interfaces provided an opportunity to systematically investigate disease variants located at or nearby subunit interfaces, potentially disrupting proper complex formation. We searched for protein coding candidate disease variants derived from Uniprot that were near predicted interfaces (within 2 amino acids from an interacting residue) (see **METHODS**). We identified 34 such cases involving 53 variants in endosome-related proteins (**Extended Data Fig. 6i,j, Supplementary Table 3**). Examples of variants within interfaces with pairwise SPOC scores typically greater than 0.3 (ipTM scores > 0.49) are shown in **Extended Data Fig. 6j**. Although TMEM230 disease alleles are controversial, we demonstrated that TMEM230-ATP11B interface variants can disrupt physical interactions, suggesting that this approach be useful in identifying interacting partners of disease proteins whose interactions are disrupted by specific variants.

### TMEM9/9B are Core Components of CLCN3/5 Chloride Antiporters

High lumenal chloride (Cl^-^) ion concentrations activate several endolysosomal enzymes and have been proposed to provide counterions to support the V-ATPase generated H^+^-gradient^52-54^. CLCN3-5 Cl^-^-H^+^ antiporters are proposed to function primarily in endosomes, while a heterotetrameric complex composed of CLCN7 α-subunits and OSTM1 β-subunits functions primarily in lysosomes^52,55^. CLCN3 variants are implicated in intellectual disability^56^ and CLCN3 deficiency leads to neurodegeneration in mice^57^. Unlike CLCN7, analogous β-subunits have not been reported for CLCN3-5. EndoMAP.v1 identified cross-links between CLCN3/5 and TMEM9/9B (**Fig. 4a**), a strong enrichment of TMEM9/9B in early endosomes^24,25^ (**Extended Data Fig. 7a**), and co-migration of CLCN3/4/5 and TMEM9/9B in BN-MS (**Fig. 4b**). Pairwise AF-M and AlphaLink2 predicted interaction of CLCN3 or 5 with 2 TM segments from TMEM9 or 9B (SPOC >0.97), including compatible cross-link distances for AF-M (**Fig. 4c, Extended Data Fig. 7b**). Since CLCN proteins form homo and heterodimers^58^, we examined tetrameric predictions of CLCN3 or 5 with TMEM9 or 9B that had the expected antiporter dimer interface, with two molecules of TMEM9 (or 9B) compatible with the cross-link distance constraint (**Fig. 4a,c, Extended Data Fig. 7c,d**). Interestingly, the relative orientation of the 2 TM segments in TMEM9 were distinct from the single TM in OSTM1 (**Extended Data Fig. 7e**). Additionally, the two β-β-α-α-α-β folds of the two TMEM9 molecules occupies a similar location as the helical lumenal “cap” domain of OSTM1, but with a distinct conformation (**Extended Data Fig. 7d,e**).

**Fig. 4.**
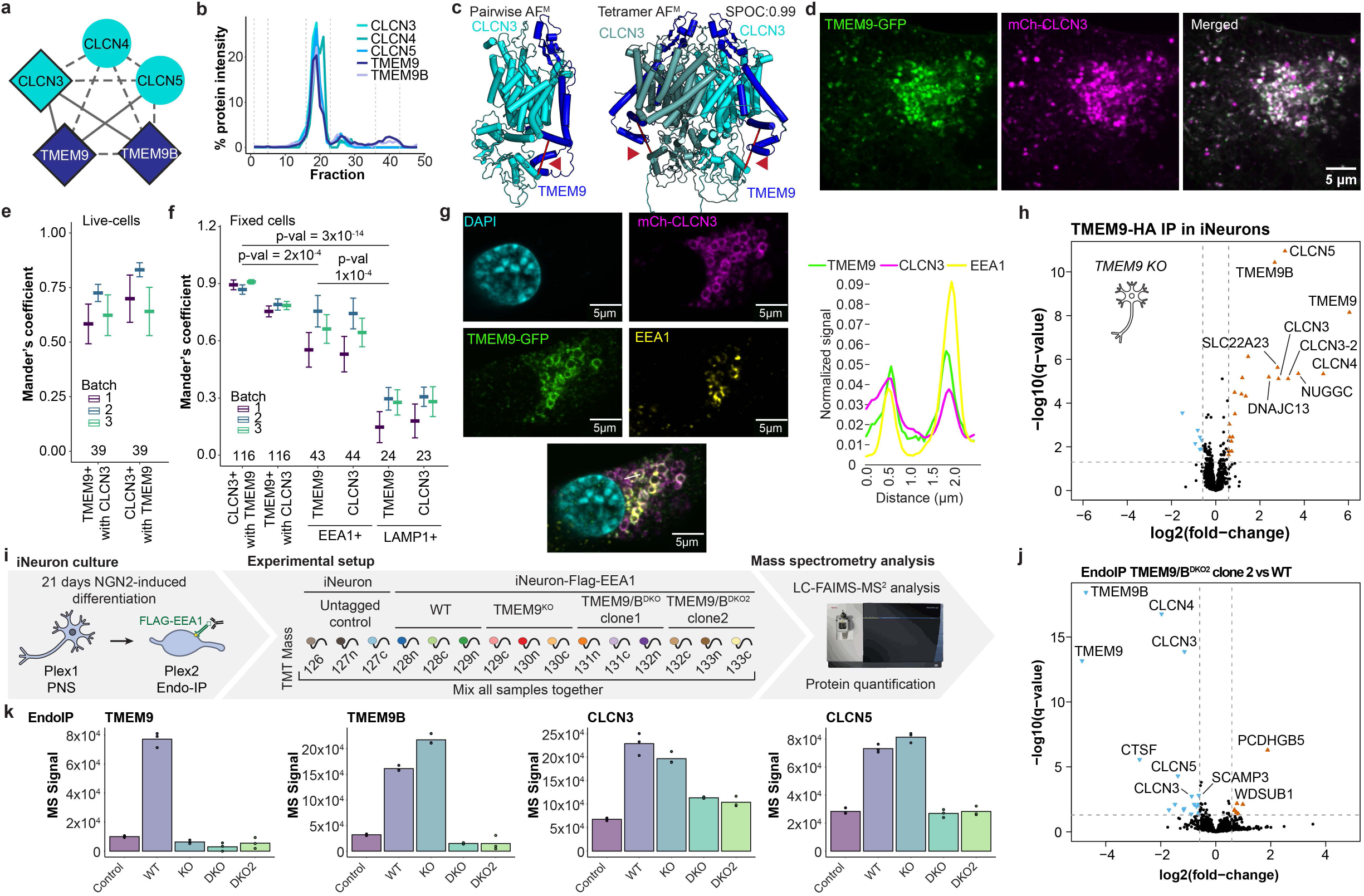
TMEM9/9B are core subunits of endosomal CLCN3/4/5 Cl^-^-H^+^ antiporters. **a**, Summary of EndoMAP.v1 interactions for CLCN3/4/5 and TMEM9/9B. Diamonds and circular nodes represent endosomal and other proteins, respectively. Solid and dotted edges represent interactions identified by at least one cross-link or only co-fractionation, respectively. **b**, BN-MS profiling for CLCN3/4/5 and TMEM9/9B. **c**, AF-M predictions for CLCN3-TMEM9 pair and heterotetramer. The locations of DSSO cross-links are indicated with the red line and arrowhead. **d,e**, Colocalization analysis of TMEM9-GFP and mCh-CLCN3 in SUM159 cells by live-cell imaging. Mander’s coefficient of GFP and mCh puncta are shown in panel **e** (n=39 in 3 independent replicates, mean ± sem), with an example of a cell shown in panel **d**. **f**, Mander’s coefficient analysis of colocalization between TMEM9-GFP, mCh-CLCN3, α-EEA1 and α-LAMP1 in fixed SUM59 cells as determined by immunofluorescence. The number of fields of view across 3 biological replicates is indicated (mean ± sem). **g**, Example of TMEM9-GFP, mCh-CLCN3, and α-EEA1 staining in a cell expressing high levels of CLCN3 (left panel), which promotes the formation of swollen endolysosomes. Line traces show the overlap of the 3 proteins in the limiting membrane of endosomes (right panel). **h**, Volcano plot showing the proteomic analysis of α-HA immunoprecipitations from TMEM9^-/-^ iNeurons with or without lentiviral expression of TMEM9-HA. **i**, Schematic of experimental design for proteomic analysis of early endosomes and PNS in 21-day iNeurons derived from WT, TMEM9^-/-^ and two different clones of TMEM9/9B^DKO^ cells in biological triplicate. **j**, Volcano plot showing the proteomic analysis of Endo-IPs from TMEM9/9B^DKO^ (clone 2) vs WT iNeurons (day 21). **k**, TMT reporter signal intensity for CLCN3/5 and TMEM9/9B in Endo-IPs from iNeurons with the indicated genotypes.

Several experiments further validated CLCN-TMEM9 interactions within early endosomes. First, TMEM9-GFP and mCherry(mCh)-CLCN3 colocalized in vesicles in live (Mander’s coefficient ∼0.64-0.72) (**Fig. 4d,e, Supplementary Video 1**) and fixed cells, where extensive colocalization with EEA1-positive vesicles compared to LAMP1-positive vesicles was observed (Mander’s coefficient ∼0.65 and ∼0.25, respectively) (**Fig. 4f**). Second, TMEM9-GFP tracked with expected swollen endosomes in mCh-CLCN3 overexpressing cells^59^ (**Fig. 4g, Extended Data Fig. 7f**). Third, Flag-CLCN3 or 5 reciprocally associated with HA-tagged TMEM9/9B in HEK293 cells by co-IP (**Extended Data Fig. 7g**) ^28^.

To systematically examine TMEM9 interaction partners, we created TMEM9^-/-^ ES cells (**Extended Data Fig. 7h**) and expressed TMEM9^HA^ in biological quadruplicate day-21 iNeurons prior to TMT-based IP-MS (**Fig. 4h, Supplementary Table 5**). CLCN3, 4, and 5, as well as TMEM9B were all highly enriched in α-HA immunoprecipitates, demonstrating specific interaction of TMEM9 with multiple CLCNs and TMEM9B-containing heterotetramers. Finally, we performed PNS and Endo-IP proteomics for WT, TMEM9^-/-^, and two different clones of TMEM9/9B double knockout (TMEM9/9B^DKO^) iNeurons (**Fig. 4i, Extended Data Fig. 7i-j, Supplementary Table 5**). Early endosome and PNS proteomics revealed a selective abundance reduction of CLCN3/4/5-TMEM9/9B complex, CLCNKA, and CTSF (Cathepsin F) (**Fig 4j,k, Extended Data Fig. 8a-d**), with reduced CLCN3 levels in TMEM9/9B^DKO^ confirmed by immunoblotting (**Extended Data Fig. 8c**). The interaction, co-localization, and selective dependency between CLCN and TMEM9/9B protein levels in iNeurons reveal TMEM9/9B as core components of CLCN antiporter complexes in endosomes and suggests a role in complex stability and/or endosomal trafficking.

### Expanding EndoMAP.v1 via Systematic 3-way Clique Predictomes

To expand the endosomal structural interactome beyond protein pairs, we identified and performed AF-M predictions on all 625 3-way cliques (combinations of 3 proteins interacting with each other) within EndoMAP.v1, with each clique requiring at least one cross-link-supported interaction. This approach yielded 172 predictions containing ≥2 well-predicted interaction interfaces (interface average models >0.5) within each clique (**Fig. 5a, Supplementary Table 6**). Fifty-nine% of these predictions matched interprotein cross-link constraints while an additional 17% involved cross-links within unstructured regions. Illustrating the use of the 3-way clique approach for analysis of complexes with >3 subunits, pairwise and 3-way clique predictions for combinations of endosomal Class II PI3 kinase (UVRAG, BECN1, PIK3C3 and PIK3R4) subunits recapitulate key inter-subunit interactions across the complex^35^, with valid cross-link distances for each pair and 3-way assembly (**Extended Data Fig. 9a, also see Fig. 1i**).

**Fig. 5.**
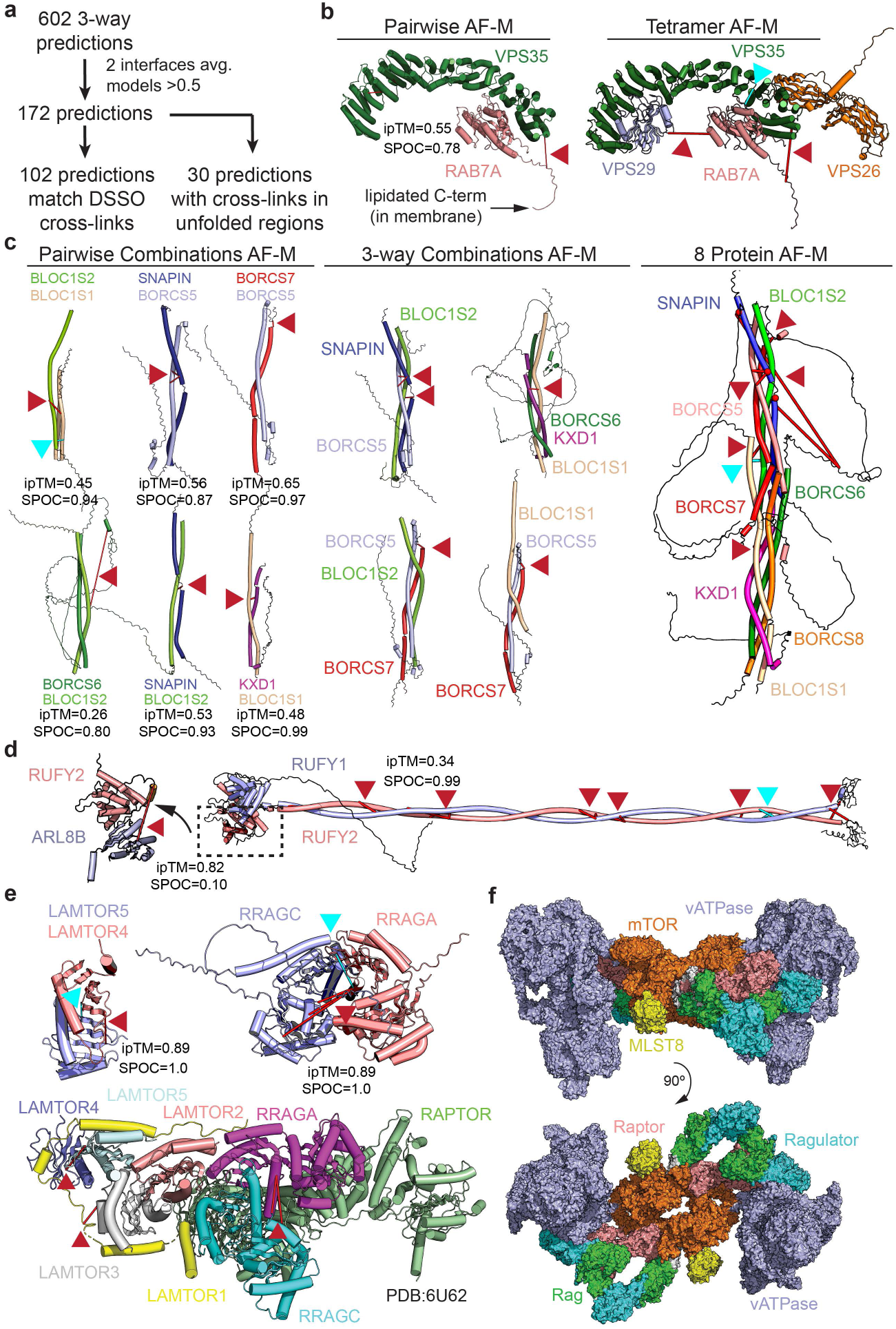
Towards a structural proteomic landscape for early endosomes. **a**, Systematic AF-M structural predictions for 3-way clique assemblies within EndoMAP.v1 and match with the cross-linker distance constraints. **b**, Pairwise AF-M prediction for VPS35-RAB7A (left) and tetramer prediction for Retromer-RAB7A complex (right) and associated cross-links from EndoMAP.v1. ipTM and SPOC scores for pairwise combination are shown. **c**, AF-M structural predictions and interprotein cross-links within the BORC endolysosomal positioning complex. Pairwise AF-M predictions (left), 3-way clique predictions (middle) and 8-protein predictions (right) are shown along with associated interprotein cross-links. ipTM and SPOC scores for pairwise combinations are indicated. **d**, Pairwise AF-M predictions and associated cross-links for a RUFY1-RUFY2 heterodimer (right) and for interaction of the RUFY2 N-terminal helical domain with ARL8B (left). **e**, Pairwise AF-M predictions and associated cross-links for LAMTOR4/5 (top left) and RRAGA/C (top right), with cross-links mapped onto the Ragulator structure (PDB:6U62)^67^ (bottom). **f**, Model for association of MTOR-Ragulator complex (PDB:7UXH)^63^ with V-ATPase^ADP^ (PDB:6WM2)^66^ based on cross-links between LAMTOR2 and LAMTOR4 with ATP6V1C1. Docking model was generated using HADDOCK (see **METHODS**). DSSO and DHSO/DMTMM cross-links are indicated with red and cyan lines and arrowheads, respectively.

Several multi-protein complex predictions were generated for core endolysosomal regulators lacking structural information through the 3-way clique AF-M approach. First, we identified a clique containing Retromer subunits VPS35 and VPS29 and the endosomal GTPase RAB7A. Here, RAB7A directly binds to the concave surface of VPS35 α-solenoid fold (SPOC=0.78), supported by both DSSO and DHSO/DMTMM cross-links, in a manner compatible with simultaneous binding of VPS26A and VPS29 to VPS35 (**Fig. 5b**). AlphaFold3^60^ prediction for VPS35-RAB7^GTP^ closely matches the RAB7A^GTP^ crystal structure and provides a plausible structural mechanism for the previously reported ability of RAB7A^GTP^ to recruit Retromer to endosomes^61^ (**Extended Data Fig. 9b**). Similarly, pairwise and 3-way clique combinations facilitate AF-M-driven assembly of the 8 subunits forming the kinesin-associated endolysosomal positioning BORC complex, for which no structural data has been reported (**Fig. 5c**). The predicted 4-helix bundle with inter-digitated subunits is supported by multiple DSSO and DHSO/DMTMM cross-links (**Fig. 5c**). Acting in opposition to BORC for retrograde endosome trafficking are RUFY (<u>RU</u>N and <u>FY</u>VE domain) proteins, which link ARL8-tethered endolysosomes with dynein motors. Multiple DSSO and DHSO/DMTMM cross-links between RUFY1, RUFY2, and/or ARL8B validate an extended RUFY1-RUFY2 dimeric coil-coil structure (SPOC=0.99), with ARL8B binding the RUN domain (ipTM=0.82, **Fig. 5d**).

SNARE (<u>s</u>oluble <u>N</u>-ethylmaleimide-sensitive factor <u>a</u>ttachment protein <u>re</u>ceptor) components facilitate endolysosomal vesicle fusion and maturation. We identified cross-links defining dozens of pairwise combinations of R-SNARE, Q-SNARE, regulatory and RAB proteins (**Extended Data Fig. 9c**), allowing generation of numerous models with supporting DSSO and DHSO/DMTMM cross-links (**Extended Data Fig. 9d-h**). Pairwise, 3-way, and tetrameric AF-M models of core SNARE complexes formed post-vesicle fusion structures with numerous cross-links found: **1)** between N-terminal Helical a, b, c (Habc) domains of Q-SNAREs (**Extended Data Fig. 9d-f**), **2)** soluble fusion factors, including NAPA (N-ethylmaleimide-sensitive factor attachment protein alpha) and NAPG, which have been reported to co-associate^28,29^, and **3)** HOPS tethering complex subunits including VPS16 (**Extended Data Fig. 9g,h**). Finally, we identified cross-links between SNARE components and two distinct classes of membrane-embedded proteins with extensive data supporting biochemical interactions: multi-TM SCAMP proteins and the single TM protein PTTG1IP (**Extended Data Fig. 9i-n**). SCAMP proteins are known to be involved in vesicle secretion but to our knowledge have not been reported to directly interact with SNAREs. Although pairwise AF-M scores are weak for VAMP2/3-SCAMP1/3 predictions (0.01<SPOC <0.36), there are numerous interactions seen by co-IP^29^ and cross-links in largely unstructured cytosolic regions of SCAMPs consistent with interactions on the cytosolic regions of both R- and Q-SNAREs (**Extended Data Fig. 9i,j**). Similarly, PTTG1IP, which is predicted in Uniprot to have a TM (residues 97-117), was cross-linked to both Q and R-SNAREs, and displayed multiple interactions with SNARE components at endogenous protein levels as annotated in Open Cell^28^ (**Extended Data Fig. 9k**). Pairwise AF-M predictions between PTTG1IP and VAMP3/VAMP8/VTI1B resulted in SPOC scores >0.47, with cross-links within the distance range for the pentameric complex prediction for the PTTG1IP-VAMP8 DSSO cross-link (**Extended Data Fig. 9l-n**). To our knowledge, PTTG1IP has not been previously linked with SNARE functions, although we note that it is enriched in lysosomes based on previous correlation profiling data^77^. Additionally, PTTG1IP abundance in HeLa cells is ∼6% of VAMP3 (see ^77^), suggesting specialized functions. Together, these data suggest a combinatorial interplay between Q/R-SNARE, RABs, soluble fusion/tethering factors, and new candidate TM containing proteins as possible regulators which likely coordinate endosomal maturation events.

Additional predictions allowed us to compiled models for complexes linked with several endosomal functions, including RAB/GEF (**Extended Data Fig. 10a-d**), channel/transporter (**Extended Data Fig. 10e-g**), Adaptor protein (AP) (**Extended Data Fig. 10h**), ESCRT/ubiquitin (**Extended Data Fig 10i**), lumenal cargo (**Extended Data Fig 10j**), HOPS (Homotypic Fusion and Protein Sorting) (**Extended Data Fig. 10k)**, and cargo trafficking assemblies (**Extended Data Fig. 10l**), with experimental validation in purified endosomes via DSSO and DHSO/DMTMM cross-links.

### V-ATPase as an Interaction Hub

Among the most extensively cross-linked complex was the V-ATPase (**Fig. 1h, Extended Data Fig. 11a**), which pumps protons into the endolysosomal lumen to maintain an acidic pH. V-ATPase can co-IP Ragulator complexes (a 5-subunit LAMTOR complex together with RRAGA/B-RRAGC/D GTPase), which bind and regulate MTOR kinase on the endolysosomal membrane^15,62^. We detected multiple DSSO and DHSO/DMTMM cross-links between Ragulator subunits, consistent with its known structure and pairwise AF-M predictions (**Fig. 5e**). Interestingly, we detected cross-links between LAMTOR2 or LAMTOR4 and the ATP6V1C1 subunit of V-ATPase, suggesting that LAMTOR comes into close contact with V-ATPase. Using the cross-linked Lys residues as a guide, we generated a docking model of the extended dimeric Ragulator-MTORC1 complex coupled onto two fully assembled V-ATPase complexes, forming a plausible V-ATPase-MTORC1 “super assembly” (**Fig. 5f; Extended Data Fig. 11b**). This model illustrates an orientation of V-ATPase interacting with MTORC1 complexes compatible with cross-linking data and the proposed organelle membrane topology for MTORC1-Ragulator^63^, highlighting how our approach may capture contacts between large dynamic protein complexes and support the design of further validation experiments.

Additionally, several cross-links were identified between components of the BORC complex and both the V-ATPase and LAMTOR proteins, (**Extended Data Fig. 11a,b**), although the AF-M predictions had low SPOC score (**Supplementary Table 3**). Additionally, LAMTOR3 was found to cross-link with ARL8B, which is known to contribute to BORCs ability to position endolysosomes within cells^30^ (**Extended Data Fig. 11a**). We speculate that V-ATPase, LAMTOR, and ARL8 may form a nexus that can associate with BORC to facilitate endolysosomal trafficking via kinesin motor complexes. Interestingly, our data further validate the previously reported structure of V-ATPase in association with MEAK7^64^, a <u>T</u>BC and <u>L</u>ysM <u>D</u>omain <u>c</u>ontaining (TLDc)-domain protein reported as a modest activator of V-ATPase activity in vitro^65^, including cross-links with V1 subunits ATP6V1D and ATP6V1B2 (**Extended Data Fig. 11c-e**). We also identified cross-links between residues on the cytoplasmic domain of the V0 subunit ATP6V0A1 and multiple endolysososomal RABs (RAB33B, RAB14, RAB4A, RAB5C) (**Extended Data Fig. 11f**), all surpassing the 1% FDR threshold as determined by Scout^38^ (**Supplementary Table 2**), although with low AF-M pairwise SPOC scores. Such an interaction would likely be specific for the V0 complex, as the ATP6V1H subunit would be expected to sterically block RAB association in a fully assembled V-ATPase complex. It is conceivable that these RABs, which are typically present at very high copy number in cells (≥10^5-6^ copies per cell)^24^ and associated with the endosomal membrane through their lipidated C-termini, could interact with ATP6V0A1 in a manner that is membrane facilitated.

## DISCUSSION

By combining protein interactions with cross-link supported structural predictions, EndoMAP.v1 provides a framework for understanding the EEA1-positive endosomal structural interactome. EndoMAP.v1 contains 4,282 interactions based on XL-MS and BN-MS with 229 structural predictions for endosomal interactions without previous structural information. This landscape can be explored through an interactive viewer containing all structural predictions, interactions and experimental data (https://endomap.hms.harvard.edu/) (**Extended Data Fig. 11g**). We demonstrated how EndoMAP.v1 can be used to identify new core subunits of membrane protein complexes, as in the case of TMEM230 and TMEM9/9B. Moreover, we showed how XL-MS can provide experimental support for large-scale hypothesis generating structural predictions in the context of an organelle, where weak protein interactions may be facilitated via membrane tethering. Future studies will further expand and address the limitations of this work, such as inclusion of additional endosome populations, improving the coverage of integral membrane proteins, and addressing the challenge of biochemical and structural validation of proposed models at scale. Finally, the pipeline described here serves as a roadmap for analogous efforts with other organelles and for understanding the diversity of organellar proteomes and interactions in diverse cell types.

## Supporting information

Key Resource Table

## ACKNOWLEDGMENTS

We thank members of the Harper lab for feedback and I.R. Smith for statistical analysis implementation. We thank K.W. Li and F.T.W. Koopmans for discussion and feedback. This work was funded by Aligning Science Across Parkinson’s (ASAP) (J.W.H.), NIH R01NS110395 (J.W.H.), NIH RO1 GM132129 (J.A.P.), and a Rubicon Postdoctoral Fellowship (M.A.G.-L.). Michael J Fox Foundation administers the grant ASAP-000282 and 024268 on behalf of ASAP and itself. For the purpose of open access, the author has applied for a CC-BY public copyright license to the Author Accepted Manuscript (AAM) version arising from this submission. J.C.W. is a Howard Hughes Medical Institute Investigator and an American Cancer Society Research Professor. We acknowledge the Core for Imaging Technology and Education (Harvard Medical School) for imaging assistance.

## AUTHOR CONTRIBUTIONS

Conceptualization: M.A.G.-L., J.W.H.; Investigation: M.A.G.-L., E.S., J.A.P., E.M.W., Y.J.; Analysis: M.A.G.-L., E.S., E.M.W., J.W.H.; Visualization and website: E.S., M.A.G.-L.; Writing—original draft: M.A.G.-L., J.W.H.; Writing—reviewing and editing: M.A.G.-L., E.S., J.A.P., J.C.W., J.W.H.

## COMPETING INTERESTS

J.W.H. is a co-founder for Caraway Therapeutics (a subsidiary of Merck Inc) and is a scientific advisory board member for Lyterian Therapeutics. All other authors have no competing interests to declare.

## DATA AVAILABILITY

All the mass spectrometry proteomics data (289 .RAW files) have been deposited to the ProteomeXchange Consortium via the PRIDE repository and will be released upon publication. All analyzed proteomic data are in Supplementary Tables 2, 4 and 5, and have been deposited at Zenodo (10.5281/zenodo.14444100). A key resource table containing all resources used in this study is deposited at Zenodo (10.5281/zenodo.14180546).

All AF-M and AlphaLink2 predictions can be downloaded from https://endomap.hms.harvard.edu. AF-M predictions have also been deposited at Zenodo (10.5281/zenodo.14447604). AlphaLink2 predictions have been deposited at Zenodo (10.5281/zenodo.14632928). Input/output files used for modeling mTORC1-Ragulator-VATPase complex using HADDOCK2.4 have been deposited in Zenodo (10.5281/zenodo.14679635). Original imaging files have been deposited at Zenodo (10.5281/zenodo.14826177 and 10.5281/zenodo.14828026) ES cells reported here will be made available upon request, but require an MTA from WiCell.

## Code and Software Availability

Scripts for endosomal proteome meta-analysis, co-fractionation, protein network and co-localization analysis has been deposited on Github (https://github.com/harperlaboratory/EndoMAP) and annotated at Zenodo (10.5281/zenodo.14444100) and will be available upon publication.

**Supplementary Table 1.** Endosomal proteome meta-analysis, label-free Endo-IP proteomics and disease over-representation analysis of the endosomal proteome.

**Supplementary Table 2.** Master dataset for construction of EndoMAP.v1 including proteomic data for individual BN-MS analyses, XL-MS analyses and AF analyses.

**Supplementary Table 3.** Large-scale AlphaFold Multimer predictions of cross-linked proteins.

**Supplementary Table 4.** Proteomic analysis of TMEM230 interacting proteins and proteomic profiling WT, TMEM230^-/-^, and TMEM230^X121W^ iNeurons.

**Supplementary Table 5.** Proteomic analysis of TMEM9 interacting proteins and proteomic profiling WT, TMEM9^-/-^, and two different clones of TMEM9/9B^DKO^ iNeurons.

**Supplementary Table 6.** AlphaFold Multimer predictions of 3-way clique within EndoMAP.v1 and confidence measurements.

**Supplementary Video 1.** Live-cell imaging showing subcellular localization of mCh-CLCN3 and TMEM9-GFP in SUM159 cells (t=2min).

## Extended Data Figure LEGENDS

**Extended Data Fig. 1.**
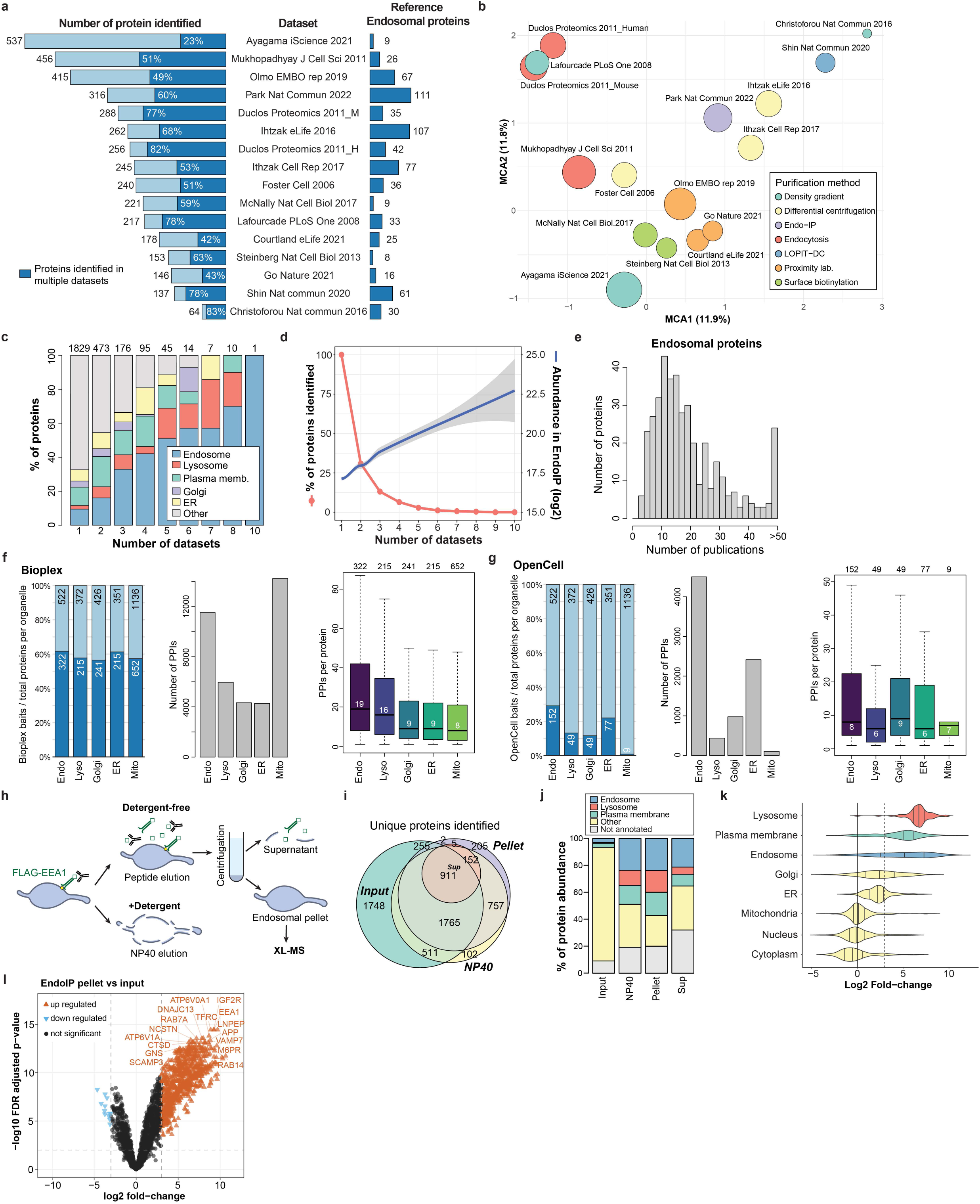
Meta-analysis of the endosomal proteome and optimization of large-scale Endo-IP. **a**, Overview of datasets used for meta-analysis, including the number of proteins identified in each and across datasets, and the number of well-known endosomal proteins identified in the indicated studies. **b**, Multiple correspondence analysis (MCA) showing an overview of the relationship among datasets from panel **a**. Each node represents a dataset color-coded by isolation method and proportional size to the total number of proteins identified. **c**, Bar plot depicting the number of proteins identified across multiple datasets for several subcellular compartments. **d**, Line graph showing the percentage of proteins identified across all 16 data sets in panel **a** and their protein abundance in our Endo-IP experiments from HEK293 cells represented as loess regression line (**Supplementary Table 1**). **f,g**, Number of bait proteins, protein-protein interactions (PPIs), and PPIs per protein in Bioplex^28^ (panel **f**) and Open Cell^29^ (panel **g**) according to organelle assignment. **h**, Schematic of the purification steps from Endo-IP to endosomal pellet used for complexomics. **i,j**, Number of proteins identified (panel **i**) and abundance per compartment (panel **j**) in endosomal pellet compared to the input (PNS), supernatant, or NP40 eluate from the Endo-IP as depicted in panel **h**. **k**, Violin plot showing the fold-change enrichment of proteins from individual organelle compartments in endosomal pellets compared to input (PNS). **l**, Volcano plot showing fold-changes and FDR adjusted p-value for proteins in endosomal pellets compared to input (PNS).

**Extended Data Fig. 2.**
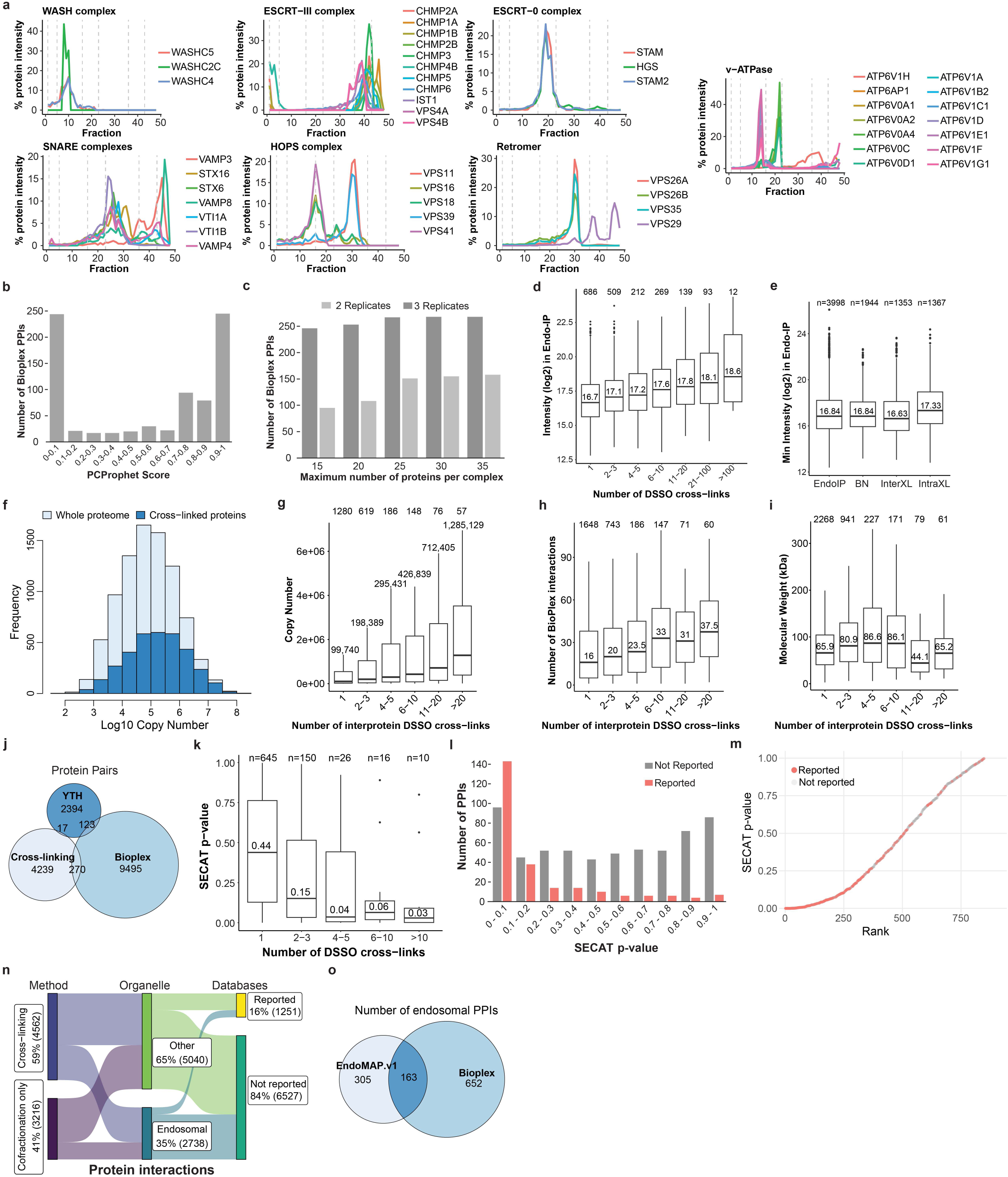
Application of correlation profiling and cross-linking proteomics to endosomes purified by Endo-IP. **a**, Co-fractionation profiles of selected protein complexes from BN-MS. **b**, Number of Bioplex interactions identified by BN-MS compared to co-fractionation PCProphet scores. **c**, Number of Bioplex interactions identified using PCProphet in either 2 or 3 replicates of the Endo-IP BN-MS compared to the maximum number of proteins per complex allowed in the analysis. **d**, Box plot depicting the protein MS signal intensity in Endo-IP compared to the number of DSSO cross-links identified for each protein. **e**, Box plot depicting the minimum protein MS signal intensity for PPIs identified by BN and DSSO cross-linking compared to all proteins identified in Endo-IP. **f**, Distribution of protein copy number (log_10_)^24^ for cross-linked proteins compared to the whole proteome. **g-i**, Box plots depicting the protein copy number (**g**), number of interactors in BioPlex (**h**), and molecular weight (**i**) compared to the number of interprotein DSSO cross-links identified for each protein. **j**, Venn diagram showing the number of protein pairs identified by yeast two hybrid (YTH), Bioplex, and cross-linking proteomics for the same set of proteins. **k**, Boxplot showing co-fractionation SECAT p-values for cross-linked proteins identified by different number of DSSO cross-links. **l,m**, Number (panel **l**) and rank (panel **m**) of cross-linked protein interactions that have been previously reported (or not) compared to their co-fractionation SECAT p-value. **n**, Overview of protein interactions within EndoMAP.v1 including the method, organelle and previous reports. **o**, Venn diagram showing the overlap of endosomal interactions between EndoMAP.v1 and Bioplex for interactions in which both proteins are present in both datasets.

**Extended Data Fig. 3.**
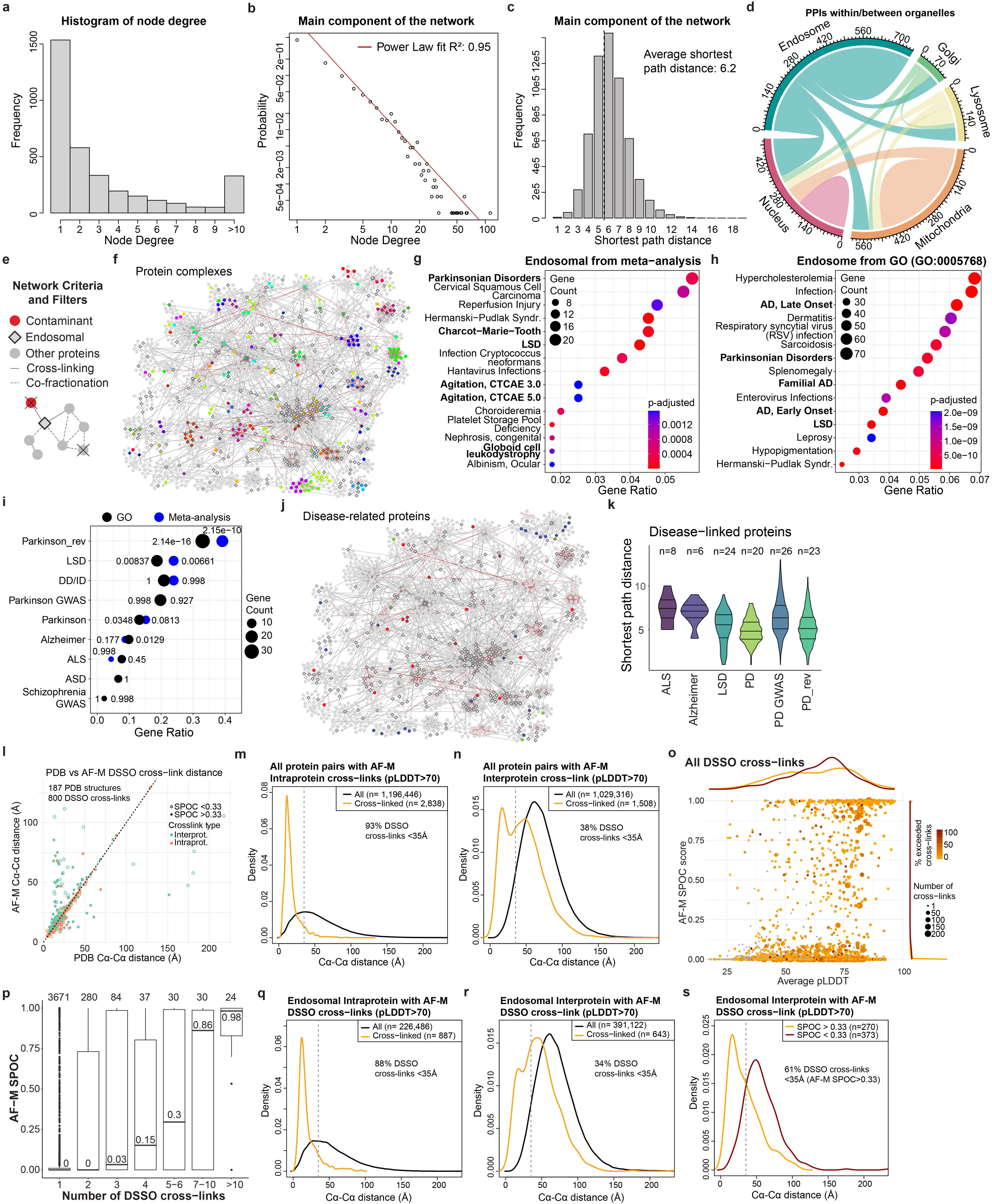
EndoMAP.v1 network characterization and application of AlphaFold-M across DSSO cross-linked protein pairs. **a**, Degree distribution (number of edges per node) of the complete network. **b**, Power law log-log plot of the complete network showing the degree of a node (number of edges) and the probability. **c**, Distribution of the shortest path distances between all proteins in the complete interaction network. **d**, Distribution and number of PPIs within and between selected organelles. **e**, Criteria for network filtering to create an integrated endosomal network (EndoMAP.v1, see **METHODS**). **f**, Mapping of known protein complexes from CORUM^114^ onto the core components of the EndoMAP.v1 network. **g,h**, DisGeNET enrichment analysis of endosomal proteins as defined in our meta-analysis (panel **g**) and Gene Ontology (GO:0005768, panel **h**). Top 15 categories by highest gene ratio are depicted. Disorders related to the nervous system are indicated in bold. **i**, Enrichment analysis of the endosomal proteome within several neurodegenerative diseases (LSD, Lysosomal Storage Disorders; ALS, Amyotrophic Lateral Sclerosis, PD, Parkinson’s disease; ASD, Autism Spectrum Disorders; DD/ID, epilepsy and severe neurodevelopmental disorder). **j**, Mapping of neurodegenerative disease related proteins onto the core component of EndoMAP.v1 network (see **METHODS**). **k**, Distribution of shortest path distances within various classes of neurodegenerative disease related proteins. Three different sources of disease genes were used to retrieve proteins related to PD (see **METHODS**). **l**, Distances between DSSO cross-linked lysines for AF-M predictions compared to structures in the PDB. Green and orange dots represent interprotein and intraprotein cross-links, respectively. Filled and empty dots represent predictions with SPOC > 0.33 or SPOC < 0.33, respectively. **m**, Distribution of Cα-Cα distances (Å) for intraprotein DSSO cross-linked lysines in all AF-M predictions compared to all lysines. **n**, Distribution of Cα-Cα distances (Å) for interprotein DSSO cross-linked lysines in all AF-M predictions compared to all lysines. **o**, Distribution of SPOC scores and average pLDDT for all AF-M predictions. Number of DSSO cross-links evaluated and exceeding the cross-linker distance restrain are indicated by point size and the color, respectively. **p**, Box plot showing the distribution of SPOC scores relative to the number of DSSO cross-links identified for each interaction. **q,r**, Distribution of Cα-Cα distances (Å) for intraprotein (**q**) and interprotein (**r**) DSSO cross-linked lysines in AF-M predictions involving endosomal proteins compared to all lysines. **s**, Distribution of Cα-Cα distances (Å) for interprotein DSSO cross-links reflecting predictions involving endosomal proteins with SPOC > 0.33 (orange) and SPOC < 0.33 (red).

**Extended Data Fig. 4.**
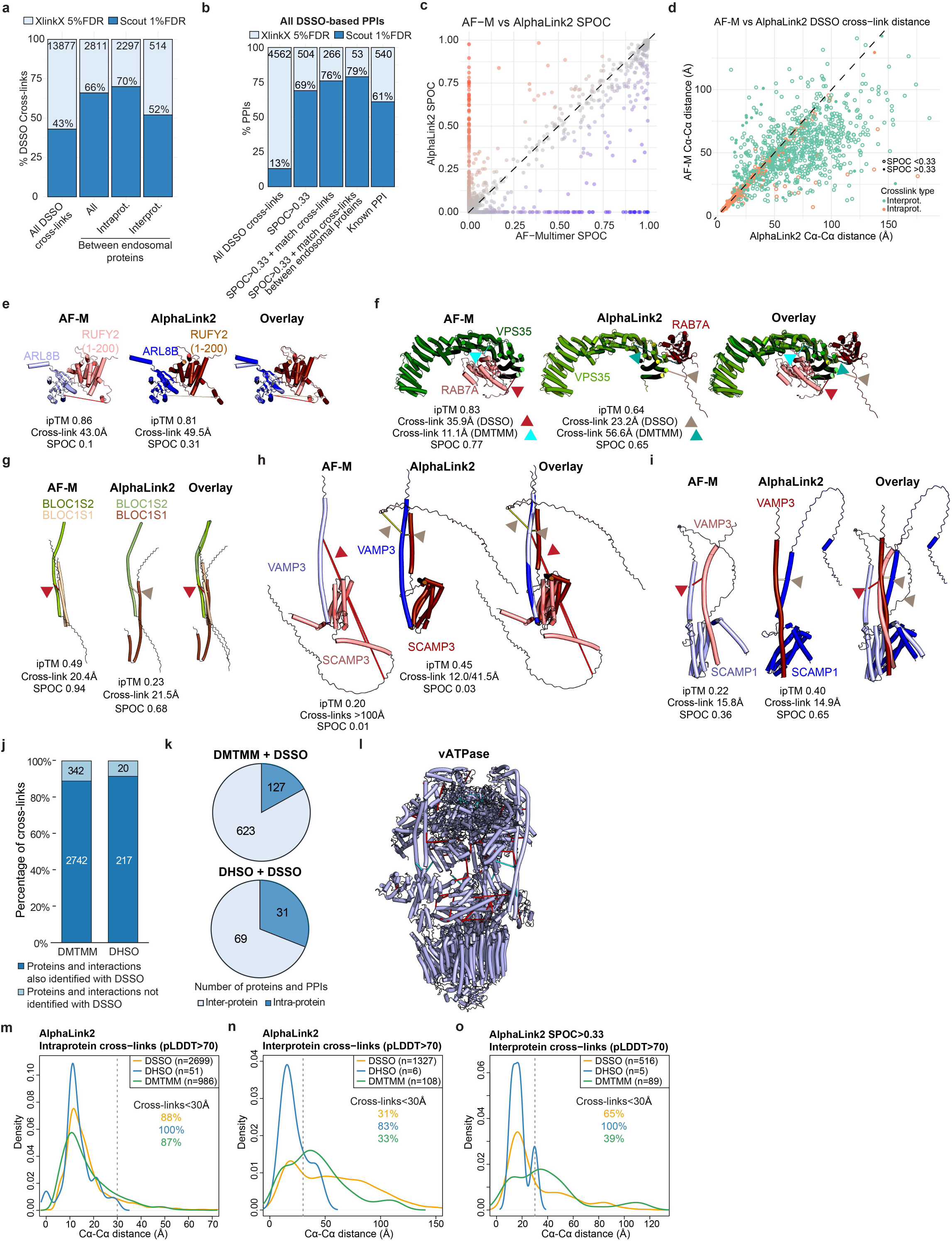
EndoMAP.v1 extension by AlphaLink2 and XL-MS using DHSO/DMTMM cross-linkers. **a**, Overlap of DSSO cross-linking data analyzed using XlinkX at 5% FDR compared to Scout at 1%FDR. **b**, Number of protein interactions based on DSSO cross-links identified with XlinkX and Scout for known interactions and across the selection criteria used in EndoMAP.v1 (i.e. filtering for AF-M score, endosomal protein and cross-link distance). **c**, ipTM scores for AF-M compared to AlphaLink2 predictions. Color gradient represents the score difference; higher in AlphaLink2 (red) or AF-M (blue). **d**, Distances between DSSO cross-linked lysines for AF-M compared to AlphaLink2 predictions. Green and orange dots represent interprotein and intraprotein cross-links, respectively. **e-i**, Individual and overlay AF-M and AlphaLink2 predictions for several protein pairs. DSSO and DHSO/DMTMM interprotein cross-links are indicated with red and cyan lines and arrowheads, respectively. **j**, Mapping DHSO/DMTMM cross-linking data to the proteins and interactions identified with DSSO. **k**, Pie chart showing the number of protein pairs identified with both DMTMM and DSSO (top) or DHSO and DSSO (bottom). **l**, Identified DSSO (red) and DHSO/DMTMM (cyan) cross-links mapped into the endolysosomal V-ATPase (PDB:6WM2)^66^. **m,n**, Distribution of Cα-Cα distances (Å) for intraprotein (**m**) and interprotein (**n**) cross-linked residues in AlphaLink2 predictions. **o**, Distribution of Cα-Cα distances (Å) for interprotein cross-links reflecting predictions with SPOC > 0.33.

**Extended Data Fig. 5.**
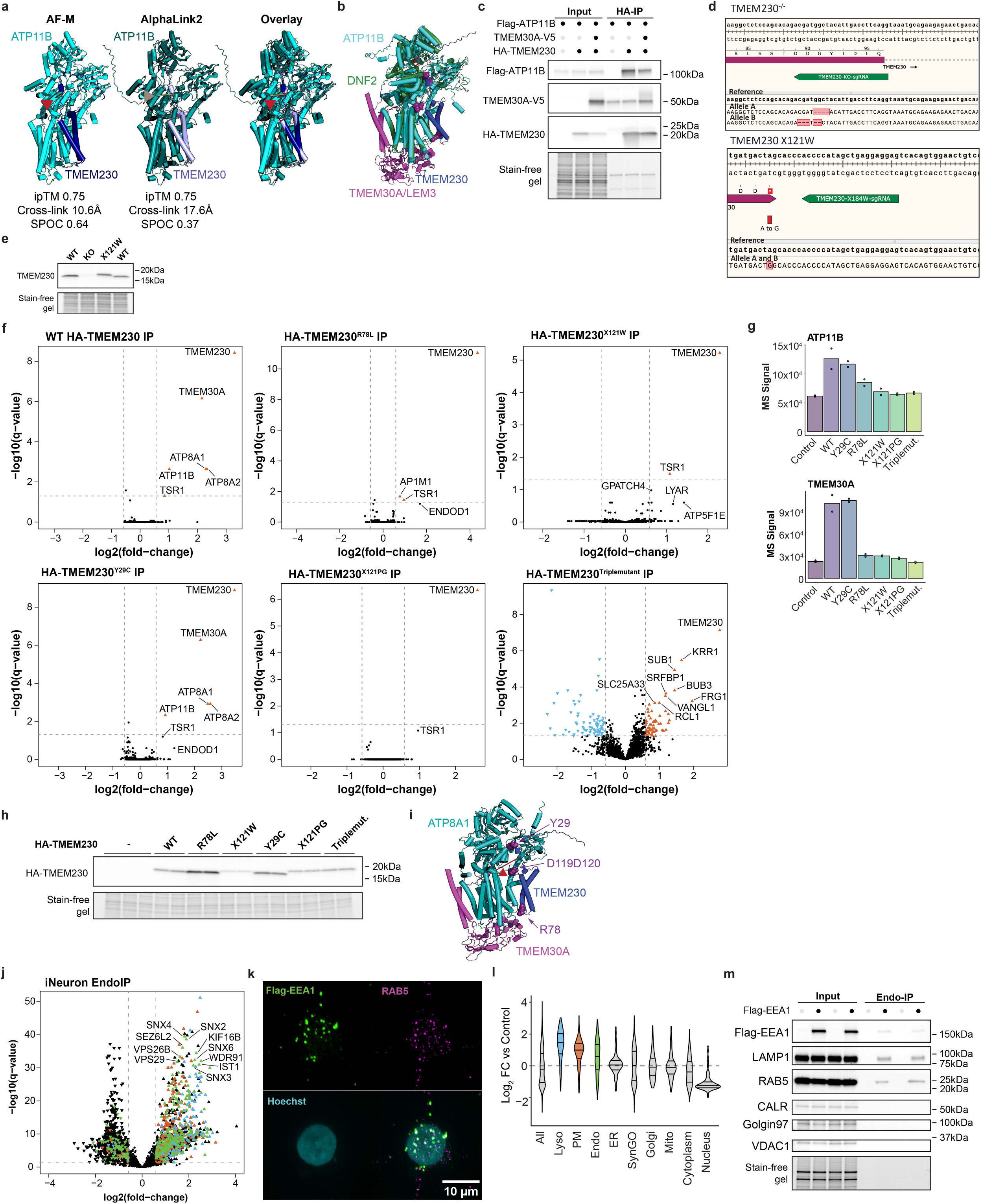
Interface variants disrupt interaction of TMEM230 with endosomal P4 lipid flippases ATP11B and ATP8A1/2. **a**, Individual and overlay AF-M and AlphaLink2 predictions for TMEM230 and ATP11B. AF-M: TMEM230 (light blue), ATP11B (cyan), cross-link (red line and arrowhead). AlphaLink2: TMEM230 (dark blue), ATP11B (teal), cross-link (wheat line and arrowhead). **b**, Overlay of yeast DNF1-LEM3 structure (PDB:7DRX) in the EP2 conformation with AF-M prediction for ATP11B-TMEM30A-TMEM230. **c**, Co-precipitation of Flag-ATP11B and TMEM30A-V5 with HA-TMEM230. The indicated plasmids were transfected into HEK293 cells and α-HA immunoprecipitates or input samples were immunoblotted for the indicated proteins. **d**, Sequence validation of TMEM230^-/-^ and TMEM230^X121W^ clones in H9^AAVS1-NGN2;Flag-EEA1^ cells (H9-Flag-EEA1), showing the location of the sgRNA used (green) and base pairs deleted to create an out of frame mutation and point mutation, respectively. **e**, Immunoblot of total cell lysates from the indicated H9-Flag-EEA1 cell lines probed with α-TMEM230. The X121W mutation adds a six-residue extension (WHPPHS), which can be detected as a band with slightly higher molecular weight. Stain-free gel was used to indicate equal loading of extracts. **f**, Volcano plots (log_2_FC relative to TMEM230^-/-^ cells) of TMEM230 immunoprecipitations in H9-TMEM230^-/-^ iNeurons with or without lentiviral expression of WT and interface variant HA-TMEM230 proteins. **g**, Mass spectrometry (MS) TMT reporter signal for ATP11B and TMEM30A in the indicated TMEM230 variant immunoprecipitation from iNeurons. Dots indicate individual biological replicates. **h**, Immunoblots of total cell extracts from TMEM230^-/-^ iNeurons transduced with lentiviruses expressing the indicated variants of HA-TMEM230 protein. Stain-free gel was used as loading control. **i**, AF-M prediction for a TMEM230-ATP8A1-TMEM30A complex (Y29, R78, and C-terminal D120-D121, purple space fill). The location of a cross-link between ATP8A1 and TMEM30A is indicated by the red line and arrowhead. ipTM = 0.74 for ATP8A1-TMEM230 prediction. **j**, Volcano plot for Endo-IP proteomic analysis from H9-Flag-EEA1 iNeurons (21 days). Proteins annotated as endosomal (green), lysosomal (blue), or plasma membrane (PM, orange) are indicated. **k**, Immunofluorescence microscopy showing the colocalization of Flag-EEA1 (green) with RAB5 (magenta) in iNeurons from H9-Flag-EEA1 cells. **l**, Violin plot showing the fold-change enrichment (log_2_) of proteins from individual organelle compartments (color-coded as panel **j**) in Endo-IP samples from H9-Flag-EEA1 iNeurons (day 21). **m**, Immunoblots of Endo-IP or input samples (PNS) from H9-Flag-EEA1 iNeurons and untagged H9 control (21 days). Blots were probed with the indicated antibodies.

**Extended Data Fig. 6.**
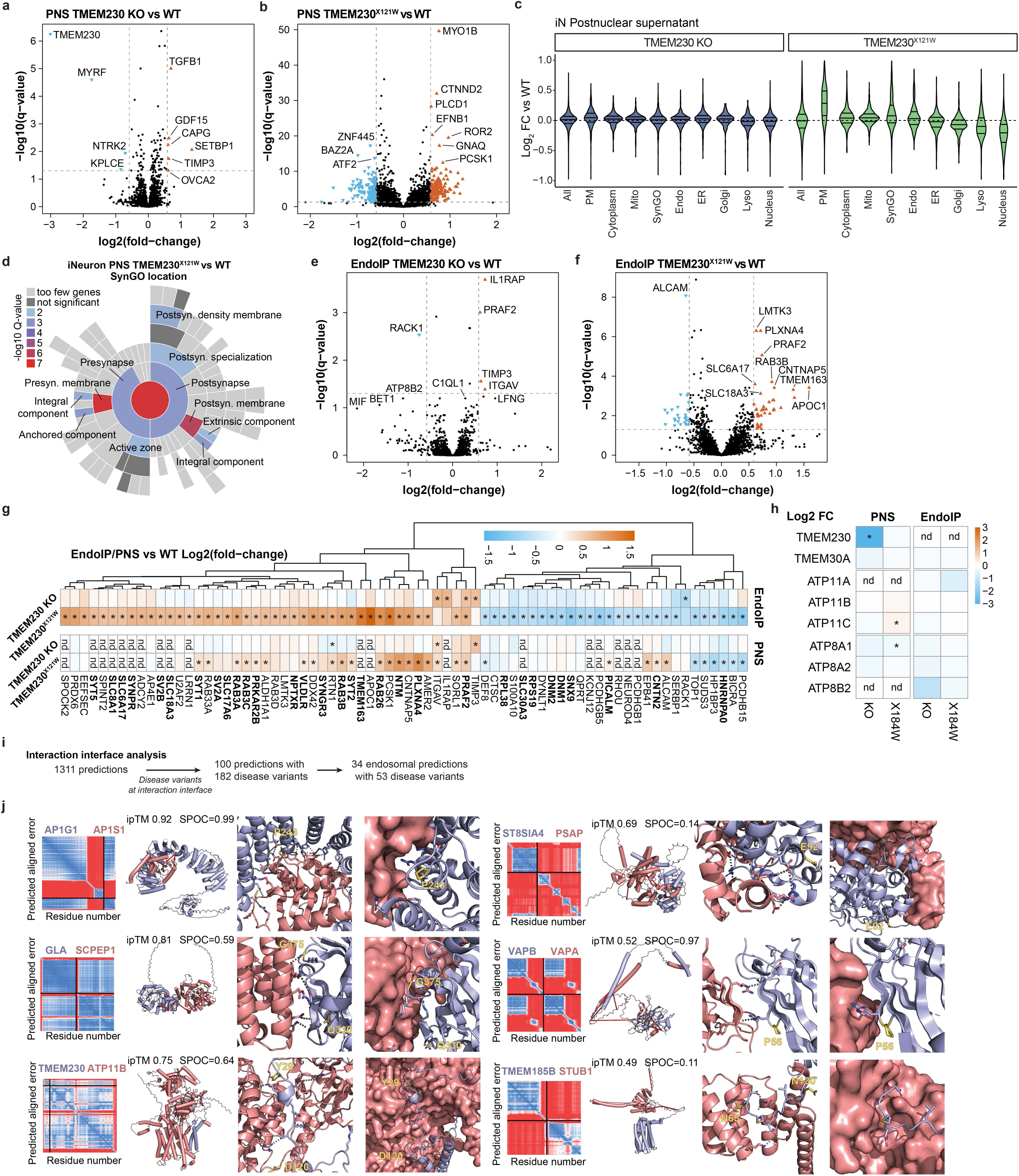
Proteomic profiling of postnuclear supernatant (PNS) and Endo-IP from TMEM230 mutant iNeurons. **a,b**, Volcano plots of PNS proteomic analysis from TMEM230^-/-^ (panel **a**) and TMEM230^X121W^ (panel **b**) iNeurons compared to WT (day 21). **c**, Violin plot showing the fold-change enrichment (log_2_) of proteins from individual organelle compartments in PNS from TMEM230^-/-^ and TMEM230^X121W^ iNeurons compared to WT (day 21). **d**, SynGO location enrichment analysis of protein significantly regulated in PNS from TMEM230^X121W^. The indicated categories were significantly enriched (-log_10_Q-value). **e,f**, Volcano plots of Endo-IP proteomic analysis from TMEM230^-/-^ (panel **e**) and TMEM230^X121W^ (panel **f**) iNeurons compared to WT (day 21). **g**, Heatmap showing the abundance fold-changes (log_2_) for all significantly regulated proteins in Endo-IPs from TMEM230^-/-^ or TMEM230^X121W^ iNeurons (21 day) compared to WT. Synaptic proteins annotated in SynGO (see **METHODS**) are indicated in bold. Asterisks indicate significantly regulated proteins (q-value < 0.05 and fold-change > 1.5). Abundance fold-changes in PNS are also indicated, except for proteins not detected (nd). **h**, Heatmap for the abundance fold-changes (log_2_FC) of selected proteins in PNS and Endo-IPs from TMEM230^-/-^ or TMEM230^X121W^ iNeurons (21 day) compared to WT. Asterisks indicate significantly regulated proteins (q-value < 0.05 and fold-change > 1.5) and nd for proteins not detected. **i**, Summary of pairwise AF-M predictions harboring disease variants within 2 amino acids of the interface for endosomal and non-endosomal proteins. **j**, Disease variants at the interaction interface of pairwise protein AF-M predictions. Predicted aligned error plots (left), predicted structures with ipTMs (center left, interprotein DSSO cross-links indicated by red lines) and close-up view of disease variant residues (yellow) at the interaction interface (center right, and right; dotted lines indicate predicted hydrogen bonds).

**Extended Data Fig. 7.**
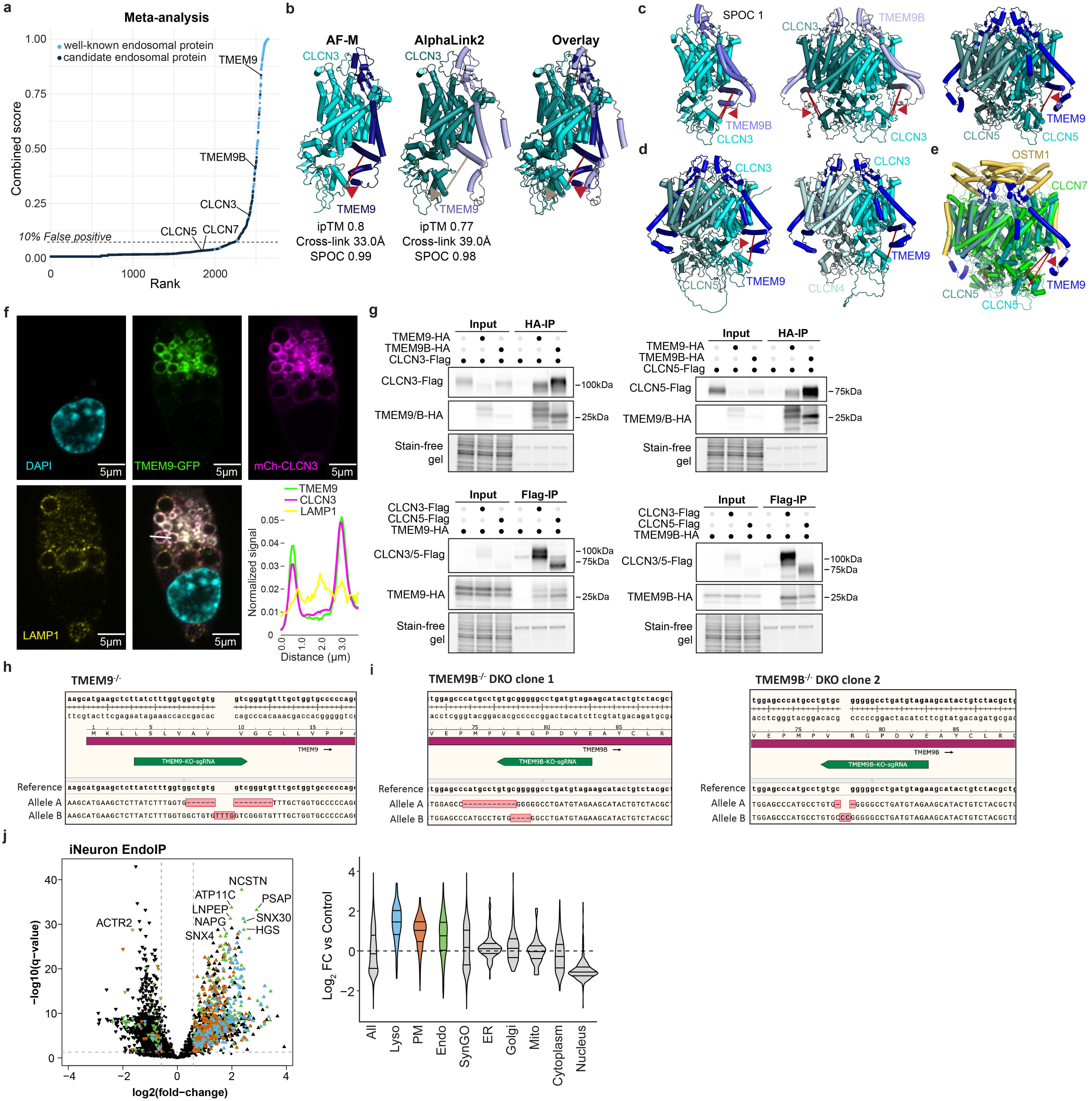
TMEM9/9B are core subunits of endosomal CLCN3/4/5 Cl^-^-H^+^ antiporters. **a**, Meta-analysis score and rank of TMEM9/9B and CLCN3/5/7, with higher values corresponding to endosomal proteins. **b**, Individual and overlay AF-M and AlphaLink2 predictions for CLCN3-TMEM9. AF-M: TMEM9 (dark blue), CLCN3 (cyan), cross-link (red bar and arrowhead). AlphaLink2: TMEM9 (light blue), CLCN3 (teal), cross-link (wheat bar and arrowhead). **c**,**d** AF-M predictions for CLCN3-TMEM9B and selected CLCN-TMEM9/9B heterotetramers. The locations of DSSO cross-links are indicated with the red line and arrowhead. The location of variants found in CLCN5 in Dent’s Disease retrieved from UniProt are shown in red (right, panel **c**). **e**, Overlay of the CLCN5-TMEM9 heterotetramer prediction with the CLCN7-OSTM1 heterotetramer structure (PDB: 7JM7). **f**, Example of TMEM9-GFP, mCh-CLCN3, and α-LAMP1 staining in a cell expressing high levels of CLCN3, which promotes the formation of swollen endolysosomes. Line traces show the overlap of the 3 proteins in the limiting membrane of endolysosomes (bottom right panel). **g**, Co-precipitation of CLCN3/5-Flag and TMEM9/9B-HA. The indicated plasmids were transfected into HEK293 cells and α-HA or α-Flag immunoprecipitations or input samples were immunoblotted with the indicated antibodies. Loading controls as stain-free gels are shown. **h,i**, Sequence validation of H9 TMEM9^-/-^ cells, showing the location of the sgRNA used (green) to create frameshift mutations in TMEM9 (panel **h**) and subsequently, used the indicated sgRNA (green) to create frameshift mutations in TMEM9B (panel **i**). **j**, (Left) Volcano plot for Endo-IP proteomic analysis from H9-Flag-EEA1 iNeurons. (Right) Violin plot showing the fold-change enrichment (log_2_) of proteins from individual organelle compartments in Endo-IP from H9-Flag-EEA1 iNeurons. Proteins annotated as endosomal (green), lysosomal (blue), or plasma membrane (PM, orange) are indicated.

**Extended Data Fig. 8.**
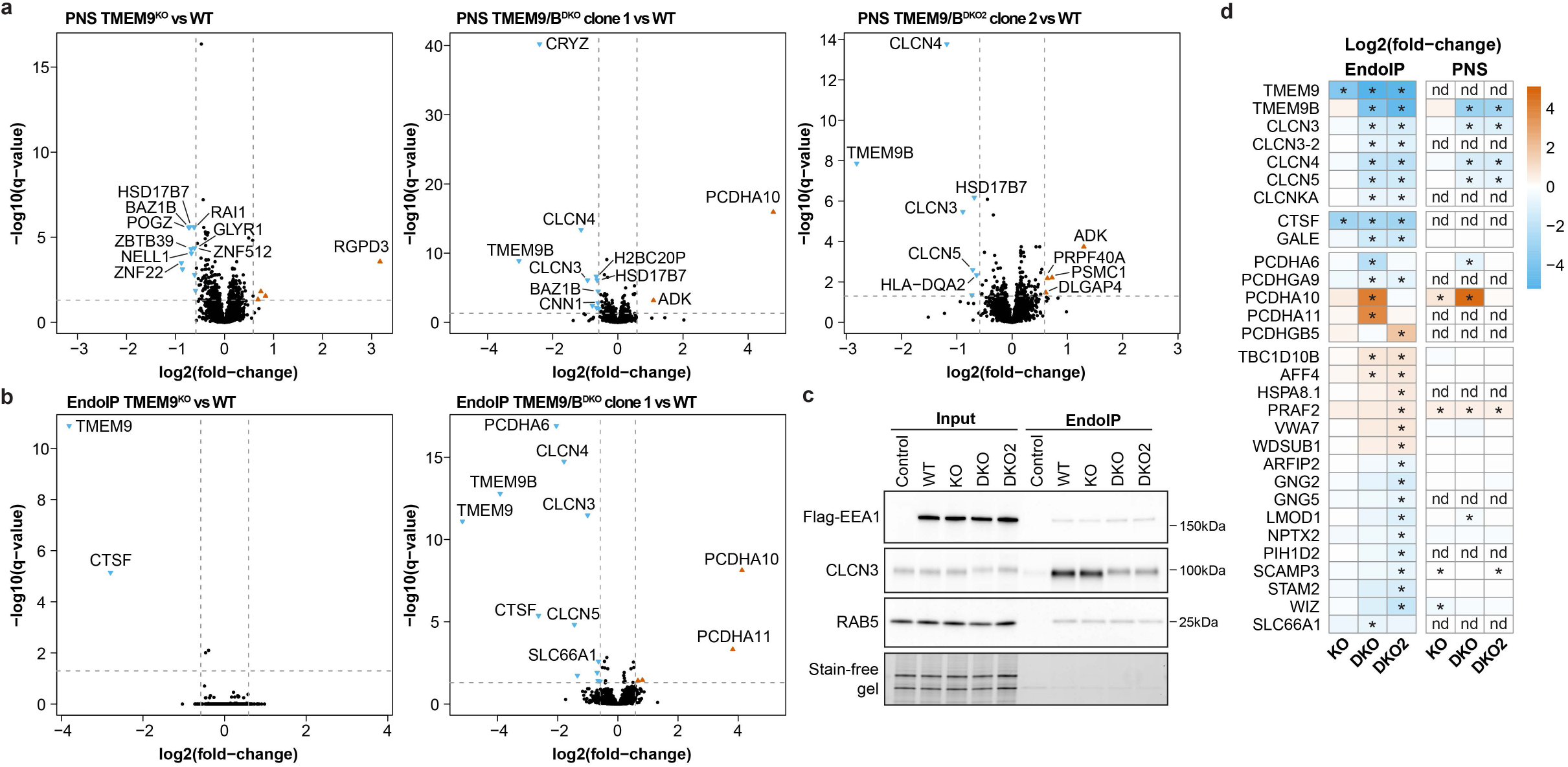
Proteomic profiling of postnuclear supernatant (PNS) and Endo-IP from TMEM9^-/-^ and TMEM9/9B^DKO^ iNeurons. **a**, Volcano plots for PNS (post-nuclear supernatants) proteomic analysis from TMEM9^-/-^ and two clones of TMEM9/9B^DKO^ iNeurons (day 21) compared to WT. **b**, Volcano plots for Endo-IP proteomic analysis from TMEM9^-/-^ and one clone of TMEM9/9B^DKO^ iNeurons compared to WT. **c**, Immunoblots of input and Endo-IP samples from the experiment outlined in **Fig. 4i**. Blots were probed with the indicated antibodies. **d**, Heatmap showing the abundance fold-changes (log_2_) for all significantly regulated proteins in Endo-IPs from TMEM9^-/-^ and TMEM9/9B^DKO^ iNeurons (day 21) compared to WT. Asterisks indicate significantly regulated proteins (q-value < 0.05 and fold-change > 1.5). Abundance fold-changes in PNS are also indicated, except for proteins not detected (nd).

**Extended Data Fig. 9.**
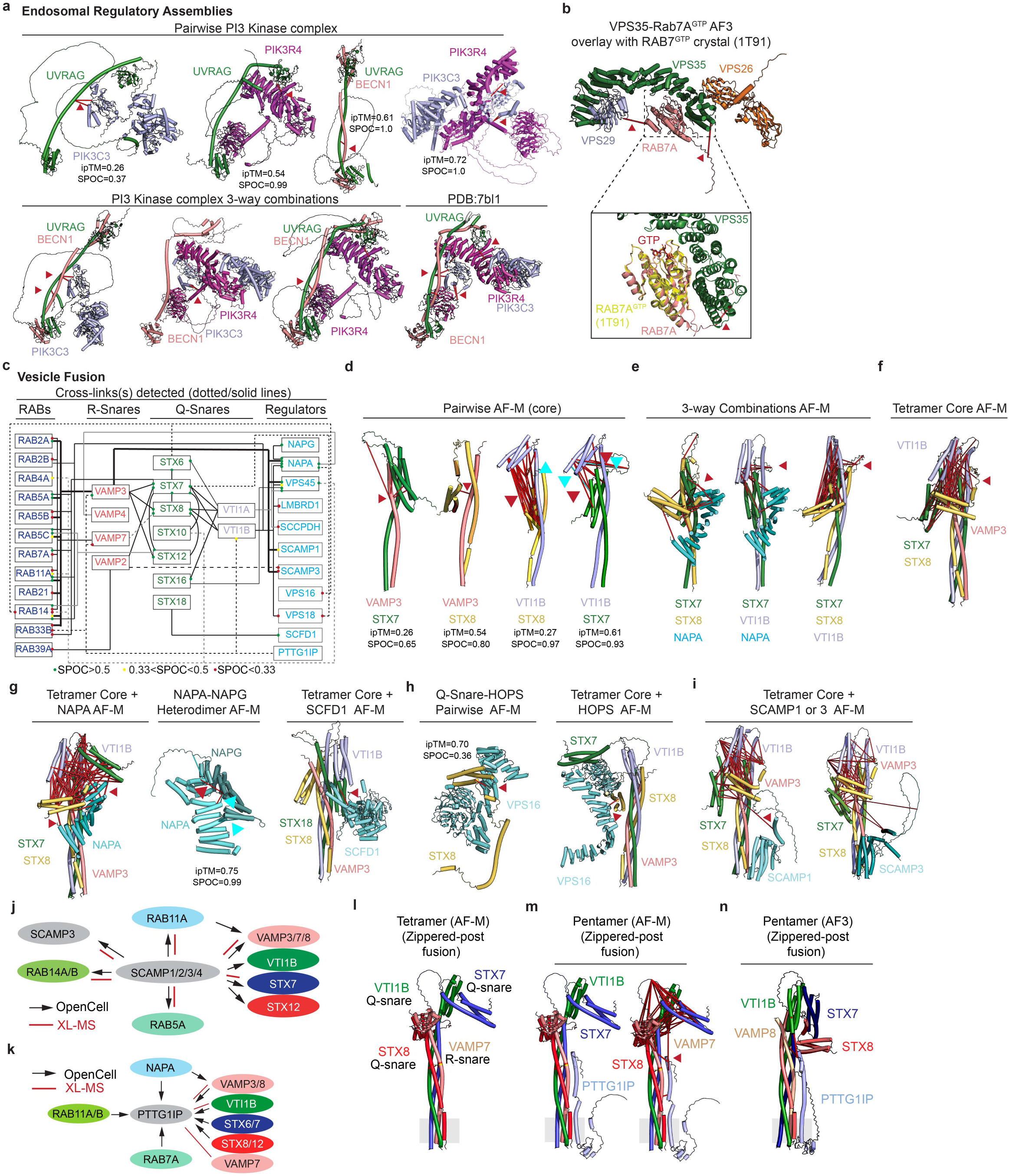
3-way Clique and higher order AF-M predictions reveal extensive SNARE interactions and assemblies. **a**, Pairwise (top) and 3-way clique (bottom) AF-M predictions and associated DSSO cross-links for components of the Class II PI3K complex. Identified cross-links are also mapped onto the cryo-EM structure of the PI3K complex (PDB:7bl1) (lower right). **b**, Overlay of AF3 predictions of a VPS29-VPS35-VPS26A-RAB7A^GTP^ complex and associated DSSO cross-links with a RAB7A^GTP^ crystal structure (PDB:1T91). **c**, Summary of cross-link and AF-M predictions for SNARE components and their interactors. R-SNARE, Q-SNARE, known and candidate regulators and RAB proteins found with cross-links within EndoMAP.v1 are shown. Lines indicate one or more cross-links and are shown in distinct forms to facilitate visualization of connections. Colored dots indicate SPOC score for each AF-M pairwise prediction. **d**, Cross-links and pairwise AF-M predictions for “core” SNARE components VAMP3, STX7, STX8, and VTI1B. ipTM and SPOC scores are indicated for pairwise combinations. **e**, Examples of a subset of 3-way clique predictions and associated cross-links involving core SNARE components as well as NAPA. **f**, Core SNARE AF-M predictions and associated cross-links. The prediction resembles a post-vesical fusion-like conformation. **g**, AF-M predictions and associated cross-links for SNARE association with soluble fusions factors. **h**, Predicted interactions and cross-links for association of VPS16 with either STX8 or STX8 in the core SNARE complex. **i**, Core SNARE assembly predictions and associated cross-links with candidate interactors SCAMP1 and SCFD1. **j**, Summary of physical interactions involving SCAMP proteins in OpenCell^29^ and cross-links identified in our study. Intraprotein cross-links are not shown (see **Supplementary Table 2**). **k**, Summary of physical interactions involving PTTG1IP proteins in OpenCell^29^ and cross-links identified in our study. **l**, AF-M prediction for tetrameric SNARE complex composed of VTI1B, STX7, STX8, and VAMP7. **m**, Pentameric prediction for VTI1B, STX7, STX8, and VAMP7 together with PTTG1IP. Grey rectangles represent the transmembrane section of the complex. Left, cross-links not shown; Right, cross-links shown. **n**, Pentameric AF3 prediction for VTI1B, STX7, STX8, and VAMP8 together with PTTG1IP. DSSO and DHSO/DMTMM interprotein cross-links are indicated with red and cyan lines and arrowheads, respectively.

**Extended Data Fig. 10.**
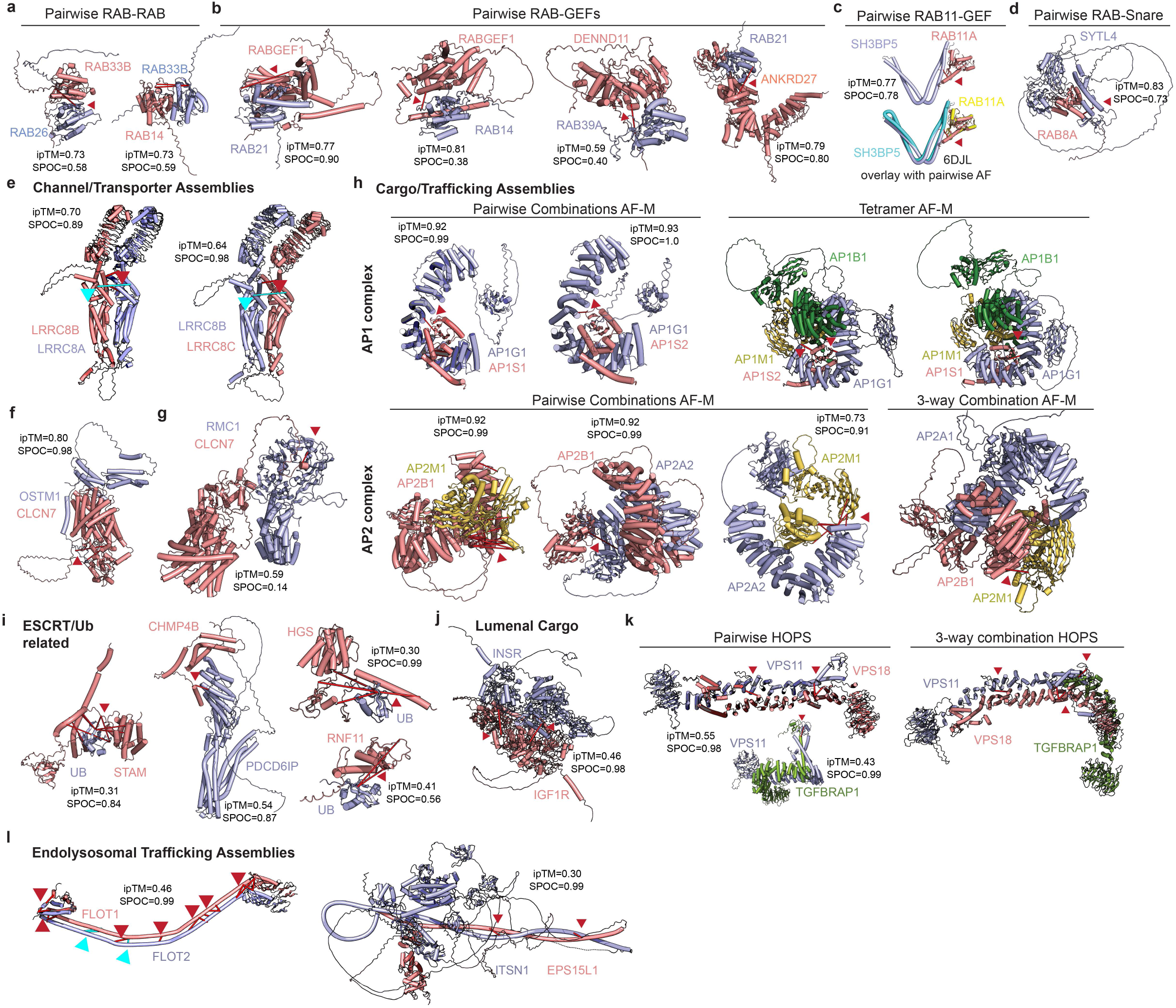
Endosomal Regulatory Proteins, Channels, Cargo, and Trafficking Complex AF-M Predictions. **a**, Pairwise AF-M predictions and associated cross-links for two pairs of RABs with high scoring predictions. **b,c**, Pairwise AF-M predictions and associated cross-links for selected RABGEF complexes present in EndoMAP.v1 (panel **b**), and for a RAB11A-SH3BP5 complex (panel **c**) overlayed with a previously determined structure of the complex (PDB:6DJL). **d**, Pairwise AF-M prediction and associated cross-links for a RAB8A-SYTL4 (synaptotagmin-like) Snare complex. **e-g**, AF-M predictions and associated cross-links for selected channel/transporter assemblies in EndoMAP.v1. LRRC8 proteins (panel **e**) form hexamers and are components of volume regulated anion channels important for cell volume homeostasis. OSTM1-CLCN7 (panel **f**) is an endolysosomal voltage-gated channel mediating exchange of chloride against protons and is known to form a heterotetramer. CLCN7 was found cross-linked to RMC1 (panel **g**), a subunit of the CCZ1-MON1 GEF for RAB7 on endolysosomes. **h**, Pairwise AF-M predictions for cross-link containing AP1 components AP1G1 and either AP1S1 or AP1S2 (left) and tetramer AF-M prediction for AP1G1-AP1B1-AP1M1 and either AP1S1 or AP1S2 (upper panel). Pairwise and 3-way clique predictions for AP2 components AP2M1, AP2B1, or AP2A2 (lower panel). **i**, Pairwise predictions and associated cross-links for ESCRT and ubiquitin (Ub)-related modules within EndoMAP.v1. **j**, Pairwise predictions and associated cross-links for INSR and IGF1R. **k**, Pairwise AF-M predictions and associated cross-links for selected HOPS complex components (left) and a 3-way clique prediction (right) that maintains compatible cross-link distances. **l**, Pairwise predictions and associated cross-links for the FLOT1/2 complex that participates as a scaffolding protein within caveolar membranes, and the ITSN1-EPS15L complex that links endosomal membrane trafficking with actin assembly machinery. For all panels, DSSO and DHSO/DMTMM interprotein cross-links are indicated with red and cyan lines and arrowheads, respectively. Intraprotein cross-links are not shown (see **Supplementary Table 2**).

**Extended Data Fig. 11.**
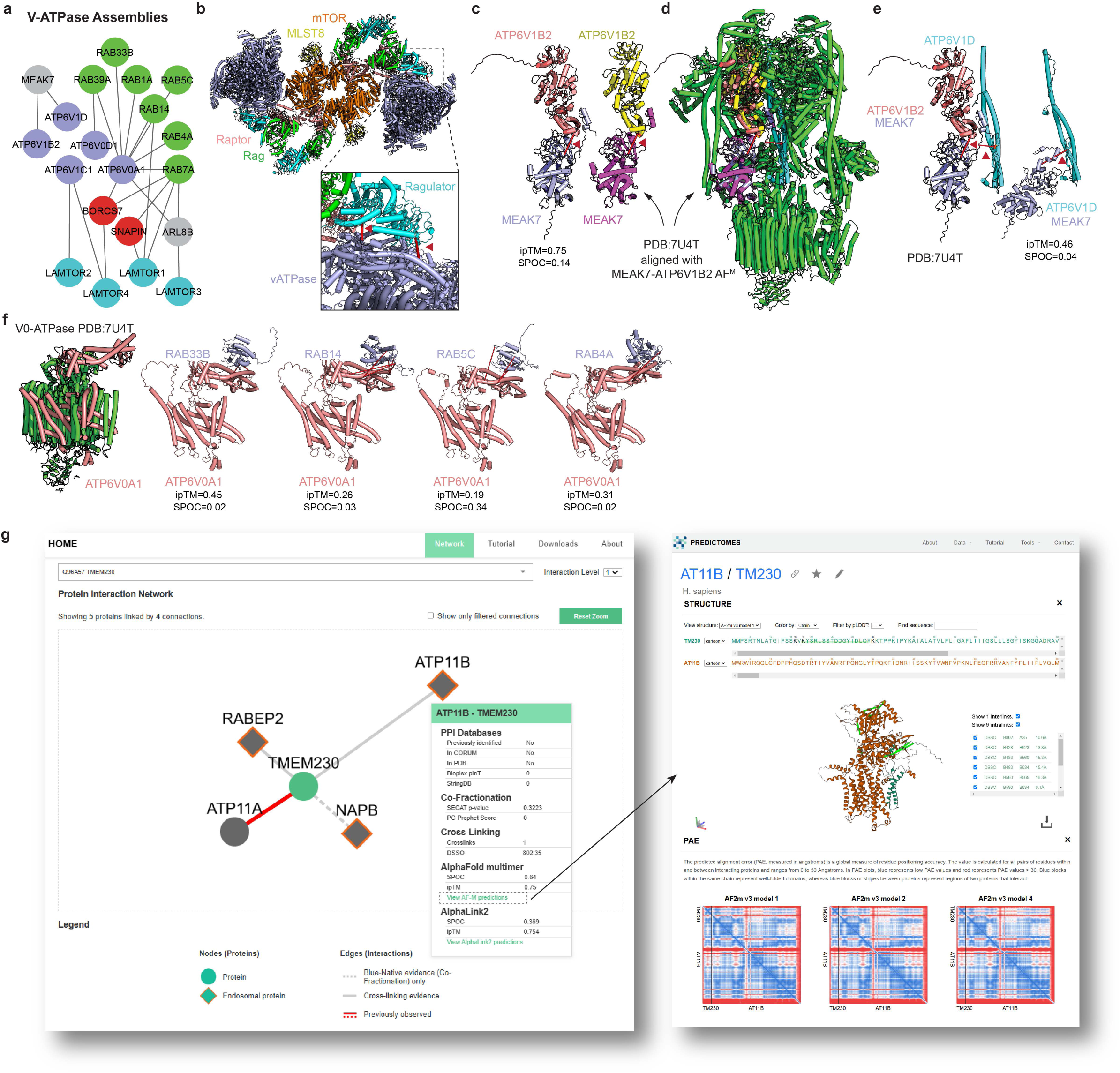
V-ATPase as an interaction hub. **a**, DSSO cross-links identified between components of the V-ATPase (purple), the BORC complex (red), the LAMTOR complex (blue) and RAB proteins (green). **b**, Ribbon diagram of a model for a V-ATPase-MTORC1 supercomplex showing the location of two cross-links identified between V-ATPase and LAMTOR subunits. **c**, Pairwise AF-M prediction and associated cross-link for MEAK7 and ATP6V1B2 (left) is compared with the MEAK7-ATP6V1B2 sub-complex from PDB:7U4T (right)^64^. **d**, MEAK7-V1-ATPase (PDB:7U4T)^64^ together with overlay of AF-M prediction for MEAK7-ATP6V1B2. Cross-links between MEAK7 and either ATP6V1B2 or ATP6V1D are shown by red lines. **e**, MEAK7-ATP6V1B2 AF-M prediction modeled on PDB:7U4T and identified cross-links with ATP6V1D (red arrowhead) is shown on the left. MEAK7-ATP6V1D AF-M prediction and associated cross-link is shown on the right. **f**, Endosomal RABs cross-link with the ATPV0A1 subunit of the V0-ATPase. The structure of the V0-ATPase complex, with ATP6V0A1 shown in salmon, is presented on the far left. The AF-M predictions for 4 endosomal RABs and the detected cross-links are also shown. Intraprotein cross-links are not shown (see **Supplementary Table 2**). **g**, Screenshot from our EndoMAP.v1 website and AF-M prediction viewer at https://endomap.hms.harvard.edu/. Left panel shows the output of a search for TMEM230. Right panel shows the output of AF-M prediction for TMEM230-ATP11B interaction, together with predicted alignment error plots.

## METHODS

### Reagents

The following chemicals and reagents were used: Dounce homogenizer (DWK Life Sciences, 885302-0002); Pierce Anti-HA Magnetic Beads (Thermo Scientific, 88837); Pierce Anti-FLAG Magnetic Agarose (Thermo Scientific, A36797); Anti-FLAG M2 Magnetic Beads (Sigma Millipore, M8823); Pierce Protein A/G magnetic beads (Thermo Scientific, 88802); IGEPAL CA-630 (Sigma-Aldrich, I8896); S-Trap micro columns (Protifi, C02-micro-80); Triethylammonium bicarbonate buffer (TEAB, Sigma-Aldrich, T7408); sodium dodecyl sulfate (SDS; Bio-Rad, 1610302); disuccinimidyl sulfoxide (DSSO; Thermo Scientific, A33545); 3,3’-sulfinyldi(propanehydrazide) (DHSO, CF Plus Chemicals, PCL042); 4-(4,6-dimethoxy-1,3,5-triazin-2-yl)-4-methylmorpholinium chloride (DMTMM, Sigma-Aldrich, 74104); n-Dodecyl β-D-maltoside (DDM, Gold Biotechnologies, DDM5); NativeMark Protein Standard (Invitrogen, LC0725); NativePAGE 4-6% Gels (Invitrogen, BN1002BOX); MultiScreen Filter Plates (Sigma Millipore, MSHVN4510); TMTpro 16plex Set (Thermo Fisher Scientific, A44520); protease inhibitor cocktail (Roche, 4906845001); tris(2-carboxyethyl)phosphine (TCEP; Gold Biotechnology, 51805-45-9); 2-chloroacetamide (Sigma-Aldrich, C0267); *S*-Methyl thiomethanesulfonate (MMTS, Sigma-Aldrich, 208795); trypsin (Promega, V511C); Lys-C (Wako Chemicals, 129-02541); hydroxylamine solution (Sigma-Aldrich, 438227), Sep-Pak C18 and C8 50mg Cartridge (Waters, WAT054955 and WAT054965), High-pH Reversed-Phase Peptide Fractionation Kit (Thermo Scientific, 84868), Bio-Rad Protein Assay Dye (Bio-Rad, 5000006); EPPS (Thermo Scientific, J61296AE); Empore SPE Disks C18 (Sigma Millipore, 66883-U); Gateway LR Clonase II Enzyme Mix (Thermo Scientific, 11791020); NEBNext Ultra II Q5 Master Mix (New England BioLabs, M0544L); Cas9-NLS (QB3 MacroLab at Berkeley), CloneR (StemCell Technologies, 05889); MiSeq Reagent Nano Kit v2 (300 cycles; Illumina, MS-103-1001); GeneArt Precision gRNA synthesis kit (Thermo Fisher Scientific, A29377); RNAeasy Qiagen kit (Qiagen, 74104); 24 Well glass bottom plates (Cellvis, P24-1.5H-N); Corning square culture dish (Corning, 431110); Nunc Nunclon Delta cell culture dishes (Thermo Scientific, 140675, 150318 and 168381); Corning Matrigel Matrix (Corning, 354230); DMEM/F12 (Gibco, 11330057); neurobasal medium (Thermo Scientific, 21103049); non-essential amino acids (NEAAs, Gibco, 11140050); GlutaMAX (Gibco, 35050061); N-2 supplement (Gibco, 17502048); neurotrophin-3 (NT3; Peprotech, 450-03); brain-derived neurotrophic factor (BDNF; Peprotech, 450-02); B27 (Gibco, 17504001); Y27632 dihydrochloride (ROCK inhibitor; PeproTech, 1293823); Cultrex 3D culture matrix laminin I (R&D Systems, 3446-005-01); accutase (StemCell Technologies, 07922); FGF2-G3 (in-house); human insulin (Santa Cruz Biotechnologies, sc-360248); transforming growth factor-β (PeproTech, 100-21); holo-transferrin human (Sigma-Aldrich, T0665); sodium bicarbonate (Sigma-Aldrich, S5761-500G); sodium selenite (Sigma-Aldrich, S5261-10G); doxycycline (Clontech Labs, 631311); UltraPure 0.5 M EDTA (Invitrogen, 15575020); 16% paraformaldehyde (Electron Microscopy Science, 15710); DMEM (Gibco, 11995073); fetal bovine serum (Cytiva, SH30910.03); Hydrocortisone (Sigma-Aldrich, H0135); Polyethylenimine (Polysciences, 23966); FuGENE (Promega, E2311).

The following primary antibodies were used (1:1,000 for immunoblotting, 1:400 for immunofluorescence): FLAG (Sigma-Aldrich, F1804), HA (Cell Signaling Technology, 3724), V5 (Invitrogen, 14-6796-82), TMEM230 (Origene, TA504888), LAMP1 (Cell Signaling Technology, D2D11), RAB5 (Cell Signaling Technology, C8B1), CLR (ProteinTech, 10292-1-AP), golgin 97 (ProteinTech, 12640-1-AP), VDAC1 (ProteinTech, 55259-1-AP), CLCN3 (Cell Signaling Technology, 13359S), GFP (Thermo Scientific, a10262), mCherry (Thermo Scientific, M11217), EEA1 (Cell Signaling Technology, C45B10). The following secondary antibodies were used (1:10,000 for immunoblotting, 1:400 for immunofluorescence): anti-rabbit immunoglobulin-G (IgG) horse radish peroxidase (HRP) conjugate (BioRad, 1706515); anti-mouse IgG HRP conjugate (BioRad, 1706516); Goat anti-Chicken IgY (H+L), Alexa Fluor 488 (Thermo Scientific, A-11039); Goat anti-Rat IgG (H+L) Cross-Adsorbed, Alexa Fluor 555 (Thermo Scientific, A-21434); Goat anti-Rabbit IgG (H+L) Cross-Adsorbed, Alexa Fluor 647 (Thermo Scientific, A-21244).

### Molecular cloning

Plasmids were made as previously described (https://dx.doi.org/10.17504/protocols.io.5jyl8p7n8g2w/v1). Entry clones from the human ORFeome collection, version 8, were cloned into their corresponding plasmids using Gateway technology (Thermo Fisher Scientific) or Gibson assembly (New England Biolabs). The complete TMEM230 triplemutant was obtained by gene synthesis (Twist Bioscience). For lentivirus transduction, pHAGE and pLenti backbones were used. For transfection, pCGS and pcDNA3.1backbones were used. The following plasmids were generated: pGCS-3xFlag-ATP11B (Addgene, 225511), pcDNA-TMEM30A-V5 (Addgene, 225510), pGCS-3xHA-TMEM230 (Addgene, 225512), pGCS-3xHA-TMEM230triplemut (Y29C, R78L, X121W, Addgene 225513), plenti-UBC-HA-TMEM230 (Addgene, 225516), plenti-UBC-HA-TMEM230(R78L) (Addgene, 225517), plenti-UBC-HA-TMEM230(X121W) (Addgene, 225519), plenti-UBC-HA-TMEM230(Y29C) (Addgene, 225520), plenti-UBC-HA-TMEM230(X121PG) (Addgene, 225518), plenti-UBC-HA-TMEM230triplemut(Y29C, R78L, X121W, Addgene 225521), pcDNA-CLCN3-3xFlag (Addgene, 225506), pcDNA-CLCN5-3xFlag (Addgene, 225507), pcDNA-TMEM9B-3xHA (Addgene, 225509), pcDNA-TMEM9-3xHA (Addgene, 225508), pHAGE-mCherry-CLCN3 (Addgene, 225514), pHAGE-TMEM9-EGFP (Addgene, 225515). The following plasmids were used for lentiviral packaging: pPAX2 (Addgene, 12259), pMD2 (Addgene, 12260).

### Cell culture, neuronal differentiation and lentiviral transduction

HEK293 cells (ATCC; RRID:CVCL_0045) were cultured in 10cm dishes with high-glucose and pyruvate DMEM supplemented with 10% fetal bovine serum. For co-immunoprecipitation experiments, cells were transfected at 60% confluency with 6μg of plasmids in a 2:1 ratio using polyethylenimine (25 kDa) and incubated for 48h at 37°C and 5% CO_2_.

SUM159PT cells (a gift from Tobias Walter (Memorial Sloan Kettering); RRID:CVCL_5423) were cultured into 6-well culture dishes (300 000 cells/well) with DMEM/F12 supplemented with GlutaMAX, 5% fetal bovine serum, 1μg/ml hydrocortisone and 5μg/ml insulin. Cells were transfected one day later with 500ng of plasmids using FuGENE and Optimem transfection reagent and incubated at 37°C and 5% CO_2_. One day after transfection, cells were selected with puromycin and plated into 24-well glass bottom culture dishes (50 000-100 000 cells/well).

Gene edited human embryonic stem cells (hESCs, H9, WiCell Institute) were cultured as described (https://dx.doi.org/10.17504/protocols.io.j8nlkoq56v5r/v1) ^68^. Cells were maintained with E8 medium on plates coated with Matrigel and split with 0.5 mM EDTA in DPBS.

hESCs with the AAVS1-TRE3G-NGN2 driver^69^ were differentiated into iNeurons as described (https://dx.doi.org/10.17504/protocols.io.x54v9p8b4g3e/v1). Briefly, stem cells were plated at 2 x 10^5^ cells/mL (differentiation day 0) in ND1 medium (DMEM/F12, N2, human 10ng/mL BDNF, 10 ng/mL human neurotrophin-3 (NT3), NEAA, 0.2μg/mL human laminin) supplemented with 2mg/mL doxycycline and 10 μM Y27632 (ROCK inhibitor). The next day the medium was exchanged with ND1 without Y27632. The following day, the medium was replaced with ND2 (neurobasal medium, B27, GlutaMAX, 10ng/mL BDNF, 10ng/mL NT3) supplemented with 2μg/mL doxycycline. Until the experimental day (day 19-21), 50% of the medium was replaced with fresh ND2 every other day. Cells were replated at 4 x 10^5^ cells per well on day 4-6. From day 10, doxycycline was removed from the ND2.

Lentiviral vectors were packed in HEK293T cells (ATCC #CRL-3216; RRID:CVCL_0045) as described (https://dx.doi.org/10.17504/protocols.io.6qpvr3en3vmk/v1)^68,70^. Cells were co-transfected at 60% confluency with pPAX2, pMD2 and the target vector in a 4:2:1 ratio using polyethylenimine. The medium was changed to ND2 the next day and collected two days after transfection. ND2 medium containing lentivirus was filtered (0.22μm) and used for transduction of iNeurons at differentiation day 11-12.

### CRISPR–Cas9 gene editing

hESCs (H9 AAVS1-TRE3G-NGN2 3xFLAG-EEA1; RRID:CVCL_D1KV) were gene edited using CRISPR–Cas9^71^. Cells were electroporated with a mixture of 0.6μg guide RNA (sgRNA) and 3μg Cas9-NLS (QB3 MacroLab, UC Berkeley) using a Neon transfection system as previously described (https://dx.doi.org/10.17504/protocols.io.ewov1qykkgr2/v1) according the the specific protocol (DOI: dx.doi.org/10.17504/protocols.io.3byl49d82go5/v1). To generate hESCs homozygous for TMEM230^X121W^ variant, a ssDNA oligo was included in the electroporation (CTACCGTGGTTACTCCTATGATGACATTCCAGACTTTGATGACTGGCACCCACCCCATAGCTGAGGAGGAGTCACAGTGGAACTGTCCCAGCTTTAAGATATCTAGCAGAAACTATAGCTG). The cells were recovered for 24-48h in a low O_2_ incubator and sorted into single cells with a Sony Biotechnology (SH800S) cell sorter. Gene editing of individual clones was verified by sequencing with Illumina MiSeq system and validated by immunoblotting and/or mass spectrometry. Guide RNAs (sgRNAs) were generated using the GeneArt Precision gRNA synthesis kit (ThermoFisher Scientific) for the sequences: TMEM230^-/-^ CCTGAAGGTCAATGTAGCCATCGT, TMEM230^X121W^ CTCCTCCTCAGCTATGGGGT, TMEM9^-/-^ TATCTTTGGTGGCTGTGGTC, TMEM9B^-/-^ TCTACATCAGGCCCCCGCAC.

### Spinning disk confocal microscopy

For immunofluorescence staining, SUM159PT cells were fixed with 4% paraformaldehyde in PBS for 15min and permeabilized with 0.5% Triton X-100 in PBS for 10min at room temperature. Cells were blocked with 3 % BSA in PBS with 0.1% Triton X-100 for 1h at room temperature. Cells were incubated with primary antibodies (1:200 dilution) in 3 % BSA in PBS with 0.1% Triton X-100 for 3h at 4°C. After washes, cells were incubated with Alexa Fluor secondary antibodies (1:400) for 1h at 4°C and nuclei were stained with Hoechst33342 (1:10000) for 5 min. Cells were washed and maintained in PBS until microscopy analysis. Immunostaining of iNeurons was performed according to the protocol: dx.doi.org/10.17504/protocols.io.kqdg32zrpv25/v1.

Cells were imaged using a Yokogawa CSU-X1 spinning disk confocal on a Nikon Eclipse Ti-E motorized microscope and a Plan Apochromat 100×/1.45 N.A oil-objective lens. Live-cell imaging was performed with a Tokai Hit stage top incubator at 37°C, 5 % CO_2_ and 95 % humidity. Images were acquired with a Hamamatsu ORCA-Fusion BT CMOS camera (6.5 μm^2^ photodiode, 16-bit) and NIS-Elements image acquisition software. All samples were measured under the same exposure time and laser power. Colocalization analysis was performed with JACoP plugin for ImageJ/FiJi^72^ using maximum intensity projection images and maximum entropy threshold. Linear mixed-effects models statistics were applied as implemented in *lme4* R package with a nested design to account for images acquired from the same culture well and same biological replicate. The number of fields of view for each of the three independent biological replicates is indicated in the figures (**Fig. 4e,f**).

### Meta-analysis

Meta-analysis was performed to define the endosomal proteome and assign an unbiased score to each protein reflecting the probability of being located in endosomes based on experimental data. The literature was surveyed for studies capturing the endosomal proteome in mammalian organisms, which resulted in 16 datasets (**Supplementary Table 1**)^24,25,73-82^. Incomplete datasets or with ambiguous organelle purifications (e.g. “vesicles” or mixed organelles) were excluded. Outdated Uniprot IDs and obsolete gene names were updated (Uniprot 2022-02). Ensembl and *BiomaRt* R package were used to retrieve and match rodent genes to their human orthologs, including all human genes when multiple matched. Subsequent analyses were based on the protein identification across datasets as metric for the meta-analysis (**Supplementary Table 1**). To evaluate the performance of meta-analysis metrics and datasets, a reference list of 292 well-known endosomal proteins was manually curated from published literature (**Extended Data Fig. 1a**)^2,4,30,83-117^. Datasets overview was visualized by multiple correspondence analysis (MCA) using *FactoMineR* R package (**Extended Data Fig. 1b**). Protein annotation to various organellar locations was based on a previous study^68^ (**Extended Data Fig. 1c**). Another metric of the meta-analysis was the protein abundance in Endo-IP obtained from the label-free proteomic analysis of endosomal pellets as described below (**Extended Data Fig. 1d, Supplementary Table 1**). Number of interactions with endosomal proteins was obtained from BioPlex 3.0, STRINGDB and CORUM (28.11.2022 Corum 4.1 release)^28,118,119^ for the reference list of well-known endosomal proteins described above. For STRINGDB, only physical interactions with experimental evidence or databases with high score (combined score >0.7) were included.

The performance of each metric to classify endosomal proteins (from the reference list described above) was evaluated by receiver operating characteristic (ROC) curves using *pROC* R package with a binomial logistic regression as predictor (**Fig. 1b**). Combined meta-analysis score was obtained by summation of the three individual metrics. Partial area under the curve (AUC) and the threshold to consider a protein as endosomal was 10% false positives based on the reference list. Meta-analysis resulted in 407 predicted endosomal proteins (14 proteins present in MitoCarta 3.0 were excluded) that were combined with the reference list of well-known endosomal protein for a total of 522 proteins (**Supplementary Table 1**). These proteins were characterized using BioPlex 3.0^28^, OpenCell^29^, and publications as retrieved from Uniprot (**Extended Data Fig. 1e-g**). Endosomal annotation for all subsequent analyses was based on this list.

### EEA1-positive endosome purification through Endo-IP affinity capture

Endo-IPs with HEK293^EL^ cells were performed as described (https://doi.org/10.17504/protocols.io.ewov14pjyvr2/v2). HEK293^EL^ cells expressing FLAG-EEA1 ^25^ were harvested from five 24.5cm square culture dishes per replicate for co-fractionation experiments (n=3) and 60 square plates per replicate (divided in two batches) for cross-linking experiments (n=2). Endo-IPs in iNeurons were performed as describe (dx.doi.org/10.17504/protocols.io.kqdg32zoev25/v1)^51^. Three 15cm culture dishes per replicate were used for experiments in iNeurons (n=3). Cells were pelleted at 1,000 x *g* for 2min at 4°C and washed once with KPBS buffer (100mM potassium phosphate, 25mM KCl and protease inhibitor cocktail, pH 7.2). Cell pellets were resuspended in KPBS and lysed in a Dounce homogenizer with 25 strokes. Samples were centrifuged twice at 1,000 x *g* for 5min at 4 °C and postnuclear supernatants (PNS) protein concentration was quantified normalized by Bradford assay. Samples were incubated for 50min at 4°C with 70μL α-FLAG Sigma magnetic beads for iNeurons experiments, 1.6mL Sigma α-FLAG Sigma magnetic beads for co-fractionation experiments and 20mL of α-FLAG Pierce magnetic beads per batch for cross-linking experiments. The beads were washed four times using a magnetic stand with KPBS. For quantitative proteomics, endosomes were eluted with 120µL 0.5% NP40 (IGEPAL) in KBPS for 30 min at 4°C and stored at −80°C until MS sample preparation. For co-fractionation and cross-linking experiments, endosomes were eluted twice with 0.8mM 3XFlag peptide in KPBS for 45 min at 4°C (**Extended Data Fig. 1h**). Peptide-eluted samples were centrifuged for 20min at 10,000 x *g* in Posi-Click tubes (Denville, c2170). Endosomal pellets were washed twice with KPBS to remove excess of 3XFlag peptide and immediately processed. An additional wash was performed for the second replicate of cross-linking experiment, which helped increase the coverage in the MS analysis.

### Protein co-immunoprecipitation (IP)

A protocol for this analysis is provided at: dx.doi.org/10.17504/protocols.io.n2bvjn5zpgk5/v1. Proteins from a 10cm culture dish of HEK293 cells or a 15cm dish of iNeurons per replicate (n=2 or 4) were extracted for 1h at 4°C with 0.5% n-Dodecyl β-D-maltoside (DDM) in 25mM HEPES pH7.4, 150mM NaCl and protease inhibitor cocktail ^120^. Samples were centrifuged twice at 20,000 x *g* for 20 min and the supernatant was incubated with 15μL α-HA magnetic beads (Pierce) or 25µL α-Flag magnetic beads (Sigma) depending on the protein tag for 2h at 4°C. For IP using endogenous antibodies, the supernatant was incubated overnight with 5μg of antibody prior to the incubation with 15μL magnetic A/G beads. The beads were separated with a magnetic stand and washed four times with washing buffer (0.1% DDM, 25 mM HEPES, 150 mM NaCl, pH 7.4). Proteins bound to the beads were eluted with 30μL 1.5X Laemmli buffer for immunoblotting or 30μL 1.5X S-Trap lysis buffer (7.5% SDS, 150mM Triethylammonium bicarbonate or TEAB pH 8.5) for MS analysis and heated at 80°C for 5 min.

### SDS-PAGE immunoblotting

Samples mixed with Laemmli buffer were incubated at 80°C for 5 min and loaded in a Criterion TGX Stain-free Precast gel for subsequent immunoblotting. After electrophoresis, gels were scanned using a BioRad ChemiDoc imager (Bio-Rad) and electro-transferred onto a PVDF membrane overnight at 10V. Membranes were blocked with 5% non-fat milk, incubated with primary antibody for 2h at 4°C and subsequently with HRP-conjugated secondary antibodies for 1h at 4°C. After washing, blot images were acquired in a BioRad ChemiDoc imager using SuperSignal West Pico PLUS Chemiluminescence substrate (ThermoFisher, catalog number 34580). Images were processed with BioRad Image Lab software (version 6.1.0). Differences in loading were normalized using the stain-free quantification of total protein amount. Protocols for this procedure can be found at: dx.doi.org/10.17504/protocols.io.q26g71eq9gwz/v1.

### Blue Native Polyacrylamide Gel Electrophoresis (BN-PAGE) co-fractionation and in-gel digestion

A detailed protocol for this procedure can be found at: DOI: dx.doi.org/10.17504/protocols.io.81wgbzm2ygpk/v1. Protein complexes from 3 independent biological Endo-IP replicates were fractionated and processed as previously described^121^. Freshly prepared purified endosomal pellets were resuspended in 40μL KPBS with 0.5% n-Dodecyl β-D-maltoside (DDM) and proteins were extracted for 45min at 4°C in rotation. Protein extracts were clarified by centrifugation at 20,000 x *g* and mixed with 10μL BN loading buffer, 1μL Coomassie G-250 mix and 0.5μL native molecular weight marker. Samples were run on a 4–16% NativePAGE gel at 150 V for 1.5h and at 250V for 20min at 4 °C. Gels were fixed in 50% ethanol and 3% phosphoric acid, followed by staining with Coomassie. Each sample was cut into 48 1-mm slices and transferred to a 96-well filter plate for in-gel digestion ^120^. Briefly, proteins were reduced with 100µL 5mM tris(2-carboxyethyl)phosphine (TCEP) in 50mM ammonium bicarbonate for 30 min at 37 °C. Proteins were alkylated with 20mM chloroacetamide in 50 mM ammonium bicarbonate for 15 min at room temperature. Fractions were destained, dried and digested with 0.2μg Lys-C for 4h at 37 °C followed by overnight incubation with 0.2μg of trypsin. Peptides were extracted, dried in a SpeedVac and reconstituted in 5% ACN, 5% formic acid for DIA LC-MS/MS analysis.

### Cross-linking and strong cation exchange (SCX) fractionation

A detailed protocol for both cross-linking procedures can be found at: dx.doi.org/10.17504/protocols.io.261ge5q2og47/v1. Freshly prepared purified endosomal pellets from 2 independent biological replicates were resuspended in 300μL KPBS and immediately cross-linked by incubating with 1 mM DSSO (Bis(2,5-dioxopyrrolidin-1-yl) 3,3′-sulfinyldipropionate, Bis-(propionic acid NHS ester)-sulfoxide, Thermo Fisher Scientific) at room temperature for 40min^7^. The reaction was quenched with 50mM Tris buffer pH 7.5 at room temperature for 30 min. Cross-linked samples were denatured in 8M urea, reduced with 5mM dithiothreitol (DTT) for 30 min at 37°C, and alkylated with 40mM chloroacetamide for 30 min at room temperature. Cross-linked proteins were digested with Lys-C (1:75) at 37 °C overnight. Sample urea concentration was diluted to 2M with 50mM EPPS (3-[4-(2-Hydroxyethyl)-1-piperazine]propanesulfonic acid) and incubated at 37 °C with trypsin (1:100) for 6h. Peptides were desalted with a 50mg C8 Sep-Pak solid phase extraction column, dried and fractionated by strong cation exchange (SCX) chromatography. A 70-min linear gradient of mobile phase (0.5M NaCl in 20% ACN, 0.05% formic acid) was used from 0 to 8% in 14min, to 20% at 28min, 40% at 48min and to 90% at 68min at a column flow rate of 0.18mL/min in a PolyLC PolySulfoethyl A column (3μm particle size, 2.1mm inner diameter and 100mm length). Fractions were collected every 30s starting at 35min for 10min, and then every minute. Fractions were dried in a SpeedVac and desalted using C8 StageTip. Around 30 fractions for each sample were reconstituted in 5% ACN, 5% formic acid and analyzed by LC-MS/MS.

An additional independent biological replicate of freshly prepared purified endosomal pellet was resuspended in 300μL KPBS and immediately cross-linked by incubating with a combination of 8 mM DHSO (3,3’-sulfinyldi(propanehydrazide)) and 16mM DMTMM (4-(4,6-dimethoxy-1,3,5-triazin-2-yl)-4-methylmorpholinium chloride) at 37°C for 90 min ^39^. Cross-linked samples were denatured in 5% SDS and briefly sonicated, reduced with 5mM dithiothreitol (DTT) for 5 min at 55°C, and alkylated with 20mM MMTS (Methyl methanethiosulfonate). Cross-linked proteins were precipitated and subjected to S-Trap™ mini-spin column digestion protocol as provided by the manufacturer (see below). Peptides were desalted and fractionated by SCX chromatography as described above. A total of 30 fractions were analyzed by LC-MS/MS.

### S-Trap sample preparation

Three independent replicates of PNS samples (10μg or 50μg of protein depending on the experiment) and Endo-IP samples were mixed with equal volume of water and subjected to sample preparation. S-Trap™ micro-spin column digestion protocol (version 4.7) was followed as provided by the manufacturer (Protifi, C02-micro-80)^122,123^ (see: dx.doi.org/10.17504/protocols.io.bys6pwhe). Briefly, each sample was mixed with equal volumes of 2X lysis buffer (10% SDS, 100mM Triethylammonium bicarbonate or TEAB pH 8.5). Protein IP samples from iNeurons (n=2 or 4) were directly collected in 1.5X lysis buffer. Proteins were reduced by incubating at 55°C for 30min with 5mM tris(2-carboxyethyl) phosphine (TCEP) and alkylated for 30min at room temperature with 40mM chloroacetamide. Samples were acidified with phosphoric acid and mixed with washing buffer (90% methanol, 100 mM TEAB pH 7.55). Samples were transferred to micro-spin columns and washed 4 times with 150µL washing buffer by centrifugation. Proteins were digested with 0.5µg Lys-C at 37 °C overnight in a humid chamber, followed by 6h incubation with 0.5μg of trypsin. Peptides were collected from the column by three subsequent centrifugation steps (with 50mM TEAB, 0.2% formic acid and 50% acetonitrile or ACN, respectively) and dried in a SpeedVac.

### TMT labeling and peptide fractionation

Protocols for labeling of peptides are provided at: DOI: dx.doi.org/10.17504/protocols.io.rm7vzjej4lx1/v1. Peptides were resuspended in 50μL (PNS samples) or 35μL (Endo-IP samples) 100mM TEAB pH 8.5. PNS and Endo-IP peptides were labelled by adding 11μL or 7μL ACN, and incubating 1h at room temperature with 10μL or 8μL of TMTpro reagent (20 mg/mL stock in ACN), respectively. The reaction was quenched by adding 10μL 5% hydroxylamine for 15min.

For PNS samples, equal peptide amounts for each sample were combined, desalted with a 100mg C18 Sep-Pak solid phase extraction column and fractionated by basic pH reversed-phase HPLC. Chromatography was performed with a 50-min linear gradient from 5% to 35% ACN in 10mM ammonium bicarbonate pH 8 at a column flow rate of 0.25mL/min using an Agilent 300 Extend C18 column (3.5μm particle size, 2.1mm inner diameter and 250mm length). The initial 96 fractions collected were combined into 24 fractions, as described previously^124^. One set of 12 non-adjacent fractions were dried in a SpeedVac and desalted using C18 StageTip. Dried peptides were reconstituted in 5% ACN, 5% formic acid and subjected to LC-MS/MS analysis.

For Endo-IP samples, equal peptide amounts for each sample were combined and fractionated using High-pH Reversed-Phase Peptide Fractionation Kit (Pierce) following the manufacturer’s protocol. Eluates were combined into 6 fractions, dried and desalted using C18 StageTip. Dried peptides were reconstituted in 5% ACN, 5% formic acid and subjected to LC-MS/MS analysis.

For protein IP samples, equal peptide amounts for each sample were combined, dried and desalted using C18 StageTip without further fractionation. Dried peptides were reconstituted in 5% ACN, 5% formic acid and subjected to LC-MS/MS analysis.

### Liquid Chromatography-Mass spectrometry (LC-MS) data acquisition

TMT-labeled samples were analyzed using a Vanquish Neo UHPLC system coupled to an Orbitrap Eclipse Tribid mass spectrometer with FAIMSpro. Peptides were separated on a 100μm microcapillary column packed with 20cm of Accucore C18 resin (2.6 μm, 150 Å). A 90-min linear gradient from 5% to 20% ACN in 80min, to 36% at 83min and 98% at 85min in 0.125% formic acid was used at 0.3µL/min. MS1 spectrum was acquired on the Orbitrap (resolution 60,000, scan range 350-1,350 m/z, standard automatic gain control (AGC) target, auto maximum injection time). Peptide fragmentation was achieved by high-energy collisional dissociation (HCD) at 36% normalized collision energy. MS2 was acquired on the Orbitrap (resolution 30,000, isolation window 0.6 m/z, TurboTMT set to All TMT Reagents, first mass 120 m/z, 200% normalized AGC, 120ms maximum injection time). FAIMS Pro was set to -30, -50 and -70 compensation voltage (CV). Unfractionated samples (protein IPs) were injected twice with FAIMS set to - 40, -60 and-80 CV for the second run.

BN-PAGE co-fractionation samples were analyzed using an EASY-nLC 1200 system coupled to an Orbitrap Exploris 480 Mass Spectrometer. A 15cm 100-μm capillary column was packed in-house with Accucore 150 C18 resin (2.6μm, 150 Å). A 90-min linear gradient from 5% to 20% ACN in 80min, to 25% at 83min and 98% at 85min in 0.125% formic acid was used at 0.3µL/min. The data-independent acquisition (DIA) method consisted on MS2 analysis of overlapping isolation windows of 24 m/z stepped through 390- 1,014 m/z mass range for the first cycle and 402-1,026 m/z for the second cycle ^125^. DIA scans were performed with 28% normalized HCD collision energy, 30,000 resolution, 145-1450 m/z scan range, 1000% normalized AGC and 54ms maximum injection time. This was followed by a parent MS1 ion scan (resolution 60,000, scan range 350-1,050 m/z, 100% normalized AGC target, auto maximum injection time).

DSSO cross-linking samples were analyzed using an EASY-nLC 1200 system coupled to an Orbitrap Fusion Lumos mass spectrometer with FAIMSpro. A 90-min linear gradient from 5% to 20% ACN in 80min, to 25% at 83min, to 40% at 85min and 98% for 2min in 0.125% formic acid was used at 0.5µL/min. An HCD-MS2 strategy was used^5^, in which the MS1 spectrum was acquired on the Orbitrap (resolution 120,000, scan range 400-1,600 m/z, standard AGC target, auto maximum injection time). Peptide with charge states 4-8 were fragmented by HCD at 21, 27 and 33% normalized collision energy. MS2 was acquired on the Orbitrap (resolution 60,000, isolation window 1.6 m/z, auto scan range, 200% normalized AGC, 120ms maximum injection time). FAIMS Pro was set to −50, −60 and−75 CV ^126^.

DHSO and DMTMM cross-linking samples were analyzed using a Vanquish Neo UHPLC system coupled to an Orbitrap Ascend MultiOmics Tribid mass spectrometer with FAIMSpro. A 90-min linear gradient from 5% to 20% ACN in 80min, to 25% at 83min, to 40% at 85min and 98% for 2min in 0.125% formic acid was used at 0.3µL/min. MS^1^ spectrum was acquired on the Orbitrap (resolution 120,000, scan range 350-1,600 m/z, standard AGC target, auto maximum injection time). Peptide with charge states 4-8 were fragmented by HCD at 21, 27 and 33% normalized collision energy. MS2 was acquired on the Orbitrap (resolution 60,000, isolation window 1.4 m/z, auto scan range, 200% normalized AGC, 120ms maximum injection time). FAIMS Pro was set to −50, −60 and−75 CV.

### Proteomics data analysis

TMT-MS data was processed with MSconverter^127^ and searched using Comet^128^ against the human canonical proteome (UniProt Swiss-Prot 2021-11), including reverse sequences and common contaminants. Experiments containing disease variant of TMEM230 were search against the human canonical proteome (UniProt Swiss-Prot 2024-01) including an additional sequence of TMEM230 with such variants. Peptide mass tolerance was set to 50ppm and fragment ion tolerance to 0.02 Da. These wide mass tolerance windows were chosen to maximize sensitivity in conjunction with Comet searches and linear discriminant analysis^129^. TMTpro labels were set as fixed modification on lysines and peptide N-terminus (+304.207 Da), carboxyamidomethylation on cysteines (+57.021 Da) as fixed modification, and oxidation on methionine residues as variable modification. Linear discriminant analysis was performed^130^and peptide-spectrum matches (PSMs) were filtered to 2% false discovery rate (FDR)^131^. TMT-reporter ions were quantified by picking the most intense peaks within 0.003 Da around the theoretical m/z, and corrected for isotopic impurity. Only PSMs with at least 200 total signal-to-noise ratio across all TMT channels and 50% precursor isolation purity were used^132^. Data summarization, normalization and statistics were performed using MSstats^133,134^. Peptide level normalization and imputation were enabled, and protein summarization method was set to “LogSum” for Endo-IP experiments from iNeurons and to “msstats” for all other experiments. The threshold used to consider significantly regulated proteins was 0.05 q-value and 1.5 fold-change. For PNS and Endo-IP experiments from iNeurons, 3 biological replicates per condition (**Supplementary Table 4-5**). For protein IP experiments in iNeurons, 4 biological replicates were analyzed per group (**Supplementary Table 5**), except for one dataset with some groups containing two replicates given the limitation of the maximum number of TMT channels (**Supplementary Table 4**). Synaptic Gene Ontology enrichment analysis was performed using SynGO ^135^ (https://www.syngoportal.org/#) using all proteins identified in each experiment as background.

DIA-MS data was analyzed using DIA-NN (version 1.8) as previously described^136,137^. Data was converted to mzML using MSconvert^127^ using demultiplex set to overlap only (10ppm mass error). A spectral library was generated from the complete human proteome (UniProt 2022-05) with a precursor m/z range of 350-1050, precursor charge 2-5 and fragment ion m/z range 145-1450. Carbamidomethylation, oxidation and N-terminal excision were included as modifications. Search was performed with 10ppm mass accuracy, match-between-runs (MBR) enabled and robust LC (high precision) quantification strategy. For Endo-IP protocol optimization samples (**Extended Data Fig. 1i-l, Supplementary Table 1**), downstream analysis was performed using MS-DAP ^138^. Only peptides quantified in all three replicates per condition (n=3) were included. Data was normalized with variance stabilization normalization (Vsn) and mode-between protein methods. DEqMS algorithm was selected for statistical analysis, using a significance threshold of 0.01 FDR-adjusted p-value threshold and log2 fold change of 3 (**Supplementary Table 1**). For BN cofractionation experiments, protein complex analysis was performed with PCprophet ^33^. Three biological replicates were analyzed with default parameters using the provided core complexes as database and BN markers for calibration as mode for collapsing hypothesis to common complexes. As previously described ^8^, co-elution scores (from rf output table) were assigned to each protein pair of the complex and used for downstream analysis. Only complexes with a minimum peak elution at 67kDa fraction and a maximum of 25 proteins per complex were considered. In addition, we only considered interactions with a score of at least 0.7 in two replicates to recover only high confidence candidate interactions (**Supplementary Table 2**). These parameters were selected based on the optimal recovery of protein interactions reported in BioPlex (**Extended Data Fig. 2b,c**) ^8^. Elution profiles and Pearson’s correlation heatmap of selected protein complexes based on CORUM^119^ were generated using the mean normalized elution profile across replicates (excluding outliers as the most dissimilar fraction to the median).

DSSO cross-linking MS data was analyzed using Thermo Proteome Discoverer (version 2.5.0.400) with the XlinkX module^139,140^. Data was searched against the human canonical proteome (Uniprot Swiss-Prot 2022-05). MS^2^ acquisition strategy was selected with 10ppm precursor mass tolerance, 20ppm FTMS fragment mass tolerance and 0.6Da ITMS fragment mass tolerance. Carbamidomethylation was included as fixed modification; oxidation and N-terminal acetylation were included as variable modifications. A maximum of 3 trypsin miscleavages were allowed and 5 minimum peptide length. FDR threshold was set to 5% and only cross-links with XlinkX score >40 were considered for downstream analysis (**Supplementary Table 2**). Protein domain information of all cross-linked positions was retrieved from UniProt (**Fig. 1f**) and copy numbers were obtained from^24^ (**Extended Data Fig. 2f**). Yeast-two hybrid data was retrieved from ^36^ and IP data from BioPlex 3.0^28^ (**Extended Data Fig. 2j**). The co-fractionation of cross-linked protein pairs in the BN dataset was evaluated using SECAT^141^. Positive and negative interaction networks from CORUM were used as provided. The target network was generated from all the cross-linking interactions for proteins identified in both cross-link and BN. The following parameters were used to ensure the generation of scores for all target protein pairs: peak picking was set to none, monomer threshold factor to 1, minimum abundance ratio to 0, maximum shift to 48 and maximum q-value of 1. SECAT p-values were used for comparison with cross-link data and previously reported interactions from STRINGDB, CORUM and BioPlex 3.0 as described above (**Extended Data Fig. 2l-n**).

DHSO and DSSO cross-linking MS data was analyzed using Scout (version 1.6.2)^38^. Data was searched against the human canonical proteome (Uniprot Swiss-Prot 2022-05) with default parameters, including 10 ppm error on MS1 level and 20 ppm error on MS2 level. Carbamidomethyl (mass 57.02146) and MMTS (mass 45.987721) were included as fixed modification for DSSO and DHSO cross-linked samples, respectively; oxidation and N-terminal acetylation were included as variable modifications. A maximum of 3 trypsin miscleavages and 2 variable modifications were allowed and 6 minimum peptide length. FDR threshold was set to 1% at all levels without separation of cross-link types. Residue Pairs table was used for downstream analysis (**Supplementary Table 2**).

DMTMM cross-linking MS data was analyzed using pLink2 (version 2.3.11)^142^. Data was searched against the human canonical proteome (Uniprot Swiss-Prot 2022-05) with 15ppm precursor mass tolerance and 20ppm fragment mass tolerance. Methylthio(C) was included as fixed modification; oxidation and N-terminal acetylation were included as variable modifications. A maximum of 3 trypsin miscleavages were allowed and 6 minimum peptide length. Filter tolerance was set to 10ppm and separated FDR threshold to 1% at PSM level. Filtered cross-liked sites were used for downstream analysis (**Supplementary Table 2**). DMTMM and DHSO cross-links were mapped to all possible protein interactions defined by DSSO cross-links considering that each DMTMM/DHSO cross-link could match multiple interactions due to shared peptide sequences.

### EndoMAP.v1 network analysis

A protein-protein interaction network was generated from all protein pairs identified by cross-link and BN. The network was initially filtered to remove proteins present in the native molecular weight markers (spiked-in proteins used as reference in BN experiments), EEA1 (overexpressed and used as handle for the endosome affinity purification), UBC (in most cases correspond to a protein modification rather than a member of a protein complex) and keratins (common contaminant). Network characterization and analysis was performed using *igraph* R package (**Extended Data Fig. 3a-c**). Proteins were assigned to subcellular location according to the following annotations: endosomal proteins from our meta-analysis described above (**Supplementary Table 1**), Golgi proteins (as curated on^132^), lysosomal proteins (*bona fide* proteins from^143^, TableS3; *bona fide* and experimentally determined proteins from^144^, TableS2 and TableS12 respectively), mitochondrial proteins (from MitoCarta3.0^145^) and nuclear proteins (based on Uniprot, proteins exclusively designated with nuclear-related terms such as “Nucleus” and “Chromosome”). These annotations and *circlize* R package were used to generate the network chord diagram (**Extended Data Fig. 3d**).

The network centered around endosomal proteins (or “EndoMAP.v1”) was generated by filtering dubious interactions (i.e. nuclear proteins) and including only endosomal proteins (as defined by the meta-analysis) and their direct interactors. Up to 8.5% of the endosomal interactions involved nuclear proteins (**Extended Data Fig. 3d**), which may be considered questionable (therefore were filtered out) and may indicate false connectivity at the PPI level introduced by sample preparation. Second order interactors of endosomal proteins were only included when connected to at least one direct interactor by cross-link and/or two direct interactors by BN (**Extended Data Fig. 3e**). The core component of the network (i.e. biggest module) was visualized using Cytoscape v3.10.1 and protein communities were detected by unsupervised edge-betweenness analysis (**Fig. 2a**). Gene Ontology (GO) enrichment analysis was performed for each community using g:Profiler with the whole proteome as background (**Supplementary Table 2**, including only significant GO Cellular Component, GO Biological Process and CORUM terms with at least two proteins). Path distance analysis between proteins assigned to complexes was based on CORUM and GO:CC (only terms related to protein complexes) (**Fig. 3b, Extended Data Fig. 3f**). Graph rewiring with the same degree distribution (100 permutations) was used as randomized control (**Fig. 3c**). Disease over-representation analysis of the endosomal proteome was performed on endosomal proteins as defined by meta-analysis and as annotated in Gene Ontology (GO:0005768, date 12-2024). Enrichment analysis for the disease gene network (DisGeNET)^146^ was performed as implemented in *DOSE* R package. Enrichment analysis for neurodegenerative disorders included Autism Spectrum Disorders (ASD), epilepsy and severe neurodevelopmental disorder (DD/ID), and schizophrenia was based on ^147^, and was performed using *clusterProfiler* R package with brain expressed genes as background. Path distances analysis between proteins linked to neurodegenerative disorders was based on Diseases 2.0^148^ (2024-02 update), Parkinson’s disease reviewed genes^21^ and Parkinson’s disease GWAS^149,150^ (**Extended Data Fig. 3j-k**).

### AlphaFold Multimer (AF-M), AlphaLink2 and structural modeling

AF-M was run with ColabFold v 1.5.2^9^ on 40GB A100 NVIDIA GPUs for all protein pairs identified by XL-MS and 3-clique combinations within EndoMAP.v1 (with a maximum of 3,600 amino acids in total). AF-M version 3 was used with weights models 1, 2, and 4 with 3 recycles, templates enabled, 1 ensemble, no dropout, and no AMBER relaxation. The Multiple Sequence Alignments (MSAs) supplied to AF-M were generated by the MMSeq270 server with default settings (paired + unpaired sequences). Structure Prediction and Omics informed Classifier (SPOC) and contact sites were calculated as described ^37^. The quality of the predictions was considered acceptable with a SPOC >0.33 for pairwise predictions and at least two interfaces with interface average models >0.5 for timer predictions. AlphaLink2 (github.com/lhatsk/AlphaLink) was performed as described^27^ using intra and interprotein DSSO cross-links. Three predictions for each protein pair were generated with AlphaLink2 by using different seeds. All PDB structures containing protein pairs identified by XL-MS were retrieved by querying the PDB API for X-Ray and Cryo-EM structures with overall resolutions <3.5Å. PDB chains were mapped to their corresponding UniProt identifiers with PDB SIFTS API. Cross-links were mapped onto the AF-M and PDB structures, and cross-linked residues with a maximum Cα-Cα distance of 35Å were considered to match the cross-linker constraints. For AlphaLink2, the maximum Cα-Cα distance considered was 30Å for all cross-linkers, a more stringent threshold since DSSO cross-links were already used to assist the prediction generation. For AF-M and AlphaLink2 predictions, only cross-linked residues with both pLDDTs >70 were considered for distance analysis. Cross-links involving HSP90AA1/B1, which present a large number of cross-links, were excluded from the distance distribution plots in AlphaLink2 predictions (**Extended Data Fig. 4m-o**) to make the analysis more representative of the entire dataset.

The association of mTORC1-Ragulator complex with V-ATPase was modeled using HADDOCK2.4 web server^151^. The cross-links identified between ATP6V1C1-LAMPTOR2 and ATP6V1C1-LAMPTOR4 were used as unambiguous restraints with upper distance limit of 23Å and center of masses restraints enabled. The complete mTORC1-Ragulator complex structure (PDB 7UXH)^66^ was included with selected subunits of V-ATPase due to the limitation in the maximum number of atoms (PDB 6WM2 chains I and J from ATP6V1E1, chain L and M from ATP6V1G1, chain O from ATP6V1C1, chain 8 and 9 from ATP6V0C)^63^. The model with the best score compatible with the expected membrane topology was selected (cluster 5, **Fig. 5f, Extended Data Fig. 11b**). Structure images were generated with PyMOL 2.6.0. All input, parameter, and output files can be obtained at Zenodo (10.5281/zenodo.14679635).

