## Supplementary Figure 1 for "EndoMAP.v1, a Structural Protein Complex Landscape of Human Endosomes"

### Figure 3d

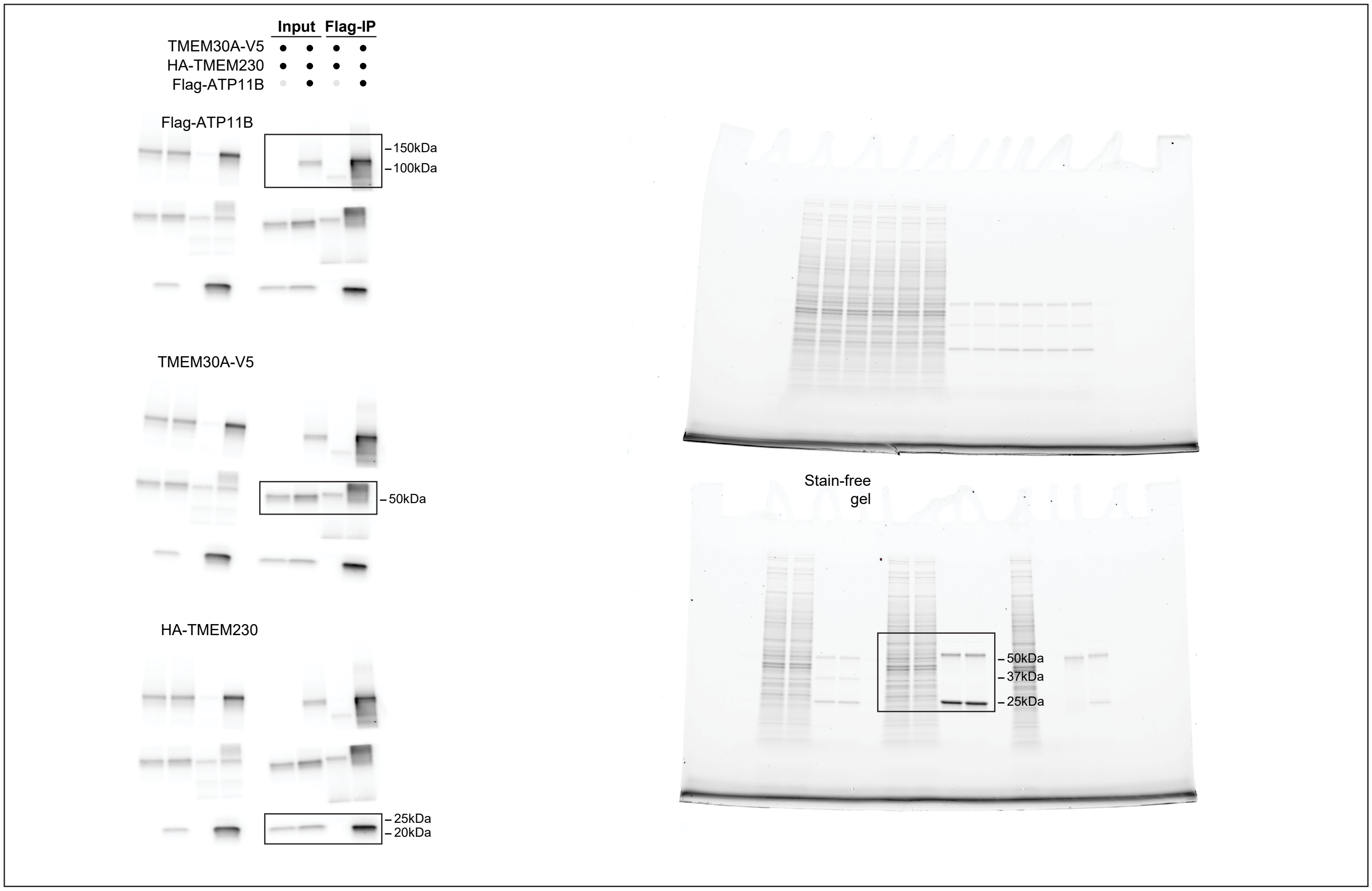


### Figure 3h

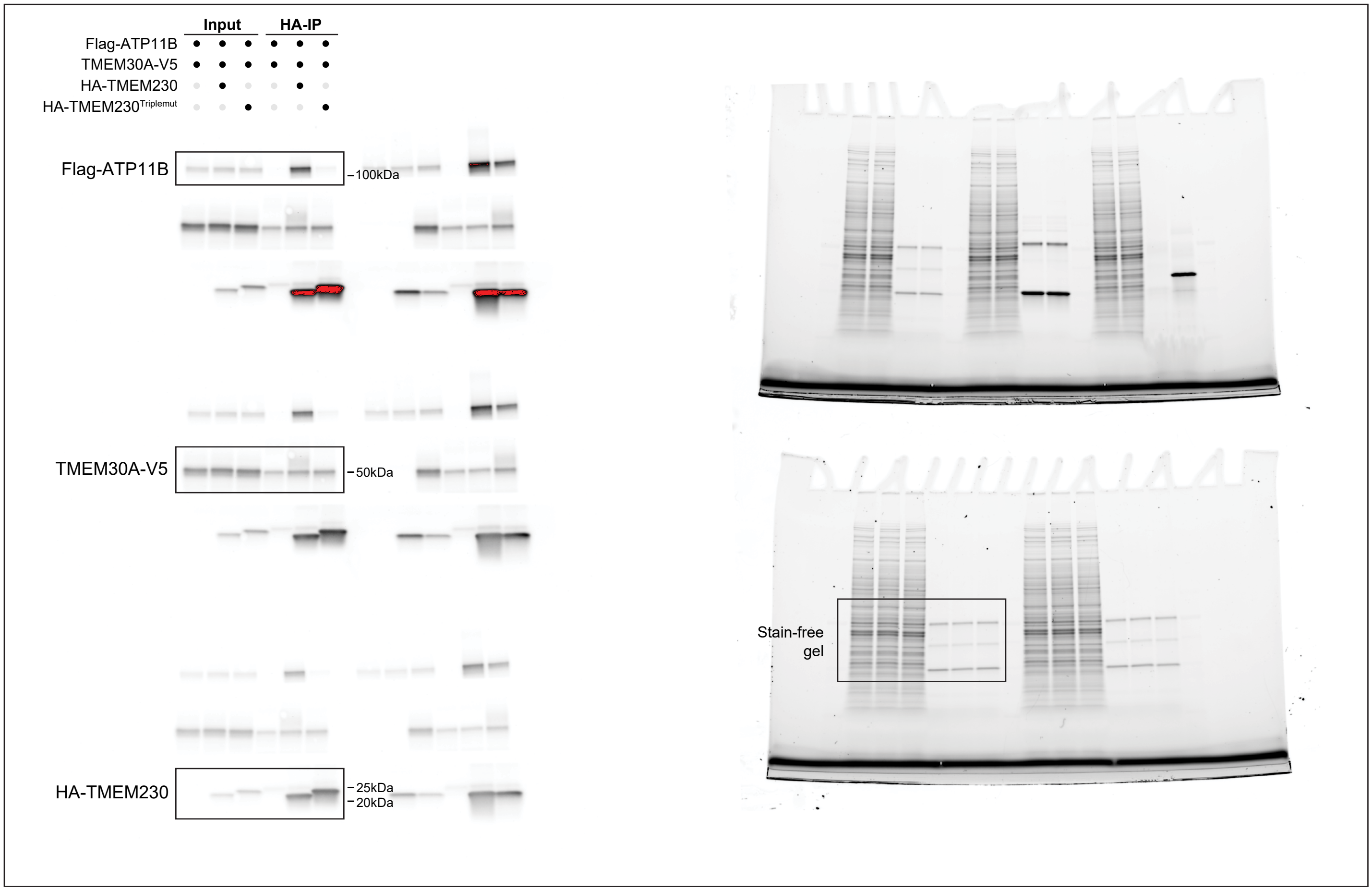

#### Extended Data Figure 5c

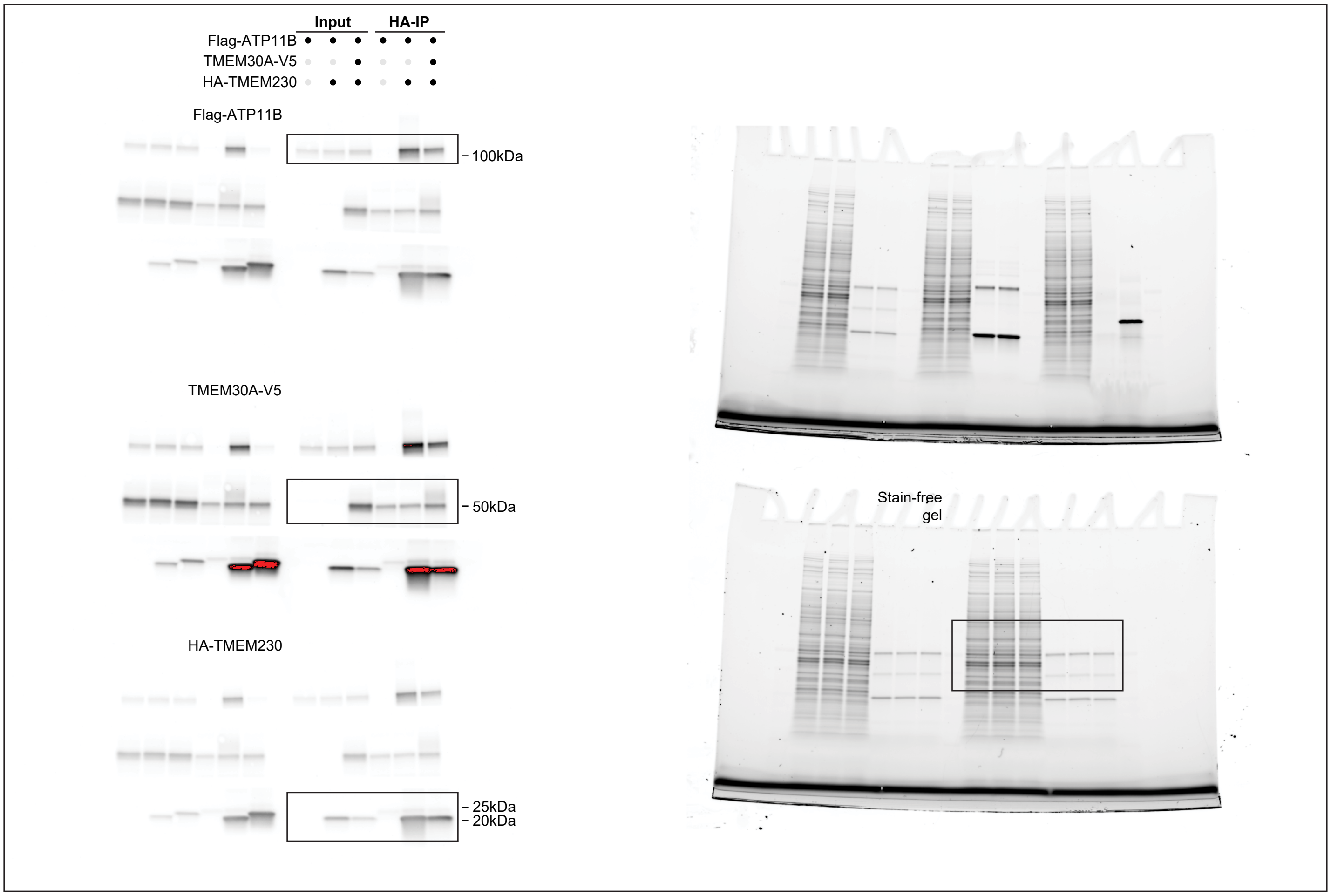


#### Extended Data Figure 5e

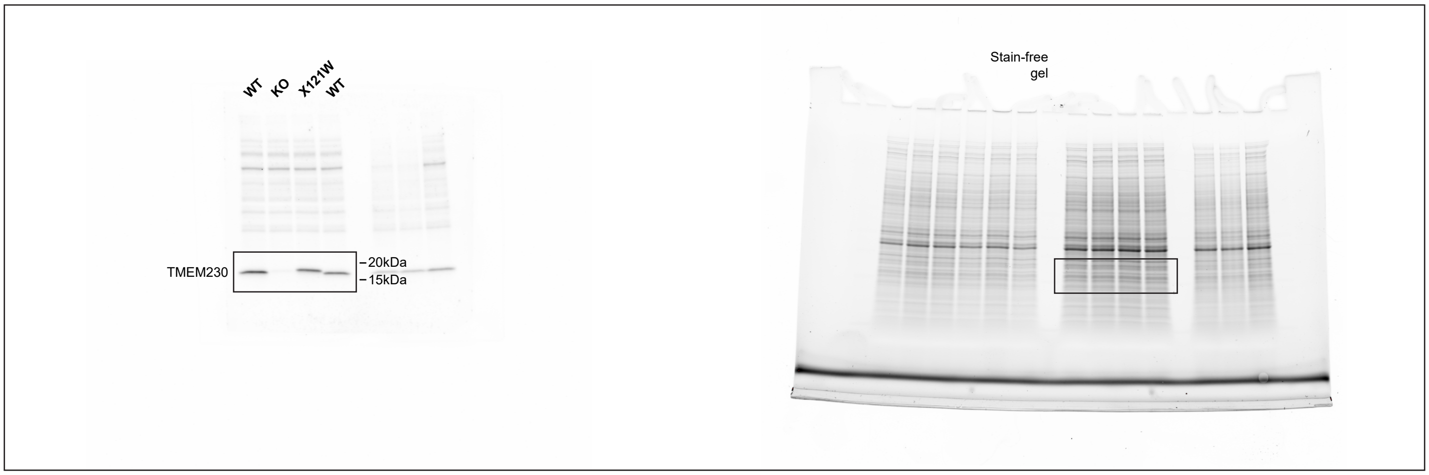

### Extended Data Figure 5h

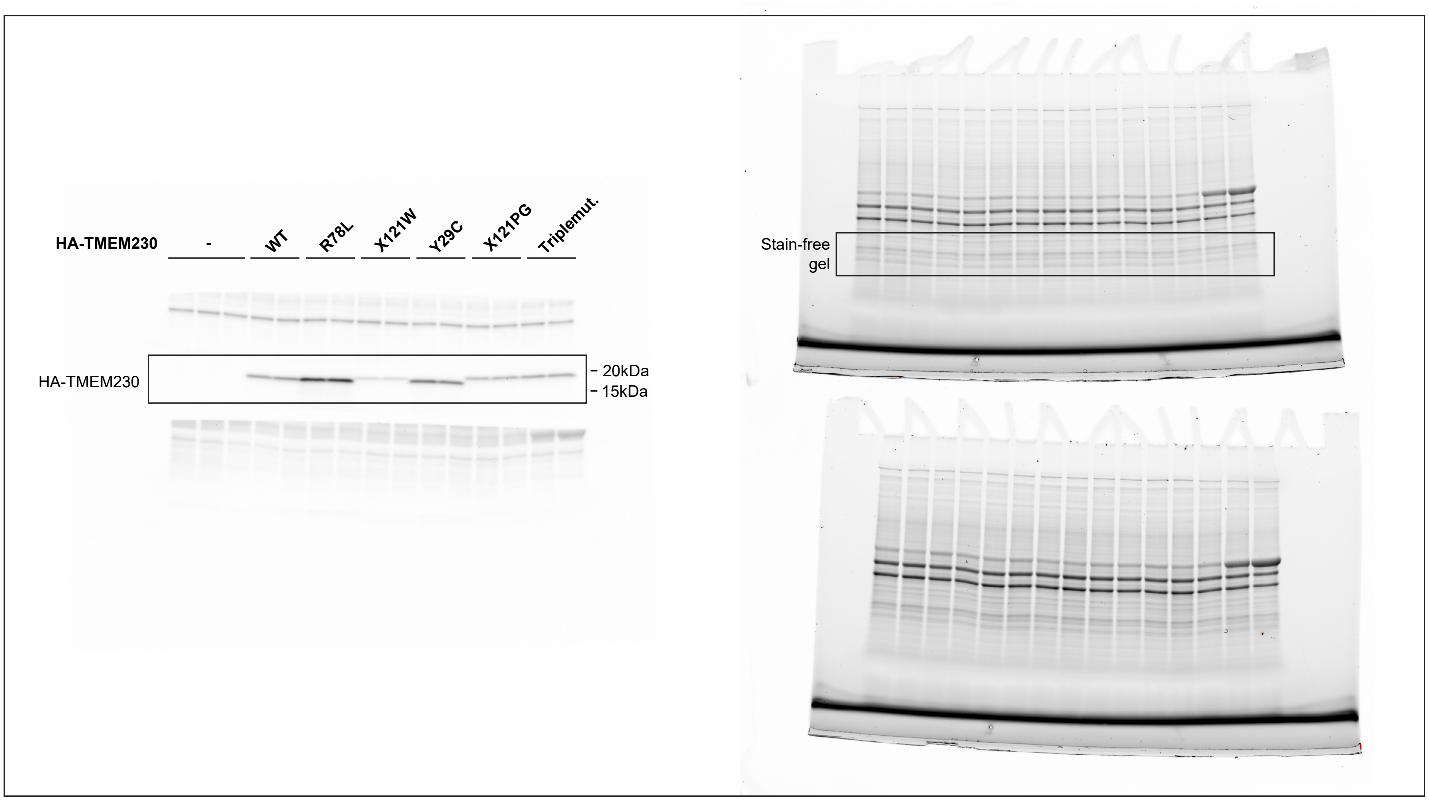

### Extended Data Figure 5m

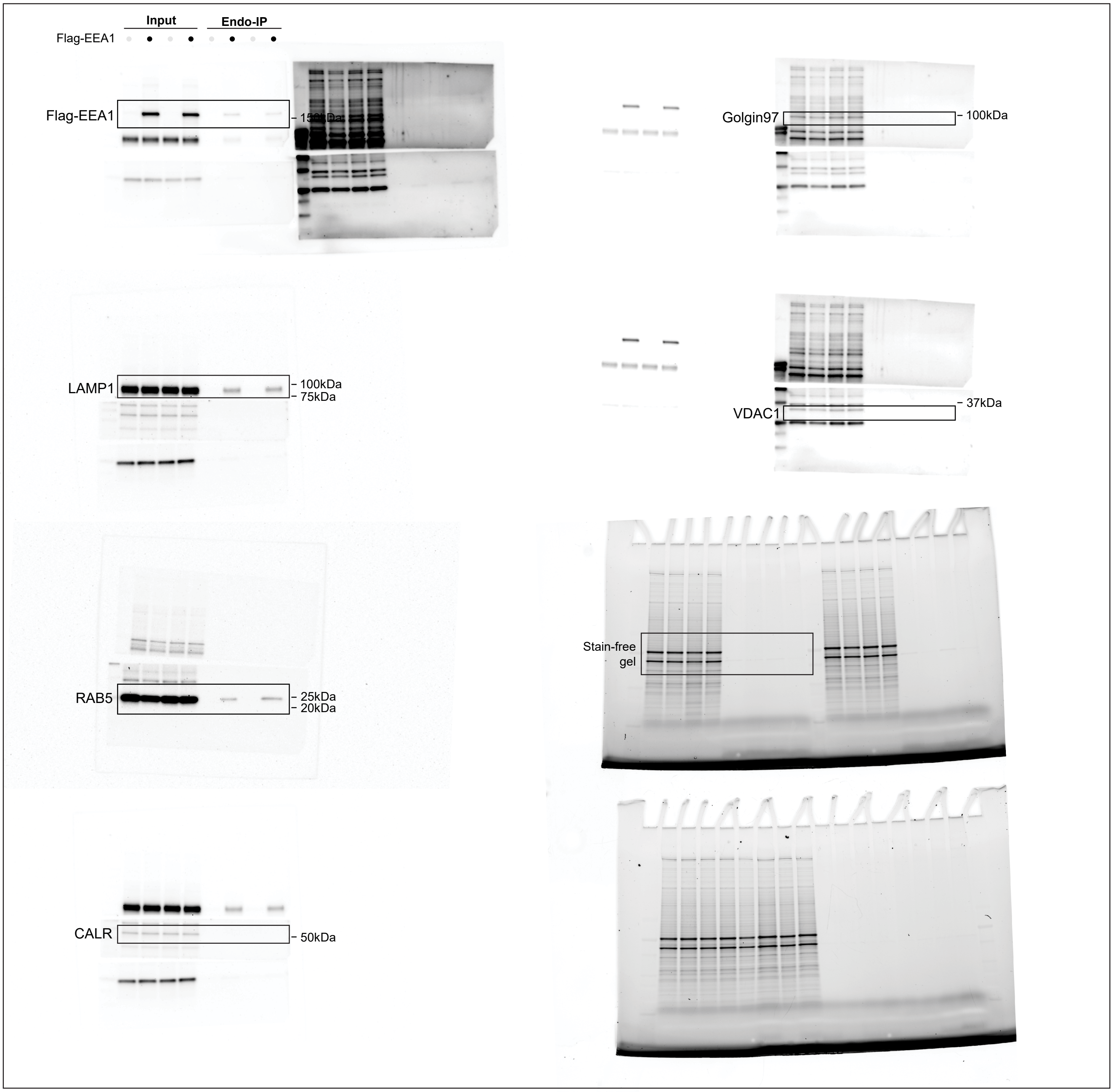

### Extended Data Figure 7g

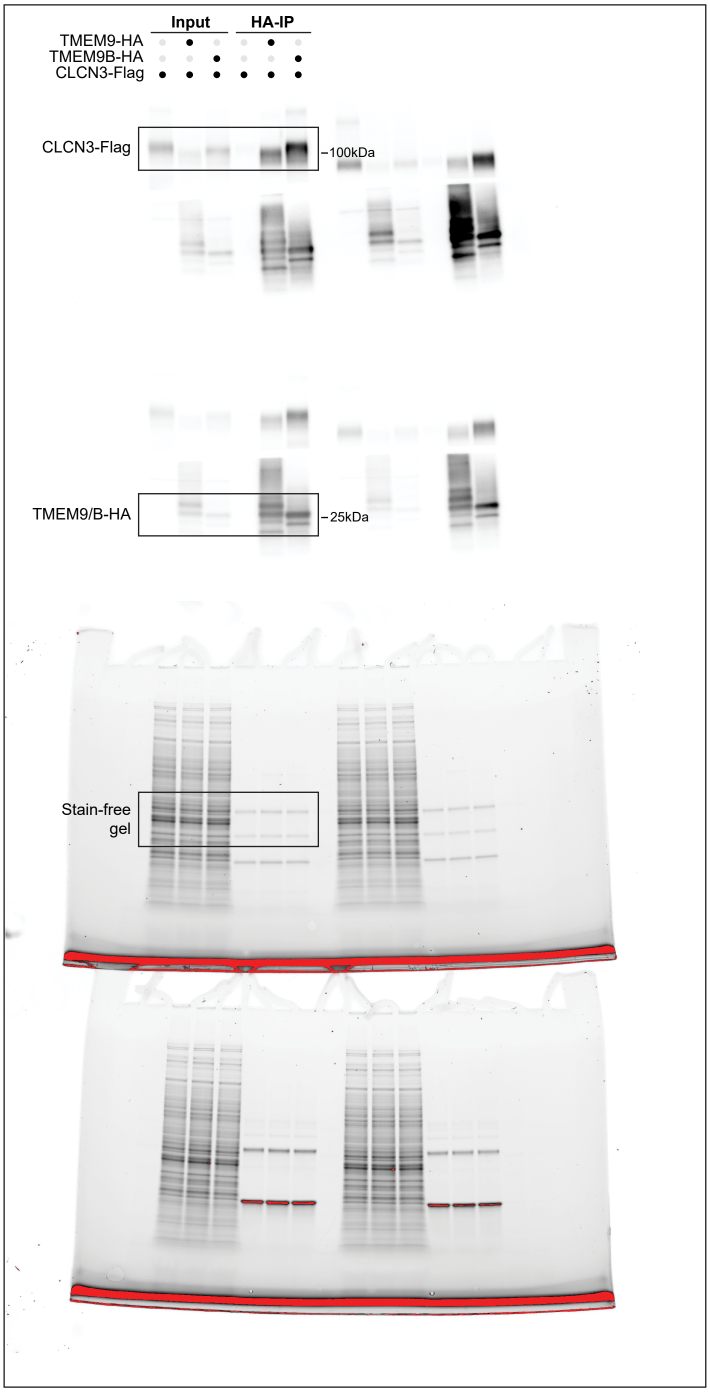


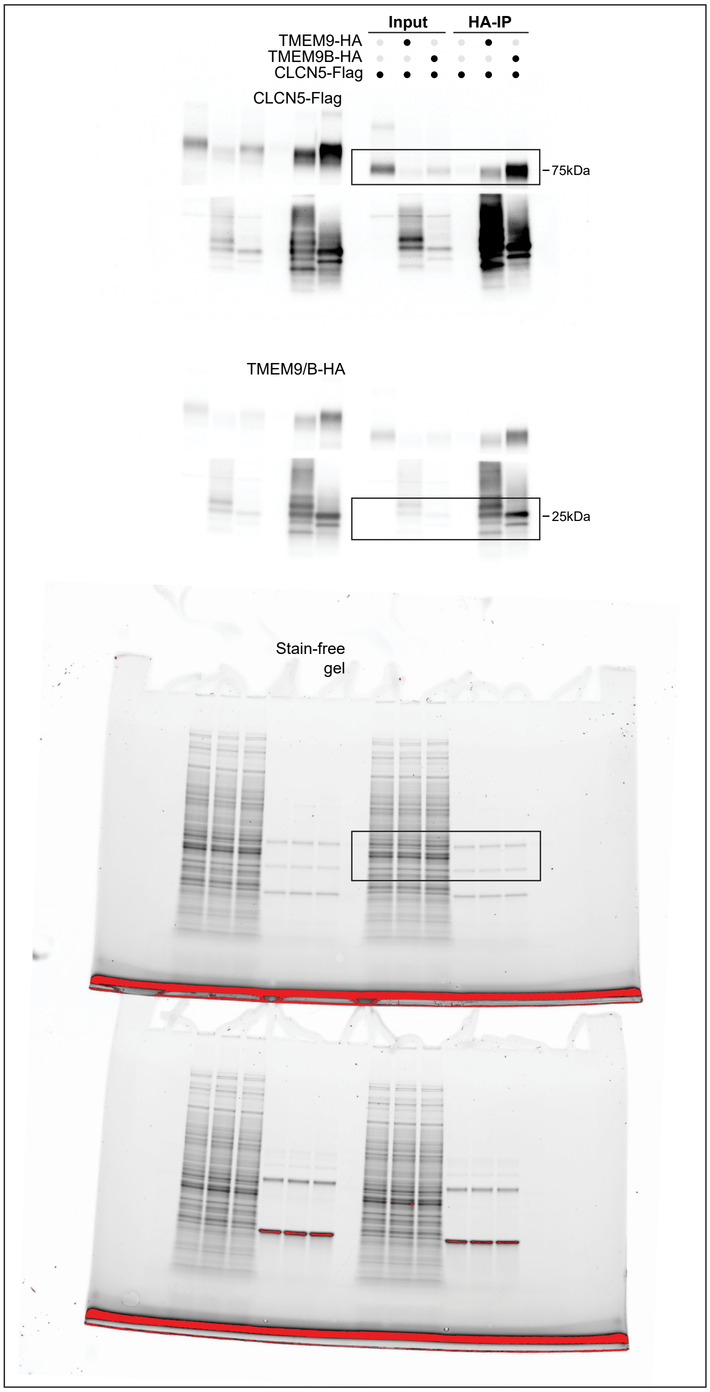

### Extended Data Figure 7g (cont.)

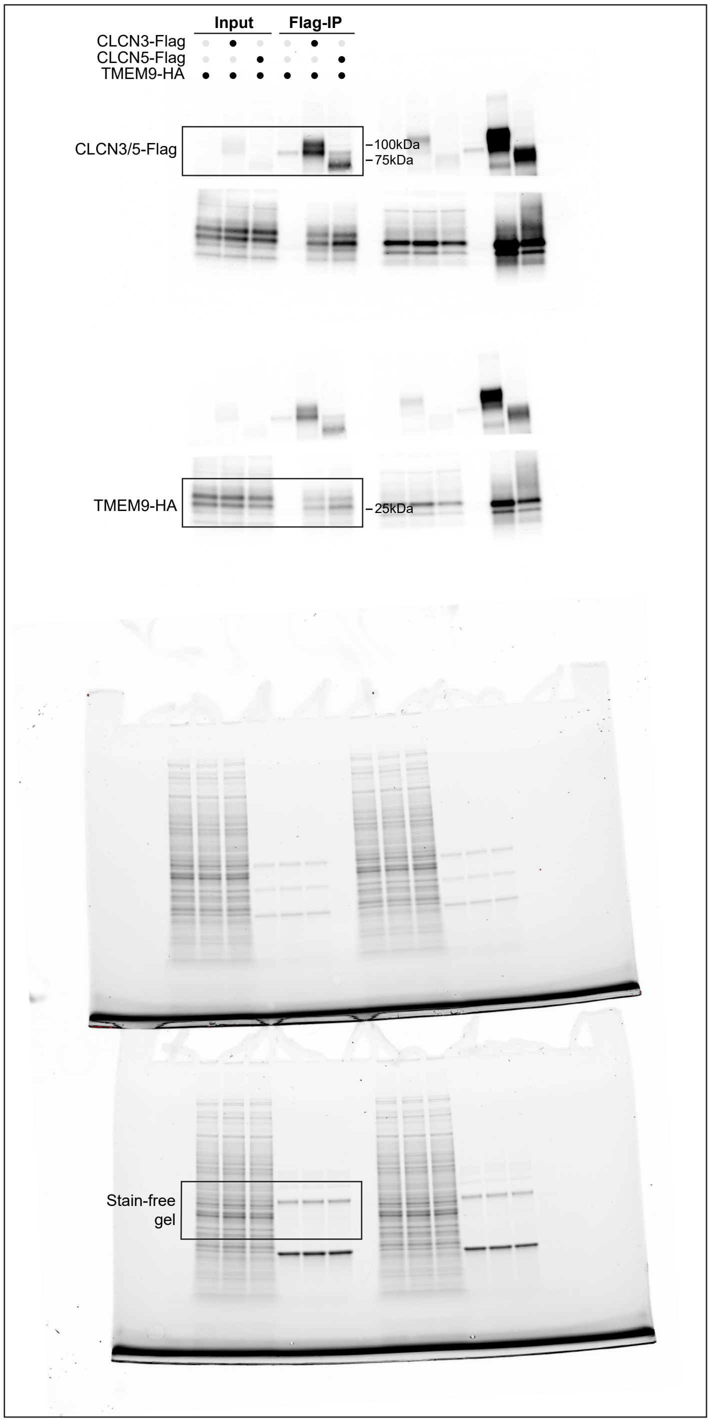


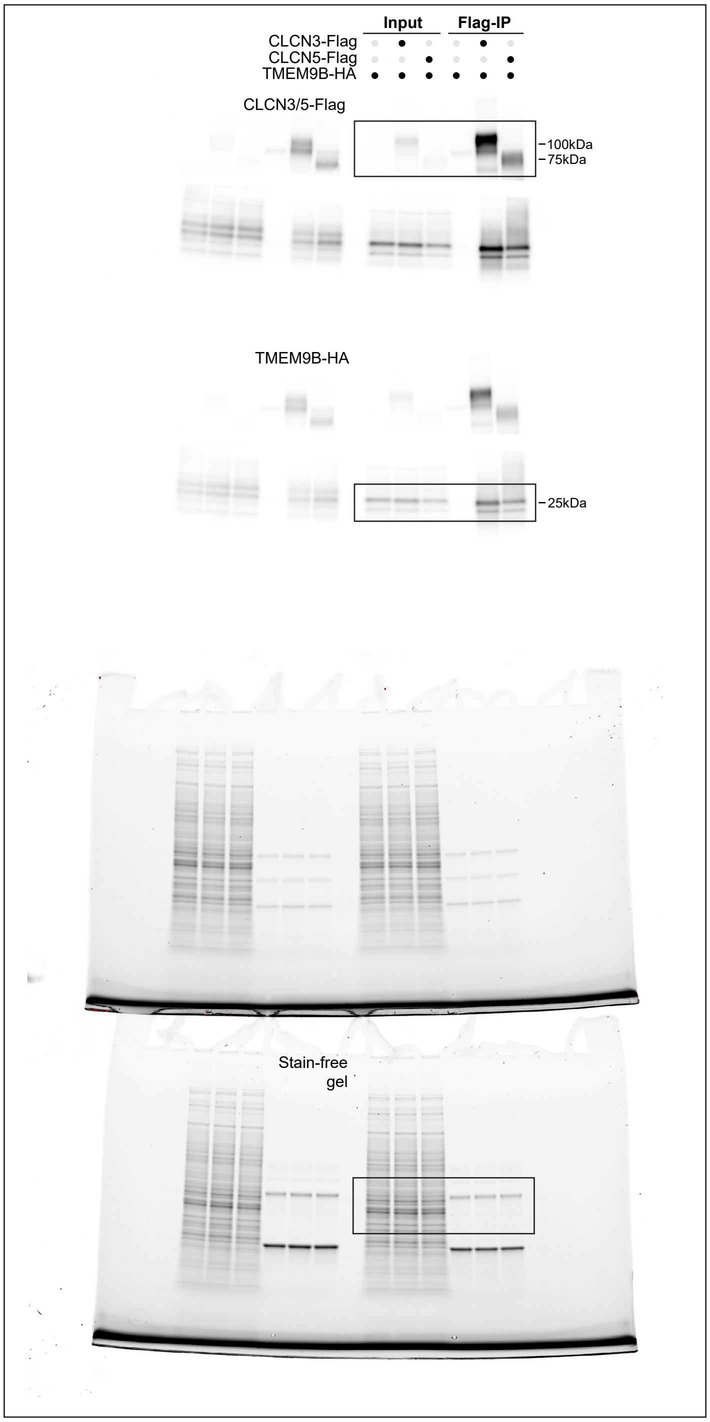

### Extended Data Figure 8c

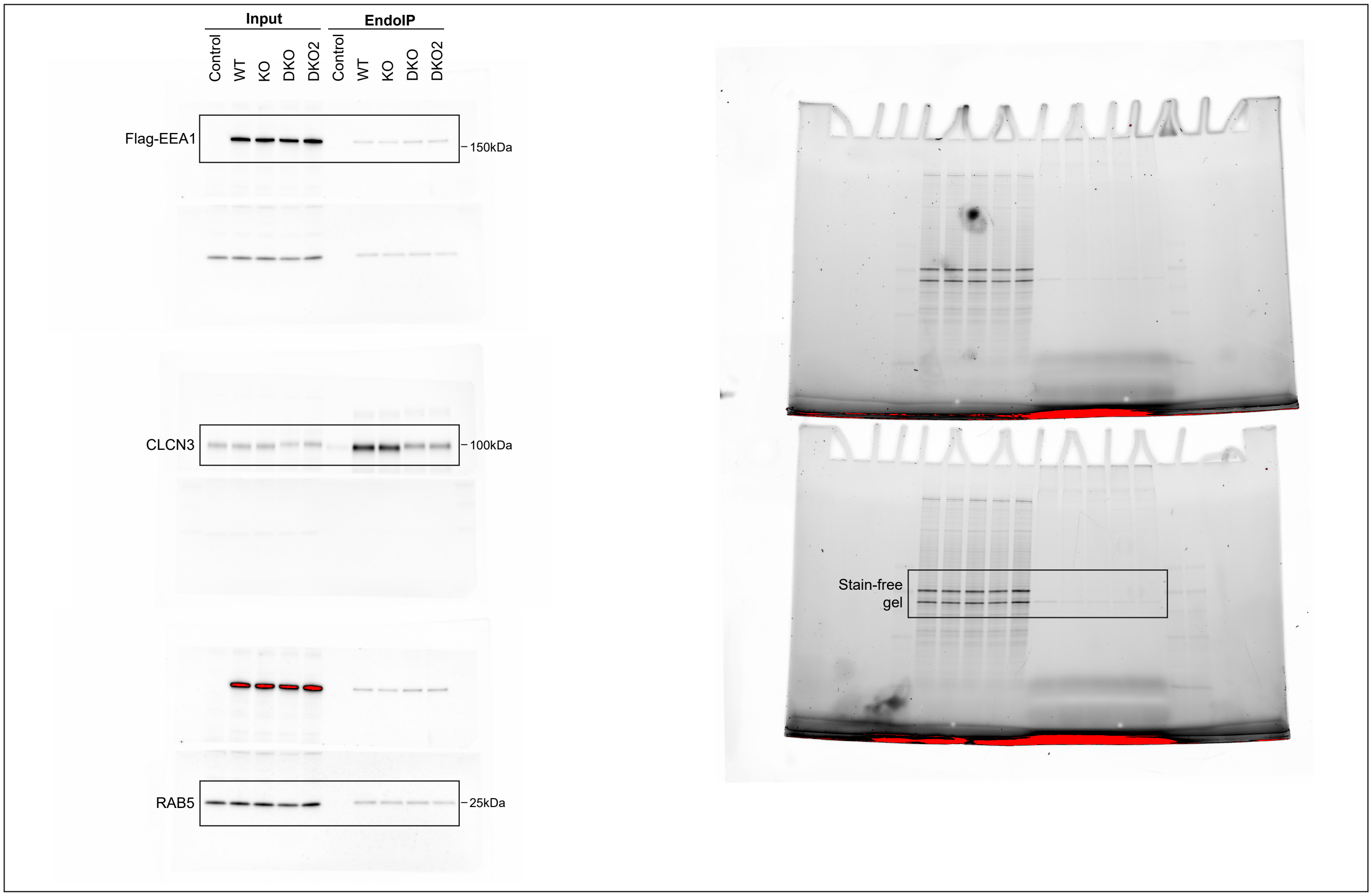
