## Supplementary figures and images for "EndoMAP.v1, a Structural Protein Complex Landscape of Human Endosomes"

### Supplementary video 1

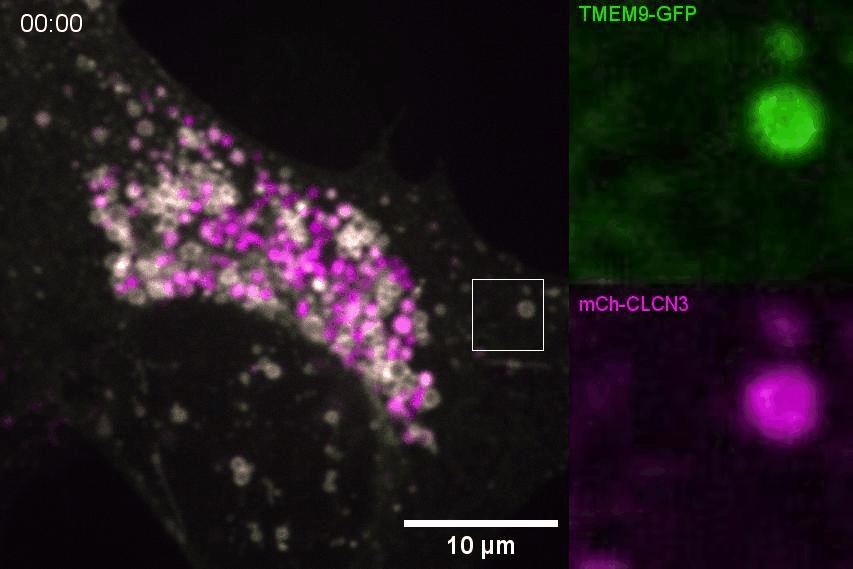
